# Dissecting Regulatory Non-Coding Heart Disease GWAS Loci with High-Resolution 3D Chromatin Interactions Reveals Causal Genes with Pathophysiological Relevance to Heart Failure

**DOI:** 10.1101/2024.10.08.617295

**Authors:** Richard Gill, Daniel R. Lu, Ittai Eres, Jiamiao Lu, Jixin Cui, Zhongsheng Yu, Tracy Yamawaki, Hong Zhou, Baikang Pei, Junedh M. Amrute, Yen-Sin Ang, Songli Wang, Kory J. Lavine, Brandon Ason, Chi-Ming Li, Yi-Hsiang Hsu

## Abstract

Heart failure is caused in part by cardiac remodeling processes that include the death of cardiac myocytes and their replacement by cardiac fibroblasts. We hypothesized that these two cell types may harbor epigenetic contexts in which heart disease-associated non-coding SNPs perturb gene expression relevant to disease. Accordingly, we generated high-resolution Hi-C data layered with chromatin accessibility and transcriptomic information to annotate and link putative distal regulatory elements in heart disease-associated loci to gene promoters. Our analysis identified several target genes with established roles in cardiac fibrosis and/or heart disease (*GJA1*, *TBC1D32, CXCL12*, *IL6R*, and *FURIN*). Perturb-seq in cardiac fibroblasts to knock out putative regulatory elements confirmed regulatory relationships involving *GJA1*, *CXCL12*, and *FURIN*, as gene editing led to changes in transcriptomic signatures associated with fibroblasts in heart failure. Our results demonstrate how integrative multi-omic approaches can delineate pathophysiologically relevant regulatory circuits that connect protein-coding genes to non-coding genetic variants associated with disease.

## Background

Heart failure is characterized by systolic and/or diastolic dysfunction and represents a rapidly growing public health problem and unmet medical need in the US. While the population prevalence of heart failure was 2.4% in 2012, this figure will increase more than ten-fold to 25% by 2030 as a surplus of Americans reach age 65 [1]. In parallel, the medical costs of heart failure are projected to rise from $20.9 billion in 2012 to $53.1 billion in 2030 [2]. Heart failure is a complex syndrome secondary to functional or structural deficits that impair ventricular filling or blood ejection, including ischemic and valvular heart disease, cardiomyopathy, congenital heart disease, and pericardial disease. The diminished contractility and reduced cardiac output due to these initiating insults and cardiac myocyte death lead to maladaptive compensatory cardiac remodeling (hypertrophy or dilation concurrent with cardiac fibrosis); this stiffens the myocardium and further impairs cardiac filling and myocardial contraction [3], ultimately leading to cardiac decompensation and heart failure.

Cardiac fibroblasts are highly relevant to the pathophysiology of heart failure and its pathogenesis. These mesenchymal cells are involved in remodeling-associated hypertrophy and dilation, as well as post-infarction replacement of dead cardiomyocytes with a collagen-rich (fibrotic) scar to maintain the heart’s structural integrity. However, if disease-activated fibroblasts become hyperactive (myofibroblasts), their excessive production and secretion of extracellular matrix (ECM) can result in the appearance of patchy or diffuse excess fibrotic tissue, and, consequently, increased myocardial wall stiffness. Fibroblast proliferation and phenoconversion to myofibroblasts are both initiated by Transforming Growth Factor B (TGF-β) and its protease regulators such as FURIN [4] and MMP family proteins [5]. This profibrotic signal is mediated by transcription factors such as SMAD3 and SMAD4, as well as Connective Tissue Growth Factor (CCN2/CTGF) [6], and is further modulated by Kruppel-Like Factor family protein 15 (KLF15) [7]. Other important factors in cardiac fibrosis include Angiotensin II (Ang II) [8], Endothelin 1 (EDN1) [9–11], Wnt-β-Catenin signaling [9, 12], and the SDF1α (CXCL12)/CXCR4/CXCR7 axis [13].

Heart failure and its antecedent diseases have a strong genetic component and have accordingly been intensively investigated in genome-wide association studies (GWAS), which have yielded numerous loci with robust statistical associations [14–28]. However, across cardiovascular and many other unrelated diseases, most associated SNPs reside within non-coding (intronic or intergenic) parts of the genome. While most (∼80%) of these loci are themselves enriched in putative regulatory elements [29, 30], their target genes are frequently not known. One approach to assign target genes to GWAS variants is expression quantitative trait locus (eQTL) mapping, which helps determine which genes’ expression may be impacted by allelic dosage of a given SNP. However, the utility of eQTL overlap alone for understanding GWAS is limited: most existing eQTL signals have been generated from bulk expression data derived from whole tissues comprised of heterogeneous mixtures of cells, have been discovered in cohorts much smaller than typical GWAS sample sizes, and as a result, have been subject to different linkage disequilibrium patterns and potential confounders [31][32] Therefore, even when co-localized with eQTLs, non-coding GWAS signals are often mapped, and mis-attributed, to the nearest gene(s), and their direct relevance to disease is not clear.

Studies in numerous disease contexts suggest that causal SNPs’ regulatory targets may often be thousands to millions of base pairs away, yet physically proximal due to individual DNA loops [33–37], as well as larger-scale regulatory structures termed topologically associating domains (TADs) [38–40]. Recent advances in all-to-all chromosome conformation capture techniques (i.e. Hi-C) coupled with next-generation sequencing now permit identification of significant DNA-DNA interactions and TADs genome-wide. However, the majority of published Hi-C data are low-resolution [41] or derived from cell lines not representative of disease contexts [42]. Capture Hi-C partially allays resolution issues by focusing on identifying loops involving promoter-centric regions at high-resolution [43], but uses baits that may introduce capture efficiency biases [44]. Two recent studies have applied this approach in cardiomyocytes derived from induced pluripotent stem cells (iPSCs) [45, 46], but, in addition to problems associated with biased interaction sampling, iPSC-derived cardiomyocytes display immature phenotypes [47], and therefore may not accurately represent adult-onset cardiovascular disease contexts.

To address these gaps and determine how non-coding GWAS loci contribute to heart failure etiology, we integrated high-resolution 3D chromatin interaction and epigenetic data in primary human ventricular myocytes and fibroblasts. Using this multi-layered omics dataset, we systematically constructed genome-wide gene regulatory circuits between heart disease GWAS SNPs and their target genes, irrespective of genetic distance. We generated Hi-C data at 2-kb resolution to capture genome-wide pairwise physical interactions, ATAC-seq data to assay regions of open chromatin and any associated transcription factor footprints (cell-specific motif-predicted TF binding sites accessible to transcriptional machinery), as well as bulk RNA-seq data to identify expressed TFs and target genes. Significant loops from Hi-C data were intersected with pairs of known gene promoters and heart disease loci prioritized with ATAC-seq and RNA-seq data to identify regulatory relationships, and these putative enhancer-promoter pairs were further compared with GTEx eQTLs. The resultant regulatory circuits’ target genes were enriched for established cardiac remodeling and heart disease associated genes and may include novel targets for therapeutic intervention. Finally, we validated a subset of high priority SNP-gene pairs using CRISPR-Cas9 deletion of the regulatory regions followed by single cell RNA-seq based quantification of the putative target genes and characterization of transcriptomic consequences following perturbation of putative enhancer regions, confirming their regulatory role and the value of multi-omic data integration for target gene identification and prioritization.

## Materials and Methods

### Cells

#### Primary Human Cardiac Myocytes and Fibroblasts

We generated Hi-C, ATAC-seq, and RNA-seq data from primary cells to model heart failure-relevant cellular contexts. Specifically, we cultured 50 million primary Human Cardiac Myocytes (HCM) cells as well as Normal Human Cardiac Fibroblasts – Ventricular (NHCFV) primary cells, which are commercially available from PromoCell and Lonza respectively.

#### Immortalized Human Cardiac Fibroblasts

We performed gene editing experiments using Immortalized Human Cardiac Fibroblasts generated from adult heart ventricle primary cells (iHCF), which are commercially available from Applied Biological Materials Inc. These experiments were performed in iHCF as opposed to primary HCM or NHCFV because primary cells have limited life span and are less tolerant of manipulations such as electroporation and gene editing, compared to cell lines. To account for this difference and enable comparisons of peak calls and gene expression levels between immortalized and primary cells, we also generated ATAC-seq and RNA-seq data from this cell line.

### Hi-C Library Construction and Data Processing

We contracted Arima Genomics to construct in-situ Hi-C libraries for 10 million HCM and NHCFV cells based on the protocol from Rao et al. [42]. For HCM cells, they used a mixture of restriction enzymes that cut at ^GATC and G^ANTC, while an enzyme cutting at ^GATC was used for NHCFV cells. We then sequenced over 2 billion paired-end reads (2.7E9 for HCM and 2.4E9 for NHCFV) using an Illumina HiSeq 4000. Reads from both biological replicates were processed together for each cell type (aligned to hg38 and read- and fragment-filtered to identify valid interaction pairs) using both a) the HiC-Pro pipeline version 2.8.1 [48] and b) the Juicer pipeline version 1.5.6 [49] on the Juicer Amazon Machine Image (AMI) on Amazon Web Services (AWS). We called genome-wide loops representing predicted genome-wide pairwise physical interactions between 2-kb regions of the genome using GOTHiC version 1.12.0 [50] and HiCCUPS included with Juicer 1.5.6 [49].

### Hi-C Loop & TAD Calling

#### GOTHiC

The BAM file from HiC-Pro containing valid interaction pairs (filtered for invalid interaction products), was converted to GOTHiC format using hicup2gothic, and split by chromosome. We ran GOTHiC in base R 3.4.4 on AWS to call significant interactions from each chromosome-specific file. Statistical significance was set at FDR <0.05 and contact counts >10 based on previous studies [51, 52].

#### HiCCUPS

Using the .hic file generated by Juicer, we ran HiCCUPS on an AWS GPU instance to identify loops at multiple resolutions (2-kb, 5-kb, 10-kb, and 25-kb) by calling peaks enriched compared to four local backgrounds: (i) the donut shape around a pixel, (ii) in the lower-left quadrant of the donut neighborhood, as well as the (iii) vertical and (iv) horizontal neighborhoods [42]. At each resolution we used the following arguments for the peak width (*p*), window (*i*), FDR threshold (*f*), and centroid distance (*d*) parameters suggested by the authors: 2-kb (*p*=10, *i*=20, *f*=0.1, and *d*=20,000), 5-kb (*p*=4, *i*=7, *f*=0.1, and *d*=20,000), 10-kb (*p*=2, *i*=5, *f*=0.1, and *d*=20,000), and 25-kb (*p*=1, *i*=3, *f*=0.1, and *d*=50,000). Vanilla coverage was used for normalization, as the Knight-Ruiz method did not converge for some chromosomes at 2-kb and 5-kb.

#### TAD Calling

We used the .hic files generated by Juicer to infer TADs with the arrowhead algorithm at resolutions of 5, 10, 25, 50, 100, and 500 kb, using Vanilla coverage normalization. To ensure robust results, we also converted our .hic files to sparse format at these same resolutions, and subsequently applied GRiNCH [53], with a neighborhood size of 5 bins and default parameters, to infer TADs via graph regularized non-negative matrix factorization and clustering. TADtool [54] was also used to infer TADs using both directionality index and insulation score. We discovered TADs using a window size 5X the resolution, and a cutoff value per-chromosome representing the top 5% of the observed or absolute values for insulation score and directionality index, respectively. The TAD intervals identified in each cell type were then overlapped with the start and end of our identified SNP-gene pairs using bedtools [55] version 2.22.1. We utilized bedtools intersect with parameter -f 1.0 to assess how many SNP-gene pair bin intervals fell entirely within the same TAD.

### Bulk ATAC-seq Data Processing, Peak Calling, and Transcription Factor Footprinting

We constructed two ATAC-seq libraries from 1 million HCM cells, a single library from 1 million NHCFV cells, and 100,000 iHCF cells using the protocol from Buenrostro et al. [56], Libraries were sequenced at >150 million paired-end reads per sample on either an Illumina HiSeq 4000 or NextSeq 500, resulting in >50 million de-duplicated retained reads per sample. We used the minimum number of iHCF cells required for ATAC-seq (10x fewer than the primary cells) due to the difficulty of culturing these cells, and to save material for other experiments. We processed raw reads and called open chromatin regions using the ENCODE ATAC-seq pipeline [57], setting hg38 as the reference genome. We retained putative open chromatin regions from naïve overlap peaks called from two pseudo-replicates for downstream analyses.

To find cell-specific transcription factor binding sites (TFBSs) in ATAC-seq peaks, we performed a nucleotide-level computational search for transcription factor (TF) footprints using HINT-ATAC from Regulatory Genomics Toolkit (RGT) release 0.11.4 [58–61]. The output included motif-predicted binding sites of known TFs from the HOCOMOCO [62] and JASPAR [63, 64] databases, as well as bit-scores of motif matching [59, 65]. We also analyzed differentially footprinted regions to compare predicted TF binding between cell types.

### RNA Sequencing and Transcript Quantification

Global transcript expression in HCM, NHCFV and iHCF cells was assessed by RNA sequencing (RNA-seq). RNA-seq was performed on a cDNA library prepared from total RNA (2 μg; RIN score > 9.5) of 100,000 cells isolated using Mirvana miRNA RNA isolation kits (Ambion, Grand Island, NY) with on-column DNase treatment. Total RNA quality and concentration was determined using the Bioanalyzer (Agilent, Santa Clara, CA) and Nanodrop (ThermoScientific, Wilmington, DE). cDNA was prepared using a modified protocol based on the Illumina Truseq RNA Sample Preparation Kit (Illumina, San Diego, CA) and the published methods for strand-specific RNA-Seq [66, 67]. After size selection of libraries (Pippen Prep; SAGE Biosciences, Beverly, MA), dUTP-containing cDNA strands were destroyed by digestion of USER enzymes (New England Biolabs, Ipswich, MA) followed by PCR enrichment for introduction of strand specificity. The enriched cDNA libraries were analyzed in Agilent Bioanalyzer and quantified by Quant-iTTM Pico-Green assays (Life Technologies). RNA sequencing reads (Illumina HiSeq platform, 75 bp paired-end sequencing) were aligned to hg38 and Fragments per Kilobase per Million sequenced, Quantile normalized (FPKQ) values were determined using Array Suite software (Omicsoft, Cary, NC) and in-house software. Genes with FPKQ values >0 were considered expressed.

### Genome Annotations

#### NHGRI SNPs

We queried the NHGRI-EBI GWAS Catalog [68] using the search terms “heart disease,” “obesity,” and “pulmonary hypertension” on September 5, 2019, and also included a heart failure GWAS preprint that is now published [27]. For each SNP, we noted the proximal gene(s) based on the annotations “Mapped Gene” (the gene(s) mapping to intragenic SNPs, or genes up-and down-stream of intergenic SNPs) and “Reported Gene” (gene(s) reported by the study author, typically based on proximity to the reported SNP). We then used LDproxy from LDlink release 3.7.2 [69] to identify all proxy SNPs (r^2^>0.8 in the European populations).

#### FANTOM5 TSSs

To exhaustively identify all human transcription start sites (TSSs), we downloaded Biomart annotated hg38 CAGE Peaks from across all human cell types and tissues assayed by the FANTOM5 consortium [70], and intersected these TSSs with gene symbols from HGNC [71]. In total we obtained 96,254 unique TSSs mapping to 20,224 unique genes.

#### GTEx eQTL Data

We downloaded single-tissue cis-eQTLs computed from 44 human tissues in GTEx release v8 [72]. We first annotated eQTL SNPs with LD-based loci to cluster them into locus-gene pairs, which we then used to annotate our Hi-C-derived SNP-gene pairs in the same loci as having eQTL support.

### Data Integration and Analysis

Our analytical approach aimed to identify and use potentially functional SNPs underlying disease GWAS signals, layered with functional genomic information (Hi-C, ATAC-seq, and RNA-seq), to trace regulatory circuits between putative enhancers and their target genes.

i. Due to linkage disequilibrium (LD), GWAS loci typically contain multiple significant disease-associated SNPs indistinguishable by effect-size and *p*-value, and it is unclear which of these SNPs (or proxy SNPs not genotyped in the original GWAS) underlie the association signals. We therefore studied all reported GWAS SNPs, as well as their proxies (r^2^>0.8 in the CEU population). To prioritize potentially functional SNPs based on overlap with putative enhancers, we layered accessible chromatin regions and noted footprints containing motif-predicted transcription factor (TF) binding sites, especially those of expressed TFs.
ii. We used 2-kb resolution Hi-C data to identify significant pairwise interactions between the SNPs in predicted enhancers from (i) and accessible transcription start sites (TSSs). We hypothesized that SNPs disrupting the function of TSS-interacting enhancers may underlie their significant disease associations.
iii. SNPs in the SNP-TSS pairs from (ii) that are (or are in LD with) significant GTEx eQTLs of the target gene have additional evidence to suggest that the SNP may affect expression of the TSS’ target gene. However, GTEx eQTLs were generated from whole-tissue mixtures of cells and may not always be concordant with SNP-TSS pairs identified from the above analyses of a homogeneous primary cell population.

This approach identifies putative enhancers whose perturbation by disease-associated common genetic variations may affect regulation of genes involved in the etiology of complex diseases. Further, this method distinguishes potential causal SNPs from their proxies at disease-associated loci.

#### Data Visualization

We visualized physical interactions between accessible SNPs and TSSs using Juicebox version 2.15 [73] and Hi-Glass version 2.5.7 [74]. We plotted Hi-C and GWAS data using PlotGardener [75]. Hi-C heatmaps depict observed/expected interaction counts, to accentuate distal interactions and reduce noise from proximal interactions along the x-axis. Since small-scale TADs (<50-kb) appeared redundant with loops, we only plotted larger TADs (50-, 100-, and 500-kb) to improve the clarity of our figures.

### Experimental Validation of High-Priority SNP-Gene Pairs

#### Generation of dual-flanking guide RNA libraries for Perturb-seq

We used the Perturb-seq gRNA-mediated KO strategy in the ventricular cardiac fibroblast cell line (iHCF) to introduce deletions at a subset of ATAC-seq regions prioritized for their overlap with heart disease-related TF footprints and interactions with accessible TSSs of heart disease-related genes. To improve deletion efficiency, we introduced two or more single-stranded guide RNAs (gRNAs) with homology to sequences proximal to the regions 5’ (upstream) and 3’ (downstream) of putative enhancers ranging from 746 to 2,863 base pairs. Valid gRNAs were predicted using the Sigma Design Tool [76] and ChopChop algorithm [77]. The gRNAs with the (1) highest predicted efficiency and specificity, (2) lowest predicted self-complementarity and promiscuity (no sites in genome that are <3 nucleotide mismatch to gRNA), and (3) target sequences most proximal to the transcription factor binding site (0 to 580 bp) were selected for cloning into the LV15 vector (Millipore Sigma). This vector expresses gRNAs under control of the human U6 promoter and gRNA scaffolds contain Capture Sequence 2 in the stem loop, which enables hybridization onto 10X Genomics 3’ gel beads and detection via Perturb-seq. OffSpotter [78] and cas-offinder [79, 80] were used to check gRNA sequences for off-target activity.

The gRNAs were designed to flank each predicted transcription factor binding site such that guides flanking the 5’-end were inserted into an LV15 vector backbone (Millipore Sigma) containing a Puromycin-resistance-BFP (blue fluorescent protein) selection marker, and guides flanking the 3’-end were inserted into an LV15 vector backbone containing a Zeocin-resistance-BFP selection marker. In total, four gRNAs were utilized for each enhancer locus. gRNAs against the exon regions of ACTA2 were also designed as a positive control for targeted deletion, and two non-targeting gRNAs were selected from a published library [81] as a negative control.

#### Lentivirus production and quality control

Cloned plasmids containing gRNA protospacer sequences were packaged into lentivirus at a minimum formulation of 10^7^ viral particles per mL in 200 uL (Millipore Sigma). Plasmid quality and yield were initially verified by plasmid restriction digest and p24 antigen ELISA titer. A functional titer was also calculated to determine lentivirus amounts for iHCF transduction.

#### Guide RNA library titration for Perturb-seq

1 mL of 2×10^5^ cells/mL iHCF (ABM Cat# T0446) cell suspension prepared with FibroGRO^TM^-LS complete media (Millipore Sigma Cat # SCMF002) + 8 µg/mL polybrene (Millipore Sigma Cat # TR-1003-G) was placed into each well of a 12-well plate. Lentivirus gRNA library was thawed at room temperature and mixed by gently tapping the tube several times with fingers. 100 µl of 10-fold dilution of the lentivirus gRNA library was prepared with FibroGRO^TM^-LS complete media+ 8 µg/mL polybrene. 50 µl of 10-fold diluted lentivirus was then used to prepare 50 µl of additional 2-fold serial dilutions ranging from 1/2 to 1/1024. 25 µl of media was added to one well as negative control. 25 µl of 10-fold or additional 2-fold diluted lentivirus was added to the rest of the wells and mixed thoroughly by pipetting up and down a few times. The plate was then incubated at 37°C for 18-20 h. The media containing virus was then removed and replaced with 1 mL of FibroGRO^TM^-LS complete media. The plate was then cultured for an additional 2 days. On day 3 post lentivirus transduction, the media from each well was removed. 200 µl of trypsin was added to each well to detach the cells for 3 min at 37°C. 800 µl of complete media was added to each well to neutralize trypsin. The cells were then transferred to 15 mL conical tubes for centrifugation at 300xg for 5 min. Each cell pellet was resuspended in 500 µl BD Cytofix^TM^ Fixation Buffer (DB Biosciences Cat # 554655) for 10 min at room temperature. The BFP expression of cells in each well was analyzed by FACS. We used the wells that had between 5% and 20% of cells expressing BFP to determine the actual titer. The actual titer of the lentivirus gRNA library is calculated by the following formula: titer = [(F x C_n_)/V] x DF

F: The frequency of GFP-positive cells determined by flow cytometry (0∼1) C_n_: The total number of cells transduced

V: The volume of the inoculum (mL)

DF: The virus dilution factor (total inoculum volume/the volume of original lentivirus gRNA library)

After obtaining the actual titer of the gRNA lentivirus library in iHCF cells, the iHCF cells were then transduced with the appropriate amount of lentivirus to achieve the desired MOI the same way as described above. The media containing virus was removed 1 day post transduction and replaced with complete media. After culturing for an additional 2 days, the cells were selected with 0.25 µg/ml puromycin for 6 days to enrich gRNA transduced cell population before harvesting cells for Perturb-seq. More than 90% of the cells were BFP positive when harvested for Perturb-seq.

#### Generation of CRISPR-Cas9 edited immortalized human cardiac fibroblasts

Pools of immortalized human cardiac fibroblasts (iHCF) (Applied Biological Materials, Inc.) were co-transduced with lentiviruses containing guides against the same enhancer locus and Cas9-Blasticidin lentivirus (Transomic, Inc.). Low-passage (<5) iHCF cells were used since the cells become senescent at higher passage numbers and less able to integrate lentivirus complementary DNA. Triple antibiotic selection with Blasticidin, Puromycin, and Zeocin was applied to infected cells to ensure that cells for Perturb-seq contained the Cas9 expression cassette, ≥1 gRNA targeting the 5’-region flanking the transcription factor binding site, and ≥1 gRNA targeting the 3’-region flanking the transcription factor binding site. Guides were transduced in a semi-arrayed manner at high multiplicity-of-infection (MOI=10) (i.e. each cell pool was transduced with two 5’ guides and two 3’ guides specific for the same locus, but the pair of 5’ and 3’ guides that infected a cell was stochastic). Our approach enabled cardiac fibroblasts to be infected at a high MOI without concern for editing of enhancer loci at multiple chromosomes in the same cell and to enable a high fraction of cells to receive gRNA. After applying antibiotic selection, cell viability and numbers were given time to recover over a period of 23 days, before cells were harvested for Perturb-seq. At the time of harvest, each cell pool contained greater than 100,000 live cells. Cells were harvested by trypsinization, 1x wash in culturing media, and 1x wash and resuspension into PBS + 0.04% BSA at a concentration of 300-500 cells/uL for droplet encapsulation. Viability for all cell pools was ≥ 85%.

#### Perturb-seq library generation and sequencing

Single-cell Perturb-seq (transcriptome and gRNA sequencing at single-cell resolution) was performed using the Chromium 3’ V3.1 method with Feature Barcoding technology for CRISPR screening (10X Genomics, User Guide CG000205 Rev D). Cells in freshly prepared PBS + 0.04% BSA were added to reverse transcription master mix with the appropriate volume of RNAse-free water per manufacturer’s guidelines. The solution containing cells and master mix was encapsulated with gel beads using partitioning oil to generate Gel Bead-in-EMulsions (GEMs) targeting 10,000 recovered cells per sample lane. After encapsulation and reverse transcription, barcoded mRNAs and gRNAs were extracted from GEMs and purified using Dynabeads (MyOne SILANE).

Next, PCR was performed using a primer cocktail (Feature cDNA Primers 1, PN #2000096) for cDNA amplification of cell-endogenous mRNA and targeted amplification of gRNAs containing CaptureSequence2. A total of 11 cycles of cDNA amplification were performed using standard temperature settings, according to manufacturer’s recommendations. During PCR clean-up using SPRIselect beads (Beckman Coulter), the amplified gRNA-containing supernatant fraction was separated from the pelleted fraction, which contained amplified cDNA from endogenous mRNA. Standard library construction of 3’ gene expression libraries (GEX) of the SPRI-purified pelleted fraction was performed according to manufacturer’s recommendations, which consisted of fragmentation, A-tailing, ligation and 12 cycles of sample index PCR to append single-end 8-bp index sequences to each library pool. The gRNA-containing supernatant was SPRI-purified and amplified using two sample index PCR rounds (15 cycles and 5 cycles, respectively) to generate single-indexed final libraries for next-generation sequencing.

Completed libraries were quantified using TapeStation 4200 HSD1000 screentapes and pooled at a molar ratio of 5:1 for GEX:gRNA libraries. Libraries were sequenced on NovaSeq6000 S4 flow cells (Illumina) using the sequencing cycle specifications of 28:8:0:91 (Read1:i7 index:i5 index:Read2). We targeted a total of 75,000 combined GEX and gRNA reads per cell.

#### Pre-processing and filtering of Perturb-seq data

Raw sequencing BCL files were demultiplexed by GEX and gRNA library single-index (i7) barcodes to generate FASTQ files for each library. GEX FASTQ reads were aligned against human reference genome GRCh38-3.0.0, and gRNA FASTQ reads were mapped to a custom feature reference table by calling the *–feature-ref* argument in *cellranger count* (CellRanger software v6.0.0, 10X Genomics [82]). The unfiltered UMI count matrices generated by CellRanger were used for pre-processing of data.

To assign expressed gRNA species to their respective cells, count matrices from different library samples were merged. Since each transduced cell pool only contained a subset of guides against the same target region for each target gene, we first removed all cell barcodes with non-zero expression of gRNAs that were not present in their respective cell pool. This removed 0.03%-3.8% of cell barcodes from each cell pool. We also conservatively removed cell barcodes that expressed less than 100 gRNA UMIs to remove cells with putatively low guide transcription from the analysis (the control cell pool that only express gene-editing enzyme was excluded from this criterion). The expression values of gRNAs were then normalized across cells using the centered log-ratio transformation and scaled [83]. Cells were assigned to perturbation classes based on their expression of gRNAs (i.e. one class per targeted enhancer locus/gene). To remove low-quality or multiplet single-cell transcriptomes, cell barcodes that contained less than 2000 features, greater than 6500 features, and greater than 10% fraction of mitochondrial reads were discarded.

#### Perturb-seq analysis of transcriptional changes following CRISPR KO

Analysis of single-cell RNA-seq was performed using the Seurat package (v3.9.9 [84]).

Raw UMI count values were first transformed using log-normalization. Cell cycle phase was quantified by applying the CellCycleScoring() function to cell cycle genes identified from Tirosh et al, 2016 [85]. Dimensionality reduction using principal component analysis (PCA) and clustering were performed to examine the confounding impact of cell cycle on transcriptional signatures following perturbation. After selecting the top 5,000 most variable features using FindVariableFeatures(), expression values were scaled and the effect of cell cycle was regressed out. The first 40 principal components were used for graph-based assignment of cell neighborhoods (shared Nearest Neighbor), and clustering was performed using the Louvain algorithm (resolution=0.6) to ensure that cells clusters segregated predominantly by biological variables (e.g. gRNA class) and not by technical variables (e.g. cell cycle or library prep batch).

During gene perturbation screening, a fraction of cells may express gRNA but not undergo genetic modification. To increase the signal-to-noise ratio in Perturb-seq experiments, it is critical to distinguish cells from the same gRNA class that were functionally altered by gene editing, from cells that were not impacted by gRNA and gene-editing enzyme transduction. We used Mixscape[86] [84], which uses a Gaussian mixture model to assign posterior probabilities to cells based on gRNA-mediated editing efficiency, to separate perturbed from non-perturbed cells. By specifying cells from the non-targeted control (NTC) gRNA class as a reference for a non-perturbed transcriptome, we found that ∼95% of cells from our assay had a perturbed phenotype, likely due to the high MOI used in our assay. Cells predicted to be unperturbed were excluded from downstream analyses.

Differential expression for all expressed features was performed using MAST [87] to identify transcriptional changes induced by genomic perturbation. Due to the stringent cell filtering criteria used, we considered features with |log_2_-fold-change| > 0.1 and adjusted p-value <0.05 to be statistically significant. Signature genes associated with SNP-containing enhancer regions were identified by removing any features that were differentially expressed between cells only expressing the gene-editing enzyme (editing-enzyme-expressing control) and NTC-transduced cells. We used clusterProfiler [88, 89] version 4.12.0 to run gene set enrichment analysis (GSEA) to search for Kyoto Encyclopedia of Genes and Genomes (KEGG) pathways that may be enriched in differentially expressed genes. Our input list of differentially expressed genes was sorted by |log_2_-fold-change| in expression averaged across cells, weighted by the percentage of cells with detectable expression of that gene.

To visualize single-cell transcriptomic phenotypes, we used linear discriminant analysis (LDA) for dimension reduction to maximize variance across different gRNA classes. The gRNA class was used to distinguish cellular populations, and the first 20 components were used to calculate a UMAP projection for the cells.

## Results

### Characterization of Functional Genomic Features Relevant to Heart Failure and Gene Regulation

#### High-resolution Hi-C identifies 3D chromatin interactions in cardiac myocytes and fibroblasts

Given the critical roles of cardiac myocytes and fibroblasts in the pathogenesis of heart failure, we hypothesized that the integration of chromatin interactions from these cell types with heart disease-associated loci would reveal gene regulatory circuitry underlying heart failure. To capture chromatin interactions, we deeply sequenced large Hi-C libraries to identify long-and short-range interactions across the genome in primary Human Cardiac Myocytes (HCM) cells and Normal Human Cardiac Fibroblasts – Ventricular (NHCFV) cells. HCM and NHCFV are commercially available primary cells derived from non-diseased human tissues.

The HiC-Pro and Juicer pipelines both called nearly 1.5 billion valid interaction pairs after removing duplicate pairs (∼20%). Over 1 billion (∼85%) of these interactions were in *cis* (on the same chromosome) and a large proportion (40-66%) were long-range (≥20-kb) (**Table S1**), indicative of high-quality library construction [42]. According to the script included with Juicer for calculating map resolution based on read depth and genome size, our Hi-C data had 2-kb resolution, which should be sufficient to unambiguously capture interactions involving most enhancers (mean length 540.7±289.8 bp, and up to 1,792 bp) and super-enhancers (mean length 3,324.5±833.7 bp, with a maximum size of 6,345 bp), based on FANTOM5 annotations [90]. Having identified the scaffolding underlying gene regulation in HCM and NHCFV cells, we proceeded to annotate the cell type-specific pairwise interactions with functional elements.

#### Leveraging ATAC-seq to identify acessible & TF-bound loci in cell types relevant to heart failure

A subset of the numerous chromatin interactions within cells represent sites of transcriptional regulation during homeostatic conditions, and their perturbation by disease-associated genetic variation may modulate gene expression during disease. To define these putative regulatory elements in cardiac myocytes and fibroblasts, we performed ATAC-seq in HCM, NHCFV, and immortalized human cardiac fibroblast (iHCF) cells (**Table S2**). Further analysis of these open chromatin regions identified predicted transcription factor binding sites, which provided additional evidence that these elements participate in gene regulation. The ENCODE ATAC-seq pipeline identified ∼180,000 nucleosome-free peaks robustly called across two HCM ATAC-seq libraries, and over 250,000 peaks that were shared by pseudo-replicates (naïve overlap peaks) derived from single NHCFV and iHCF ATAC-seq libraries. The HCM libraries had a higher duplicate fraction and lower library complexity compared to those of the NHCFV and iHCF cells, but with the availability of technical replicates, we were still able to call robust ATAC-seq peaks in HCM cells (Irreproducible Discovery Rate [IDR] peaks which represent higher-confidence, reproducible peaks).

HINT identified over 300,000 footprinted regions each in HCM, NHCFV and iHCF cells. After merging overlapping footprinted regions (footprints containing multiple predicted TF binding sites) within each cell type, we found that 67.6% of unique footprinted regions were common between the primary and immortalized cells. Of these shared regions,157,307 (nearly 40%) also spanned open chromatin regions. The footprinted regions contained 2,694,471 unique motifs corresponding to binding sites corresponding to 840 known TFs in HOCOMOCO and JASPAR. Notably, our analyses identified several well-characterized cardiac-relevant transcription factors such as TEAD and SMAD family proteins.

#### Expression data enable filtering for expressed TFs and genes

We also performed bulk RNA-seq on HCM, NHCFV, and iHCF cells to further prioritize chromatin interactions by quantifying gene expression in our cell types of interest (**Figure S1** and **Table S3).** The expression distribution (log_2_FPKQ) of expressed genes and Pearson correlation coefficient (*r*) reveal that iHCF are more similar to NHCFV than HCMs (**Figure S1A**). NHCFV and iHCF shared expression of cardiac fibroblast genes (e.g. *ACTA2, AIFM2*), and low expression for cardiomyocyte markers (e.g. *TTN, MEF2C*), though we observed differences in certain fibroblast marker genes (e.g. *DCN*), which is expected when comparing primary and immortalized cells (**Figure S1B**). Overall, this transcriptomic data layer informs which genes identified from primary cells can be tested in a cell line that is amenable to genetic perturbation.

### Data Integration to Generate Prioritized Regulatory Circuits

We integrated multiple layers of functional genomic data to identify interacting pairs of SNPs (perturbations in potential disease-related regulatory regions) and TSSs (target genes, whose dysregulation may underlie GWAS signals). **Figures 1A** and **1B** summarize our analysis workflow. We intersected significantly interacting bin pairs from HCM and NHCFV cells (**Figure 1C**) with accessible SNPs and TSSs, to obtain accessible SNP-TSS pairs, and further identified pairs with SNPs in footprints of expressed TFs as well as TSSs of expressed genes. We then annotated SNPs that were in LD with GTEx tissue eQTLs associated with genes corresponding to their interacting TSSs.

**Figure 1:**
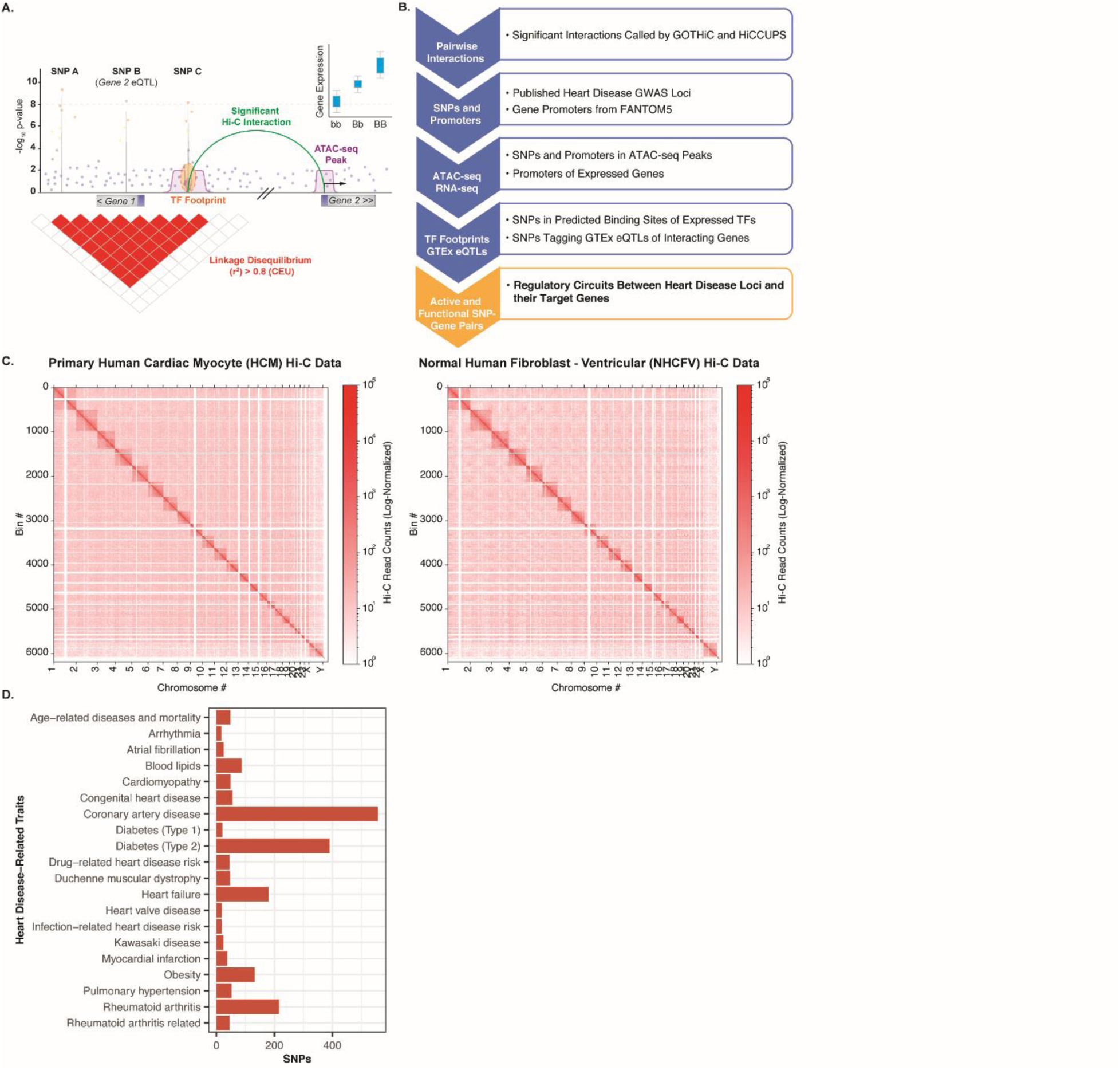
Tracing regulatory circuits from GWAS signals to target genes using functional genomics. **(A)** Graphical summary of analyses annotating GWAS loci with 3D chromatin interaction and open chromatin data to prioritize functional variants and identify target genes that may underlie disease associations. Significant eQTLs further corroborated these SNP-gene relationships. **(B)** Analysis workflow for integration of genetic, expression, and functional genomic data. We layered significant pairwise interactions (chromatin loops) with SNPs and TSSs filtered for accessible regions and highlighted those in expressed TFs’ footprints and expressed target genes, as well as GTEx eQTLs, to identify putative active and functional SNP-gene pairs. **(C)** Genome-wide contact maps of high-resolution Hi-C interactions from primary human cardiac myocytes (HCM) and normal human cardiac fibroblasts –ventricular (NHCFV) **(D)** Number of GWAS SNPs associated with heart disease-related traits, including heart failure.

Our comprehensive search for genetic variants relevant to heart failure yielded 5,534 SNP-phenotype associations, representing 3,880 unique SNPs across heart disease-related pathologies (**Figure 1D**). As expected, there was an over-representation of genetic signals associated with traits likely not directly related to cardiac fibroblast and myocyte pathophysiology, considering our search utilized broad criteria to include heart failure risk factors (e.g. atrial fibrillation, coronary artery disease, hypertension) as well as diseases related to metabolic syndrome. This approach enabled identification of loci linked to multiple cardiometabolic traits, both related and unrelated to cardiac remodeling and fibrosis in heart failure. We augmented the heart disease SNPs with proxy SNPs, resulting in a total of 73,203 variants, representing 3,090 interconnected clusters of correlated SNP pairs (“loci”). Of the 3,880 previously reported SNPs, 389 did not have proxy SNPs with r^2^>0.8 in Europeans. There were 14 SNPs with proxies that did not cluster with other SNP pairs.

We next proceeded to integrate heart disease-associated genetic variants and gene promoters with ATAC-seq, Hi-C, and RNA-seq data in HCM and NHCFV cells (**Figures 2A**). First, we intersected open chromatin regions derived from ATAC-seq in HCM and NHCFV cells with heart disease-related SNPs and genome-wide TSSs. About 15% of accessible SNPs overlapped with HINT footprint regions, and a smaller subset (7-9%) overlapped with TFBSs of known TFs from HOCOMOCO and JASPAR (**Figure 2B, Table S4**). The vast majority of accessible SNPs and TSSs overlapped with significant interactions (chromatin loops) called from our Hi-C data, although the degree of this overlap varied between GOTHiC and HiCCUPS (**Table S5**). These interactions were all intra-chromosomal and predominantly long-range (≥20-kb). While GOTHiC called more short-range interactions overall, HiCCUPs primarily identified long-range SNP-gene pairs (**Table S6A**). We used ATAC-seq and bulk RNA-seq to identify loops involving SNPs in putative active enhancers (accessible to transcriptional machinery) and active genes’ promoters (accessible TSSs of expressed genes) (their properties are described in **Table S6B**). Although putative enhancers containing footprints with expressed TFs were more likely to be functional, we did not hard-filter for this attribute, in order to be inclusive of regions that may contain TF binding motifs that have not been discovered. Using bulk RNA-seq, several candidate interactions were excluded despite their overlap with putative enhancers overlapping GWAS loci, due to almost undetectable expression of predicted target genes (e.g. *GLP2R*, *NAALAD*, *CPNE5*). In total, we identified over 3,000 unique SNP-gene pairs in each cell type using GOTHiC and HiCCUPS (**Figure 2C**). In both cardiac myocytes and fibroblasts, there is a modest overlap between SNP-gene pairs called by GOTHiC (short-range) and HiCCUPS (long-range) (**Figure S2**), and GOTHiC identified the same SNP-gene pairs in both cell types more frequently compared to HiCCUPS. Notably, over two-thirds of SNP-gene pairs involved target genes that were distal to the SNP reported in the original GWAS, and therefore involved novel target genes. This proportion is also consistent with the overabundance of long-range *cis* interactions. A small subset of these SNPs was in LD with significant GTEx eQTLs associated with their target genes in heart regions (∼2%), although a larger proportion (∼4%) corresponded to significant GTEx eQTLs in any human tissue.

**Figure 2:**
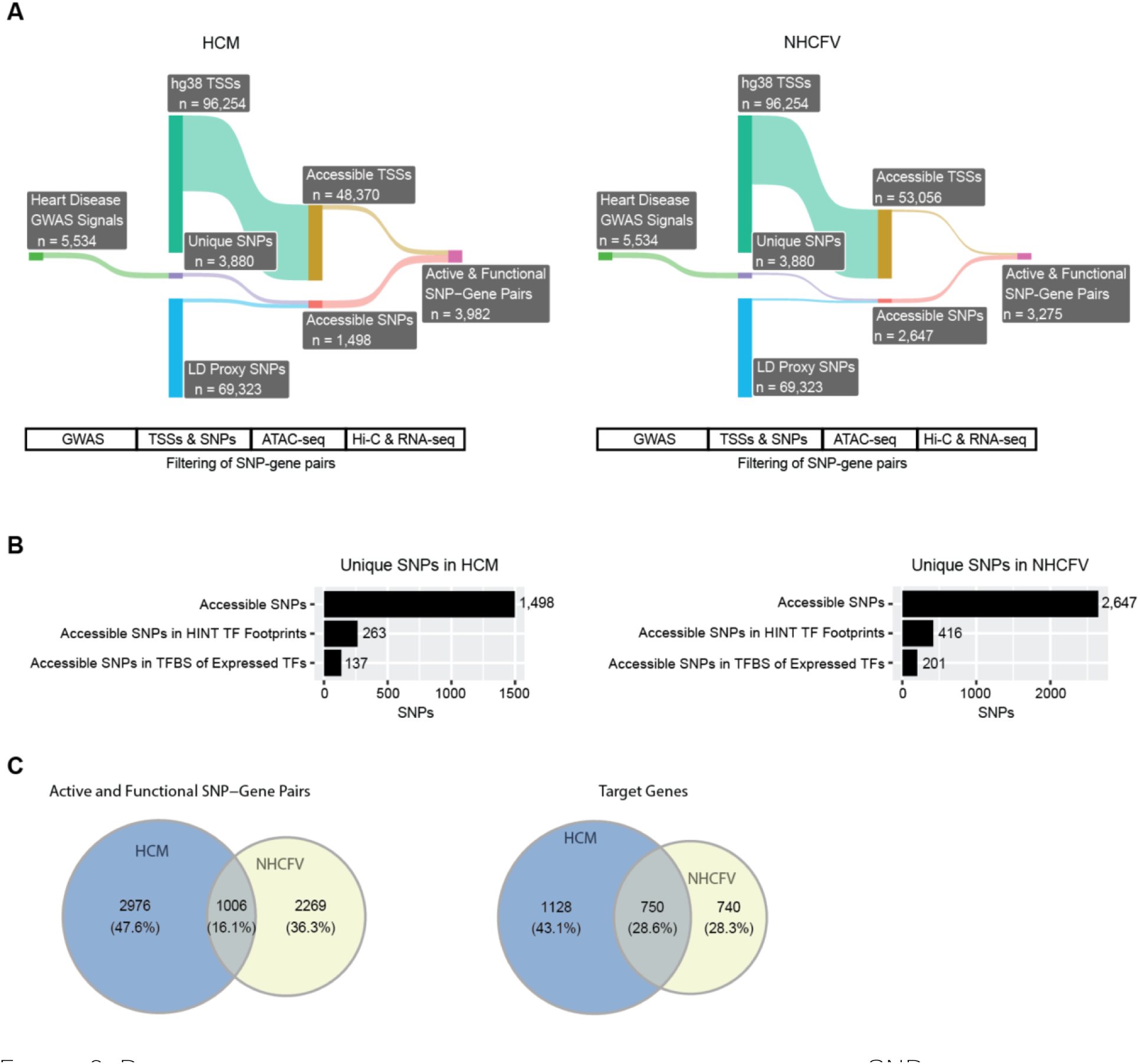
Prioritization of active and functional heart disease-associated SNP-gene pairs in human cardiac myocytes (HCM) and normal human cardiac fibroblasts – ventricular (NHCFV) combining genetics, epigenetics, and transcriptomics. **A)** Sankey diagram of the cell type-specific variation observed in heart disease-associated SNPs and gene accessibility after filtering for open chromatin, chromatin interaction, and expression features, as outlined in Figure 1B. **B)** Quantification of accessible SNPs identified in HCM and NHCFV that are within known transcription factor (TF) footprints of TFs that also have detectable mRNA expression. **C)** Number and overlap of active and functional SNP-gene pairs and corresponding unique target genes identified by GOTHiC and HICCUPS across HCM and NHCFV cells.

In addition to loops, topologically associating domains (TADs) are larger-scale structures often thought of as “local neighborhoods” which constrain interactions between genes and regulatory elements. Hence, we assessed the proportion of our identified SNP-gene pairs where the variant and gene are both contained within the same TAD across various Hi-C resolutions in both NHCFV and HCM cells. The total number of TADs inferred in each cell type varied considerably depending on algorithm and resolution (**Table S7**). For TADs computed using the directionality index or insulation score, a maximum of 41% of SNP-gene pairs were found within the same TAD, while TADs discovered via the arrowhead algorithm fared slightly better, with a maximum of 63% of SNP-gene pairs falling within the same TAD. In contrast, across all resolutions, the more recently published GRiNCH algorithm [53] inferred TADs that *at a minimum* showed 86% of SNP-gene pairs falling into the same TAD.

Notably, the GRiNCH TADs tended to be smaller than those inferred by other tools, and SNP-gene pairs *not* contained within the same TAD were consistently farther apart than those within the same TAD (**Figure S3**). The full extent of TADs’ practical relevance is debated [91], but these findings suggest functionally important TADs may be more readily identifiable by modern methods that utilize smoothing and non-negative matrix factorization, such as GRiNCH. For this reason, we will only present GRiNCH TAD calls in the main results figures, but we will provide results with the other TAD-calling algorithms in Supplemental Figures. Our TAD analyses highlight the importance of applying different methods at various scales on deeply-sequenced Hi-C data, to robustly identify novel regulatory interactions.

Our high resolution Hi-C data affords us the unprecedented ability to identify direct interactions between individual SNPs (proxies for enhancers and super-enhancers) and TSSs.Our analysis of these data revealed promiscuous interactions between SNP- and TSS-spanning bins (**Figure S4**). These situations may arise when a SNP bin interacts with multiple bins containing distinct genes’ TSSs (enhancers with multiple target genes) (**Figure S4A**) or if two or more distinct SNP bins interact with a TSS bin (target genes interacting with multiple regulatory elements) (**Figure S4B**). More than half (81.4% in HCM and 63.3% in NHCFV) of the SNPs in expressed TFs’ footprints were in loops with more than one target gene, suggesting that the enhancers they perturb may be regulatory hubs affecting multiple target genes. On the other hand, about 30% of HCM and NHCFV expressed genes’ TSSs interacted with more than one SNP in footprints of expressed TFs. These cases may represent multiple perturbations of a single enhancer regulating a target gene, perturbation of two or more distinct enhancers regulating a disease-related target gene, or both.

While these promiscuous interactions appear primarily driven by complex underlying regulatory circuitry, we also encountered ambiguous SNP-TSS pairings, which are primarily driven by technological limitations. Despite the high resolution of our Hi-C data, 2-kb windows are still larger than both SNPs and TSSs (mean length 25.2±21.3 bp and up to 249 bp in the FANTOM5 data, respectively). Hence, there will inevitably be cases of bins containing two or more adjacent or overlapping TSSs interacting with SNP-spanning bins, as well as situations where bins spanning two or more indistinguishable SNPs interacting with TSS-overlapping bins:

i. Ambiguous Target Gene (**Figure S4C**): One-fifth of footprint SNPs interacted with bins containing TSSs of two or more distinct genes (19.7% in HCM and 21.6% in NHCFV).
ii. Ambiguous SNP (**Figure S4D**): Nearly half (46.6% in HCM and 44.9% in NHCFV) of the SNP-gene pairs involved a target gene interacting with more than one SNP.
iii. Ambiguous Target Gene Bin (**Figure S4E**): Cases where gene TSSs span more than one 2-kb bin are not an issue, as they result in redundant interactions but no information loss. These occurred at about 10% frequency in HCM and NHCFV.

### SNP-Gene Interactions Involving Known Heart Disease Genes

For a subset of the significant SNP-gene interactions, our multi-omic approach revealed high concordance between chromatin regulation, gene expression, and involvement of TFs and target genes with established roles in the pathophysiology of heart disease, cardiac fibrosis, and heart failure (**Table 3**). Such gene regulatory networks reveal epigenetic contexts that underlie human genetics-driven disease-modifying changes in cellular behavior.

#### rs7772537 (in FOXJ3 footprint) and *TBC1D32*, rs1919875 and *GJA1* (at chr6q22.31)

rs7772537 is an intergenic SNP that is in LD with another intergenic SNP rs12664873, which was identified in a GWAS of atrial fibrillation (OR 1.08 [1.05-1.12], *P*=1.2E-8) [18], **(Table 1 Figure 3**). rs7772537 resides within a TF footprint of Forkhead Box J3 (FOXJ3) in cardiac fibroblasts, and physically interacts with the promoter of *TBC1 Domain Family Member 32* (*TBC1D32*) over 850-kb away. An additional independent atrial fibrillation signal was reported at rs1919875, a variant between rs7772537 and *TBC1D32* (OR 1.08, *P*=3.0E-11) [28]. rs1919875 is in an open chromatin region and tags the intergenic variant rs13195459 (mapped to *HSF2*) but interacts with G*ap Junction Protein Alpha 1* (*GJA1*) over 400-kb away. The rs7772537-*TBC1D32*, and rs1919875-*GJA1* loops are proximal to TAD boundaries called by all four algorithms, particularly Arrowhead and Insulation Score (**Figure 3**, **Figure S5**). Notably, neither rs7772537 nor rs1919875 tag a GTEx eQTL for either *TBC1D32* or *GJA1*. Our bulk RNA-seq shows that *GJA1* is highly expressed in HCM, NHCFV, and iHCF cells, while *TBC1D32* is lowly expressed in these cells.

**Table 1:** Significant interactions between heart disease GWAS SNPs involving known heart disease genes and/or transcription factors

| SNP | SNP Position | Associated Diseases | Mapped Genes | Min GWAS p-value | Min GTEX eQTL p-value | TF Footprints (motif match) | Cell Type | Gene | Gene Expression (FPKQ) in NHCFV, HCM, and iHCF cells | TSS(s) | TSS Range | Loop Source | Loop Bin Distance | Loop Reads | Min Loop q-value |
| --- | --- | --- | --- | --- | --- | --- | --- | --- | --- | --- | --- | --- | --- | --- | --- |
| rs1919875 | chr6: 121,849,906 | AF | *AL603865.1*,<br>*HMGB3P18*,<br>*HSF2* | 3.0E-11 | N/A | N/A | NHCFV | *GJA1* | 325.5<br>160.5<br>40.7 | P2 | chr6:121,435,591-121,435,766 | GOTHIC | 414,000 | 14 | 3.28E-22 |
| rs7772537 | chr6: 122,185,395 | AF | *GJA1*,<br>*LOC105377979*,<br>*RNU1-18P* | 1.0E-08 | N/A | FOXJ3 (13.7) |  | *TBC1D32* | 1.0<br>2.7<br>0.1 | P1, P2,<br>P3 | chr6:121,334,406-121,334,739 | GOTHIC<br>HiCCUPS | 850,000 | 6-68 | 9.15E-19 |
| rs1680636 | chr10:44,180,492 | CAD | *CXCL12*,<br>*LINC00841* | 2.0E-16 | N/A | ARNT (11.3)<br>ARNTL (14.1)<br>*ATF3* (10.8)<br>BHLHE40 (13.0)<br>BHLHE41 (11.6)<br>CREB3L2 (12.1)<br>MLX (13.7)<br>SREBF1 (11.0)<br>SREBF2 (9.8)<br>TFE3 (14.1)<br>TFEB (12.6) | NHCFV<br>HCM | *CXCL12* | 145.0<br>72.9<br>498.4 | P1 | chr10:44,376,910-44,385,105 | HiCCUPS | 200,000 | 58 | 5.84E-04 |
| rs7549250 | chr1:154,431,860 | MI, CAD | *IL6R* | 6.0E-16 | 1.6E-20 | KLF12 (10.2) | NHCFV | *IL6R* | 0.4<br>2.4<br>0.3 | P19 | chr1:154,405,199-154,434,982 | GOTHIC | 24,000 | 11 | 2.14E-84-1.18E-24 |
| rs4129267 | chr1:154,453,787 | C-reactive protein |  | 2.0E-14 | 2.4E-09 | ELK3 (11.1)<br>REL (13.6) | NHCFV |  |  | P18 |  |  | 18,000 | 21 | 1.61E-54 |
| rs35346340 | chr15:90,884,641 | MI, CAD | *FURIN*, *FES* | 2.0E-25 | 1.1E-15 | *EGR1* (11.9)<br>*EGR2* (11.9)<br>*KLF15* (11.3)<br>KLF6 (10.5)<br>MAZ (13.1)<br>MESP1 (12.8)<br>MXI1 (10.9)<br>*SNAI1* (12.0)<br>SP1 (9.9)<br>SP2 (10.1)<br>SP3 (12.2)<br>SP4 (10.2)<br>VEZF1 (10.2)<br>ZEB1 (11.7)<br>ZFX (11.1) | NHCFV<br>HCM | *FURIN* | 77.7<br>15.0<br>65.1 | P4, P6,<br>P10, P11 | chr15:90,868,580-90,875,744 | GOTHIC | 12,000 | 12 | 1.21E-24 |

Genes and transcription factors involved in cardiac fibrosis and/or heart disease are in red.
AF: Atrial fibrillation; CAD: coronary artery disease; MI: myocardial infarction.
NHCFV: Normal human cardiac fibroblasts (ventricular); HCM: Human cardiac myocytes; iHCF: Immortalized human cardiac fibroblasts

**Figure 3:**
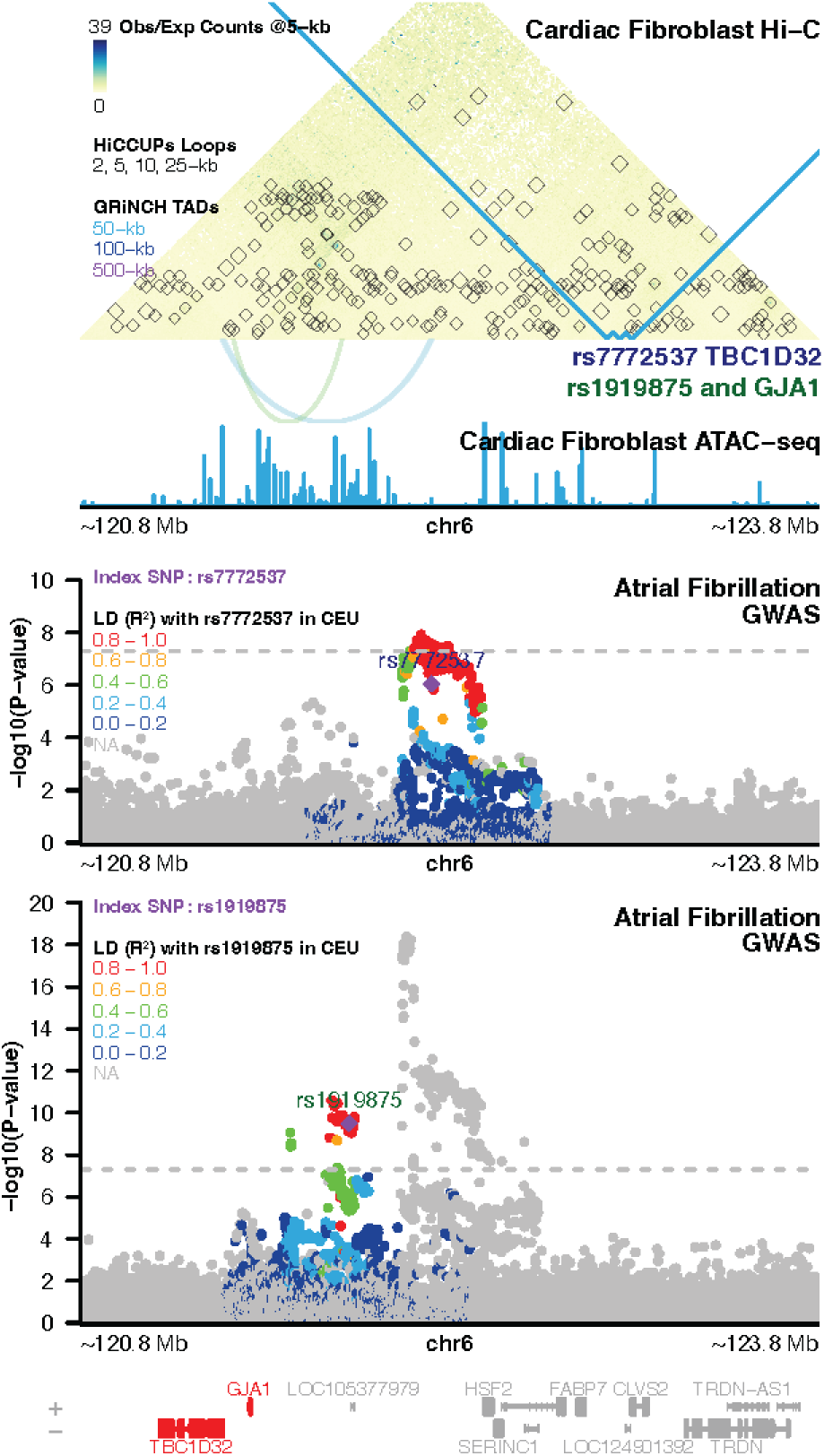
Significant interactions between an atrial fibrillation-associated putative regulatory elements, at rs7772537 and rs1919875, with *TBC1D32 and GJA1*, respectively, in cardiac fibroblasts, overlaid with GRiNCH TAD calls.

#### rs1680636 and *CXCL12* (at chr10q11.21)

rs1680636 is an intergenic SNP in footprints of several TFs and interacts exclusively with *C-X-C motif chemokine ligand 12* (*CXCL12*) in a 200-kb loop, in both cardiac fibroblasts and cardiac myocytes (**Table 1**, **Figure 4**). *CXCL12* is highly expressed in HCM, NHCFV, and iHCF cells. rs1680636is an intergenic SNP mapping to both *CXCL12* and *LINC00841*, and is a tag SNP for the intergenic variant rs2457480, which is significantly associated with CAD (OR=1.111 [1.109-1.115], *P*=2.0E-16) [17] (**Table 1**, **Figure 4**). Consistent with the abundance of interactions at the locus surrounding *CXCL12*, boundaries of larger TADs were identified in both cell types across most TAD calling tools (**Figure 4**, **Figure S6**). *CXCL12,* whose serum levels have been linked to atherosclerosis and atrial fibrillation [92], is highly expressed in HCM, NHCFV, and iHCF cells, but rs1680636 and its proxies are not CXCL12 eQTLs in GTEx tissues.

**Figure 4:**
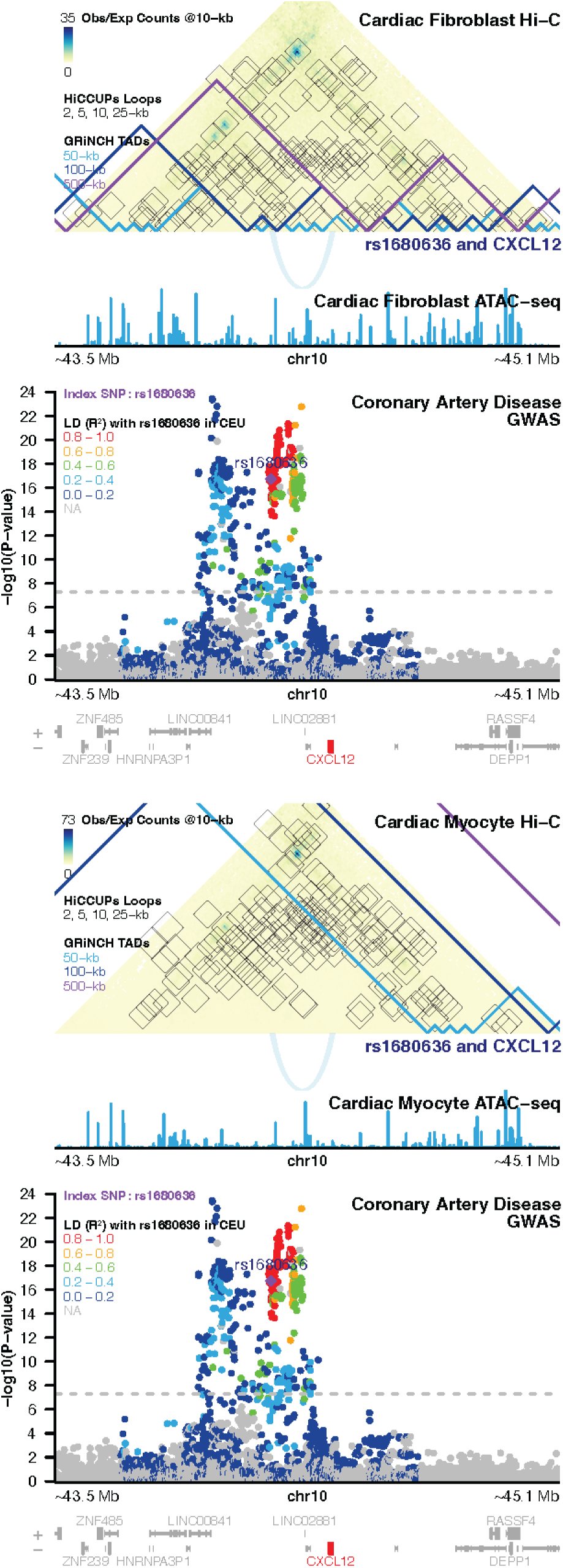
Significant interaction between a coronary artery disease-associated open chromatin at rs1680636 and *CXCL12* in both **A)** cardiac fibroblasts and **B)** myocytes, overlaid with GRiNCH TAD calls.

#### rs7549250, rs4129267 and *IL6R* (at chr1q21.3)

rs7549250 and rs4129267 both map to intronic regions of cytokine receptor *Interleukin 6 Receptor* (*IL6R*), and their proxy SNPs have been associated with multiple heart disease-related traits. Proxies for rs7549250 were identified in GWASs of myocardial infarction (MI) (rs12118721, OR 1.06 [1.04-1.08], [15]) and coronary artery disease (CAD) (rs6689306, OR 1.06 [1.04-1.08], [15]; rs6689306, OR 1.05 [1.03-1.07], [22]; rs4845625, OR 1.0458 [1.0453-1.0463], [17]), while a proxy SNP of rs4129267 was associated with decreased C-reactive protein levels (rs4537545, -11.5% [-14.4% to -8.5%, =1.3E-12) (**Table 1**, **Figure 5**). Both rs7549250 and rs4129267 overlap with several transcription factor footprints and interact with *IL6R* TSSs ∼20-kb away in cardiac fibroblasts but not cardiac myocytes. While there were no large GRiNCH TAD calls at this locus, other algorithms identified large-scale TADs nearby (**Figure S7**). rs7549250 is an *IL6R* eQTL in the left ventricle (*P*=1.6E-20) and atrial appendage (*P*=1.4E-06), while rs4129267 does not tag any *IL6R* heart eQTLs but is a significant eQTL of *IL6R* in tibial artery (*P*=2.4E-09), whole blood (*P*=4.6E-07), esophagus muscularis (*P*=1.9E-06), and transverse colon (*P*=1.5E-05). However, *IL6R* is lowly expressed in both primary cell types as well as immortalized fibroblasts.

**Figure 5:**
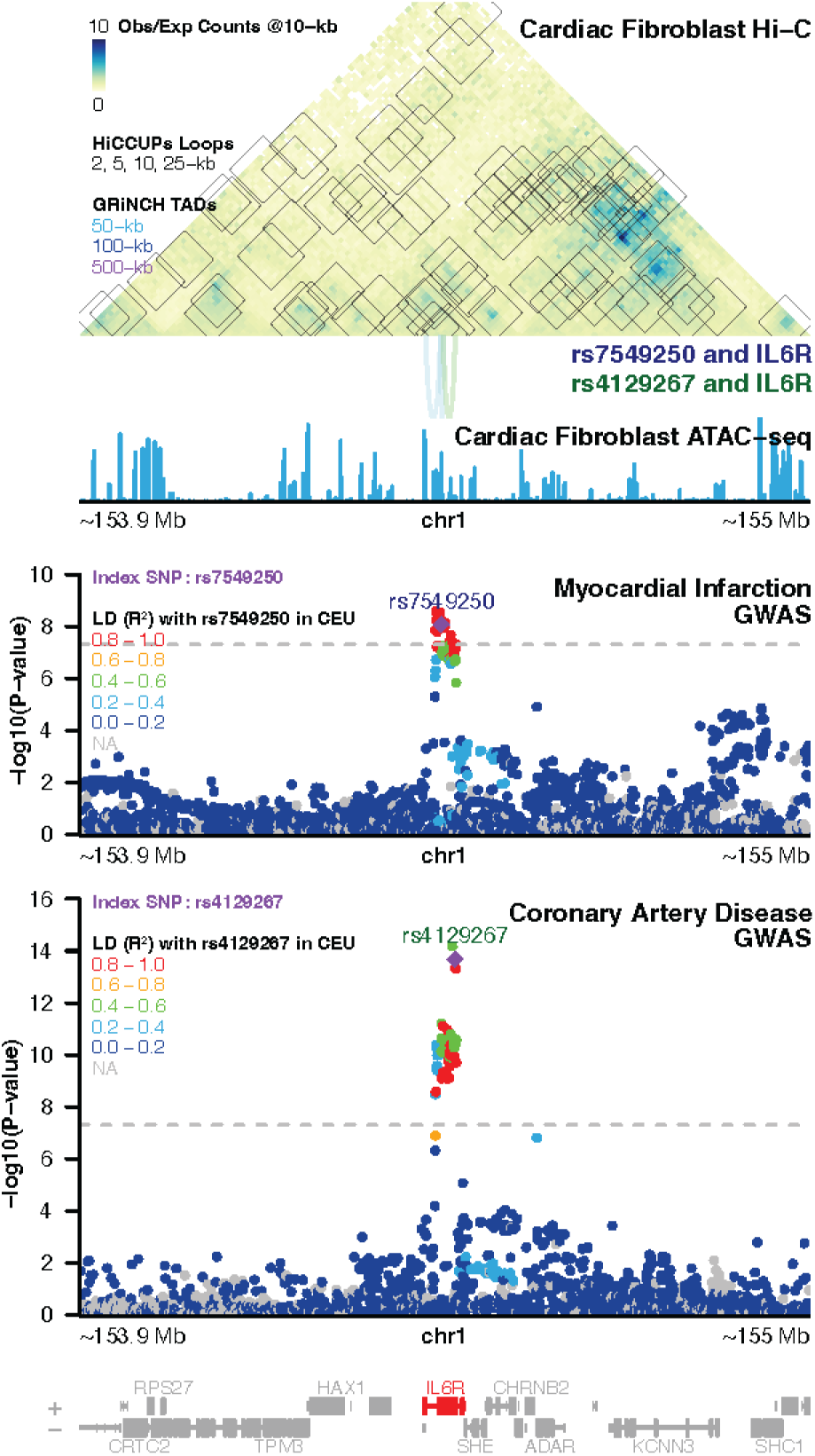
Significant interactions between myocardial infarction- and coronary artery disease-associated open chromatin at rs7549250 and rs4129267 with *IL6R* in cardiac fibroblasts, overlaid with GRiNCH TAD calls.

#### rs35346340 and *FURIN* (at chr15q26.1)

rs35346340 is in LD with two heart disease GWAS SNPs: rs2521501 was associated with CAD (OR=1.06 [1.04-1.08, *P*=5.0E-08) and MI (OR=1.07 [1.04-1.09], *P*=1.5E-07) [15], while rs4932373 was associated with MI only (OR=0.93 [0.92-0.94])[17] (**Table 1**, **Figure 6**). rs35346340 also resides in cardiac fibroblast and cardiac myocyte ATAC-seq footprints corresponding to the binding sites of multiple TFs, including KLF15. Interestingly, although rs35346340 is in the first intron of *Feline Sarcoma* (*FES*), a gene adjacent to *FURIN*, this SNP only interacts with the *FURIN* in our Hi-C data. GRiNCH, directionality index, and insulation score did not find any TADs at this locus, but the arrowhead algorithm found larger TADs in this region, particularly cardiac fibroblasts (**Figure S8**). rs35346340 is a significant *FURIN* eQTL in esophageal mucosa (*P*=2.0E-13) and a nominally significant eQTL of this gene in aortic artery (*P*=1.0E-04), and *FURIN* is highly expressed in both primary cardiac fibroblasts and myocytes, as well as immortalized fibroblasts.

**Figure 6:**
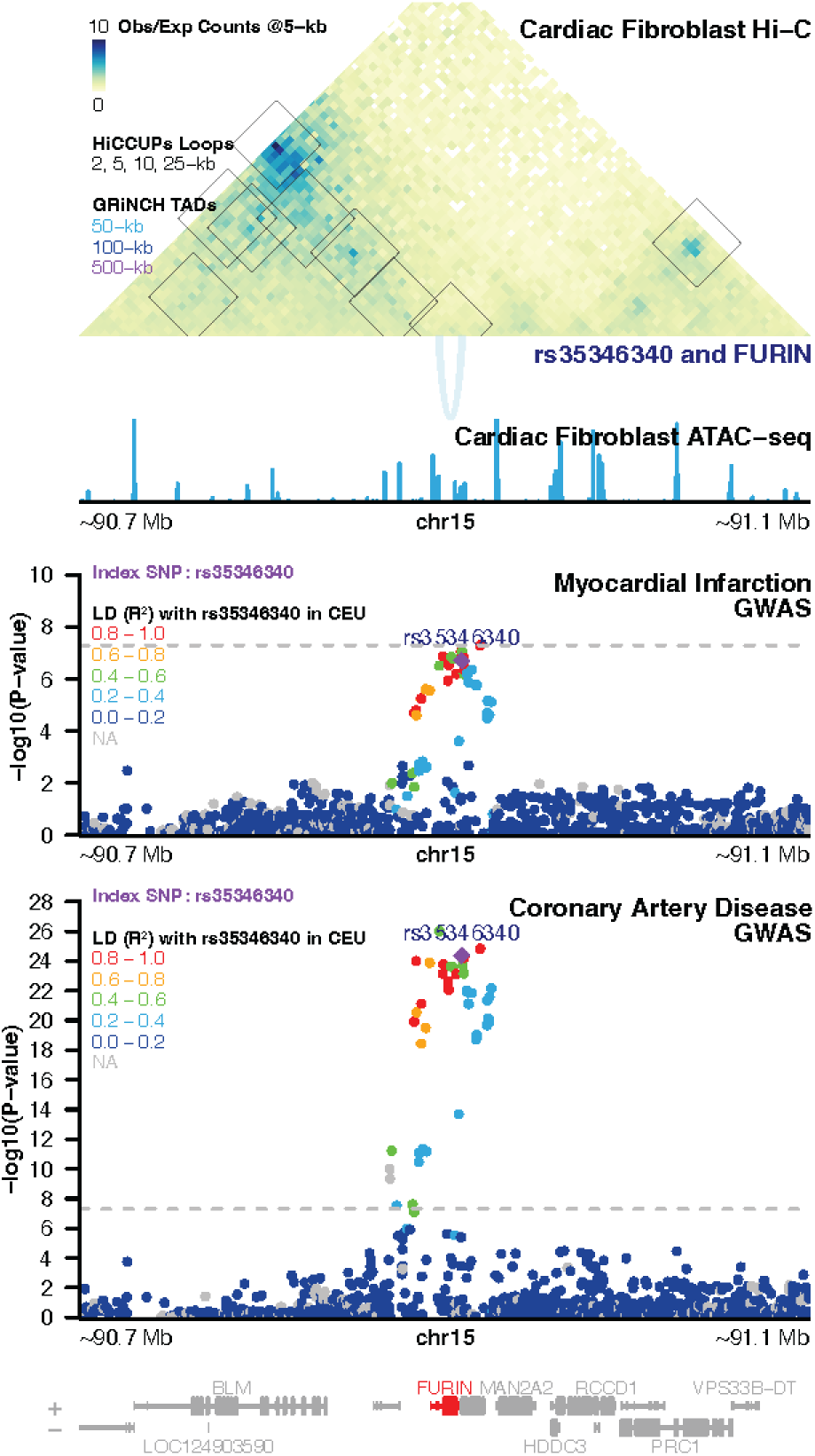
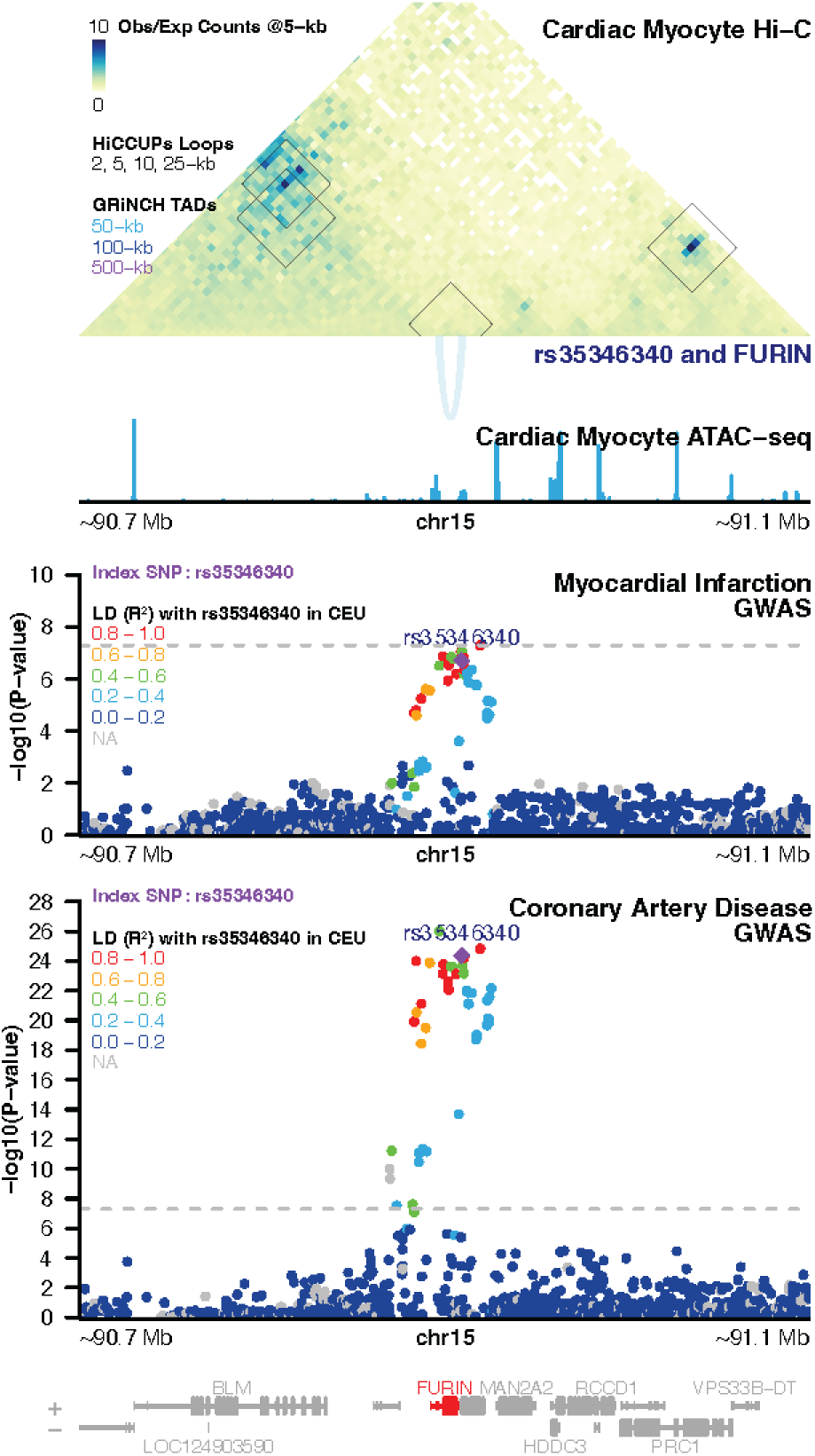
Significant interaction between myocardial infarction- and coronary artery disease-associated open chromatin at rs35346340 with *FURIN* in both **A)** cardiac fibroblasts and **B)** cardiac myocytes, overlaid with GRiNCH TAD Calls.

We further confirmed that identified genes were expressed in diseased adult human heart cells by querying human cardiac myopathy single-nuclear RNA-seq (snRNA-seq) and spatial transcriptomic datasets and explored how expression changes across cell types, disease states, and zones in the heart [93, 94]. All five target genes and corresponding heart disease-associated transcription factors involved in the prioritized SNP-gene interactions were expressed in both fibroblasts (3-33% of cells for target genes, 1-36% of cells for transcription factors) and cardiomyocytes (1-26% of cells for target genes, 0.1-35% of cells for transcription factors) (**Figure S9A**). Of the target genes, fibroblasts more highly express *CXCL12*, *TBC1D32*, *IL6R* and *FURIN*, while cardiomyocytes are the highest expressors of *GJA1* of any cell type in the heart. Comparison of target gene expression in snRNA-seq and spatial data revealed differing gene expression levels among acute myocardial infarction (AMI) patients compared to ischemic and non-cardiomyopathy patients (**Figure S9B-C**). In addition, we observed co-localization between fibroblasts/cardiomyocytes and each target gene in spatial transcriptomic human myocardial infarction data (**Figure S9D-E**). Expression is generally concentrated in the ischemic zone of AMI patients and less prevalent in control patients (**Figure S9F**). Together, the variation in gene expression across disease types and heart regions suggests that tissue-dependent contexts, in addition to genetic variation, coordinate to regulate target gene expression in cardiac fibroblasts and myocytes.

### Validation of SNP-function relationships using single-cell profiling of enhancer deletions

To measure the effect of non-coding, disease-associated loci on predicted target genes as well as their broader transcriptomic consequences, we used a modified direct capture Perturb-seq approach to knock out genomic loci in immortalized human cardiac fibroblasts (iHCF) **(Figure 7A)**. For each locus, two guide RNAs flanking each side of the predicted transcription factor (TF) binding footprint were designed to induce deletion of the TF footprint. Guides that flanked the 5’-end of the TF footprint were cloned into vectors containing Puromycin resistance (PuroR), and guides that flanked the 3’-end of the TF footprint were cloned into vectors containing Zeocin resistance (ZeoR). All guides contained a modified scaffold stem loop sequence that could be hybridized to NextGEM gel beads and enable reverse transcription.

**Figure 7:**
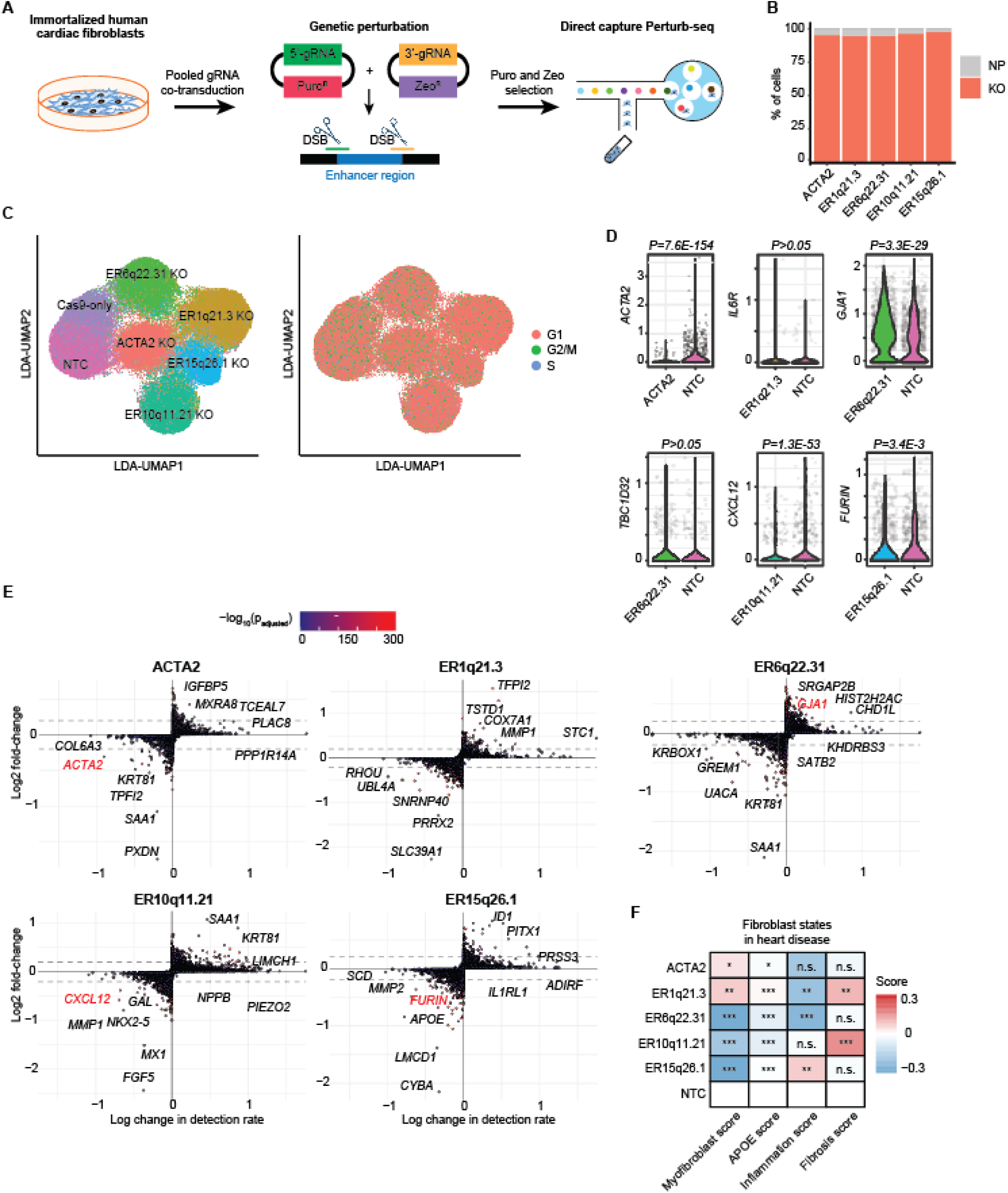
Validation of enhancer regulation of target genes in human cardiac fibroblasts using single-cell Perturb-seq. **A)** Diagram of gene perturbation strategy to delete enhancer regions and perform Perturb-seq. DSB: double-stranded DNA break. **B)** Stacked bar plot showing percentage of cells with effective KO alteration as inferred by Mixscape. NP: non-perturbed, KO: knockout **C)** Linear-discriminant analysis UMAP projections of cardiac fibroblasts, labeled by gene perturbation (left) or by cell cycle state (right). **D)** Violin plots showing gene expression changes in predicted target genes of specific enhancer sites. Exon-targeting *ACTA2* gRNA was used as a positive control for gene editing activity. *P*-values were calculated using MAST differential expression analysis. **E)** Scatter plots showing transcriptome-wide changes in per-cell detection rate and log-fold-change between a perturbation class and the non-targeting control (NTC) group. Black gene labels show the top differentially expressed genes that are unique to each perturbation. Red gene labels show predicted target genes of enhancer regions. **F)** Gene module analyses of genetically edited fibroblasts using gene markers of fibroblast cell states during human heart failure (myofibroblast differentiation, APOE^+^ fibroblasts, inflammation, and fibrosis). *P-*values were calculated using Wilcoxon rank sum test. *P<0.05, **P<0.001, ***P<0.00001, n.s.: not significant

Guides that targeted the same locus were transduced into cells, and dual antibiotic selection using Puromycin and Zeocin ensured that all surviving cells contained at least one guide flanking either side of the TF footprint. Upon recruitment of Cas9 to the loci, CRISPR-mediated deletion would excise and inactivate the TF footprint region and, consequently, genes that are regulated by these regions. We opted not to use CRISPR interference (CRISPRi), since our regions of interest ranged from 700-2800 base pairs and CRISPRi has been reported to inhibit regions less than 300 base pairs [95].

Since primary cardiac fibroblasts are unable to continuously proliferate after over twenty passages, we used low-passage iHCFs for Perturb-seq, which can maintain cell doubling times after Cas9 selection and gRNA selection. Separate iHCF cultures were transduced with a pool of guides against chromosome regions 1q21.3, 6q22.31, 10q11.21, and 15q26.1 (we use the “ER” prefix to designate the edited enhancer region). We also included a separate pool of guides targeting exonic regions of *ACTA2*, which is highly expressed in cardiac fibroblasts and served as a positive control gene to ensure successful CRISPR-mediated knockout. Twenty-three days after gRNA and Cas9 co-transduction, cell cultures were harvested and processed using the Chromium NextGEM 3’ V3.1 chemistry to measure transcriptome-wide mRNA expression and guide RNA expression.

After filtering cell transcriptomes for gene quality, we detected 7,519 to 16,436 cells for each perturbation, across two replicate libraries per perturbation group. We detected a high abundance of gRNA expression within cells transduced with gRNA (average of 200-900 gRNA per cell) but not with cells only receiving the gene-editing enzyme. Average guide RNA expression levels per cell were typically 70-200 times higher than endogenous mRNA levels. For each locus, cells were analyzed using Mixscape [86] to distinguish perturbed from non-perturbed cells, and >95% perturbation efficiency was observed for all groups **(Figure 7B)**. After regressing out cell cycle effects **(Figure 7C)** and clustering cells by perturbation group using linear discriminant analysis (LDA), single-cell differential expression analysis was performed using MAST to characterize the transcriptional consequences of knocking out loci. Cells receiving *ACTA2* gRNA showed a significant decrease in *ACTA2* mRNA expression **(Figure 7D-E)**, confirming effective gRNA delivery and Cas9 activity. For all groups of cells receiving guides, including the non-targeting guides, we observed differential expression of DNA-sensing *IFN* and *OAS* family genes, and hence these genes were excluded from consideration.

For regions ER6q22.31, ER10q11.21, and ER15q26.1 whose predicted target genes were *GJA1*/*TBC1D32*, *CXCL12*, and *FURIN*, respectively, we observed significant differential expression of at least one target gene following gene editing at each predicted regulatory element (**Figure 7F**). *CXCL12* and *FURIN* were significantly downregulated following deletion of ER10q11.21, and ER15q26.1, while *GJA1* was upregulated after we deleted ER6q22.31. When comparing NTC vs Cas9-only cells these target genes were either not significantly differentially expressed or showed opposing differential expression trends, suggesting that expression changes observed following gene editing were a result of regulatory element deletion, rather than artefacts of cell stress associated with cell transduction and antibiotic selection. Excision of region ER1q21.3 did not significantly downregulate predicted target gene *IL6R*, nor did we detect changes in *TBC1D32* expression after ER10q11.21 knockout. The extremely low expression levels of both *IL6R* and *TBC1D32* from bulk (**Table 1**, **Figure S10A-B**) and single-cell RNA-seq (**Figure S9**) indicate that the specific cell line used may not be a reliable model system to interrogate regulation of these particular genes. KEGG-based pathway analysis of differentially expressed genes following knockout found enrichment of additional genes and pathways relevant to heart failure (**Table S8, Figure S10C**). Following deletion of enhancer region ER10q11.21 (predicted to interact with *GJA1* and *TBC1D32*), pathway analysis identified *COX7A1* downregulation in multiple heart disease-relevant pathways (cardiac muscle contraction (hsa04260), diabetic cardiomyopathy (hsa05415), and vascular smooth muscle contraction (hsa04270)). In addition, deletion of the enhancer interacting with *FURIN* (ER15q26.1) resulted in decreased expression of its substrate, *MMP2*, and of *ACTG1*, a core enrichment gene in dilated (hsa05414) and hypertrophic (hsa05410) cardiomyopathy pathways. No heart disease-related KEGG pathways were detected from the differentially expressed genes of ER1q21.3 and ER10q11.21 knockout cells.

We next performed in-depth analyses of transcriptional changes induced by CRISPR KO of predicted enhancers at loci associated with coronary disease. Using multi-gene signatures previously defined by single-cell RNA-seq of fibroblast cell states in acute myocardial infarction [93], we observed changes in genes associated with fibrotic (*POSTN, COMP, COL1A1, THBS4, COL3A1, FBN1, PPRX, FOSX1, MEOX1, RUNX1, EDNRA*), inflammatory (*CCL2, CCL11, THBS1, PTGDS, GPC3*), and myofibroblast (*ACTA2, TAGLN*) states after CRISPR KO (**Figure 7F**). Of note, knockout of ER15q26.1, which reduced expression of myofibroblast-promoting protease *FURIN*, also reduced expression of myofibroblast gene signatures. This suggests that perturbation of these sites has the potential to affect the expression of interacting genes (identified by Hi-C) as well as downstream effectors that may alter cardiac fibroblast function.

Overall, our Perturb-seq screen validates the regulatory relationships between three of four predicted enhancer regions harboring heart disease-associated GWAS loci: 6q22.31-*GJA1* (atrial fibrillation), 10q11.21-*CXCL12* (coronary artery disease), and 15q26.1-*FURIN* (myocardial infarction and coronary artery disease). We also identified possible roles for these enhancer sites in modulating global gene expression programs associated with fibrosis and inflammation, reinforcing the direct pathophysiological roles of these non-coding sites on cellular phenotypes.

## Discussion

In this study, we traced regulatory circuits from heart disease SNPs to their target genes, by integrating genome-wide interaction, epigenomic, and transcriptomic data from human cardiac fibroblasts and cardiac myocytes. We identified 3D chromatin interactions using both GOTHiC and HiCCUPs to ensure that we captured both short- and long-range loops [51].

Notably, across both Hi-C loop calling tools, over 2/3 of SNP-gene interactions prioritized for regulatory relationships involve target genes that were not mapped to the original SNP, a proportion consistent with findings from orthogonal approaches [96]. ATAC-seq provided evidence to suggest that GWAS loci reside within putative regulatory elements, and transcriptomic profiling confirmed the cell-specific expression patterns of target genes. Deletion of predicted enhancer regions demonstrated that these regions actively regulate cellular transcriptional programs and, in some cases, are sufficient to modulate target gene expression. Our multi-omic, integrative approach revealed novel disease genes that otherwise would not have been identified by considering the genes most proximal to GWAS signals alone.

Our target genes and the TFs corresponding to binding motifs in ATAC-seq footprints were enriched for several known cardiac remodeling and fibrosis genes and TFs, including Cx43, further affirming the value of integrating GWAS signals with multi-omics data from relevant tissues. Such integrative multi-omics approaches are especially important for heart failure, where disease heterogeneity in addition to other factors have precluded a comprehensive understanding of its genetic basis, particularly polygenic forms of heart failure [97]. Below, we summarize the evidence suggesting pathophysiological relevance of several of our identified transcription factors and target genes.

### Transcription Factor FOXJ3’s Target Genes *GJA1* and *TBC1D32*

G*ap Junction Protein Alpha 1* (*GJA1*) encodes Connexin 43 (Cx43), a gap junction component involved in pathogenic trans-differentiation of fibroblasts into myofibroblasts, which migrate to injured areas to deposit extracellular matrix. Specifically, Cx43 mediates Transforming Growth Factor B (TFG-β)-induced fibroblast-to-myofibroblast phenoconversion in rats, and Cx43 knockdown and overexpression respectively inhibited and stimulated the promoter activity of α-SMA (a myofibroblast marker) in cardiac fibroblasts [98]. Furthermore, another study found that knockdown of *Connective Tissue Growth Factor* (*CCN2*/*CTGF*), which encodes a primary target and downstream effector of TGF-β, reduced expression of Cx43 [99]. These prior findings and our own prioritization results are all consistent with the notion that *GJA1* plays a key role in heart failure-associated myocardial fibrosis. *TBC1 Domain Family Member 32* (*Tbc1d32*) was identified in a recessive forward genetic screen in fetal mice in which the authors performed dense phenotyping for congenital heart defects and exome sequencing to identify recessive mutations. Specifically, the *Tbc1d32* mice showed laterality defects that may be related to sonic hedgehog (SHH) signaling transduced or modulated by the cilium [100]. Although deletion of ER_6q22.31 did not alter *TBC1D32* expression, we detected decreased expression of *COX7A1*, which encodes the contractile muscle-specific and major cardiac isoform of cytochrome c oxidase (Cox). Homozygous and heterozygous *Cox7a1* knockout mice demonstrate dilated cardiomyopathy [101]. While these target genes will require deeper characterization to clarify potential novel therapeutic avenues, our analyses point to the key gene targets, TFs, and pathways implicated at this atrial fibrillation –associated GWAS locus.

### Target Gene *CXCL12* and Transcription Factor ATF3

C-X-C motif chemokine ligand 12 (*CXCL12*), also known as *Stromal Cell Derived Factor 1* (*SDF1*), is a ligand for C-X-C motif receptor 4 (CXCR4), a G protein-coupled receptor with roles in angiogenesis, myocardial ischemia, and injury-induced restenosis [102]. In our data *CXCL12* interacts with CAD-associated rs1680636, which is in a binding motif of Activating Transcription Factor 3 (ATF3), a basic leucine zipper family TF involved in cardiac remodeling. Specifically, ectopic expression of *Atf3* in *Atf3* knockout mice inhibited angiotensin II-induced cardiac fibrosis and hypertrophy [103]. Furthermore, previous cancer studies have implicated an interaction between *Atf3* and *CXCL12* in tumor progression [104, 105]. Taken together, these results suggest *CXCL12* may be an important target of *Atf3* involved in the etiology of heart failure and associated cardiovascular outcomes.

### Target Gene *IL6R* and Transcription Factor KLF12

The Interleukin 6 Receptor (IL6R) ligand IL-6 is a cytokine involved in cell growth and differentiation, as well as immune response, and is tightly associated with the liver-derived inflammatory biomarkers C-reactive protein (CRP) and fibrinogen. All three are significantly associated with increased risk of heart disease [106]. IL-6 may mediate cardiac remodeling consequent to Ang II, as IL-6 knockout mice were resistant to Ang II-induced cardiac dysfunction, myocardial inflammation, and fibrosis compared to wild-type mice [107].Consistent with the notion that *IL6R* may be directly involved in inflammation that leads to heart disease, a meta-analysis found that a non-synonymous SNP in *IL6R* (rs8192284 [p.Asp358Ala]) and its proxy rs7529229 are robustly associated with coronary heart disease (per-allele OR 0.95 [0.93-0.97], *P*=1.5E-05), as well as increased IL-6 and decreased levels of downstream effectors CRP and fibrinogen [108].Interestingly, another significant SNP at this locus, rs7549250, is in a TF footprint of Kruppel-like factor 12 (KLF12), which is associated with rheumatoid arthritis. The FDA-approved monoclonal antibody tocilizumab, which is commonly prescribed for rheumatoid arthritis and other chronic inflammatory diseases, competitively inhibits IL6R, and, like rs8192284, is associated with increased IL-6 and decreased CRP and fibrinogen. However, the drug has a directionally inconsistent effect on cardiovascular disease due to its pro-atherogenic lipid profile [108]. These findings suggest crucial roles for IL6R in heart disease and point to a need to more deeply characterize KLF12 targets and mechanism(s) of action in cardiac-relevant tissues.

### Transcription Factors EGR1, KLF15, and SNAI1, and Target Gene *FURIN*

Another SNP, rs35346340, is in the TF footprints of several TFs which have been associated with phenotypes related to heart failure: Early Growth Response 1 (EGR1), Kruppel-like factor 15 (KLF15), and Snail 1 (SNAI1). EGR1 is a C2H2-type Zn finger transcription factor and *Egr1* knockout mice show increased cardiac fibrosis [109]. In cardiac fibroblasts, *KLF15* inhibits expression of *CCN2*/*CTGF* [7], thereby preventing cardiac fibrosis. SNAI1 is a Zn finger transcriptional repressor, and *Snai1* was co-expressed with the fibrosis marker periostin in post-MI mice, suggesting a role in post-MI *de novo* fibrosis [110]. The target gene of rs35346340, *Furin, Paired Basic Amino Acid Cleaving Enzyme* (*FURIN*), is important to the renin-angiotensin system and sodium-electrolyte balance, and was prioritized in an integrative polygenic analysis of GTEx aorta eQTLs and GWAS data as the likely causal gene underlying a systolic blood pressure association signal [111]. Expression of *FURIN* has also been found to be elevated in canines [112] and mice [113] with heart failure. The lower FURIN expression and reduced inflammatory signature we observed after rs35346340 knockout suggest that the enhancer at this locus promotes a fibroblast phenotype that is detrimental during heart failure. Intriguingly, we found that deletion of the enhancer interacting with *FURIN* resulted in differential expression of γ-actin (*ACTG1*), which is the major actin isoform in the heart and important in cardiac muscle contraction [114].

### SNP-Gene Pairs and eQTLs

We made note of SNP-gene pairs in which the footprint SNP or a proxy were eQTLs of the target gene. While this represents additional evidence of a regulatory relationship, GTEx eQTLs are based on whole-tissue information and these relationships may not be consistently recapitulated in our regulatory circuits derived from more homogeneous primary cell populations. In addition, GTEx eQTLs were limited to SNPs within 1-Mb of gene TSSs [115]. It is further worth noting that the utility of eQTL overlap for understanding GWAS signals, regardless of eQTL discovery specifics, may be relatively limited. In addition to the possibility that systematic differences in evolutionary pressures and discovery methods for eQTL and GWAS may preclude holistic overlap [32], it is also important to recognize that the measured intermediate molecular phenotype for eQTLs – transcript expression – is not the most downstream effector of biological function (i.e. protein abundance). Indeed, some studies have found surprisingly low overlap (less than or equal to ∼50%) between protein quantitative trait loci (pQTL) and eQTL [116–118], whereas others have explicitly observed significant buffering effects wherein final protein levels differ substantially from measured transcript levels[119–121]. Such findings suggest incorporation of further proteomics data could be beneficial for characterizing GWAS hits’ targets and mechanisms of action. Unfortunately, proteomic assays have historically lagged behind transcriptomic assays, but recent technological and methodological advances should enable more comprehensive protein assessments and, consequently, better functional characterization of disease-associated variants[122, 123].

### Limitations

As mentioned, our Hi-C data, even at high 2-kb resolution, could not always resolve to a single gene’s TSS, depending on local genetic architecture. Hi-C is also limited to assaying pairwise interactions, and we thus may miss further characterizations of genes and variants with multiple interaction effects (although some proportion of these will be found in orthogonal pairwise interactions). Additionally, our use of ATAC-seq data to identify enhancer regions is by no means comprehensive. Some enhancers are only active in stimulated cells, as evidenced by studies of immune cells [124, 125], and may not have been represented in our ATAC-seq data of wild-type primary cells. Furthermore, not all intergenic ATAC-seq-derived open chromatin peaks necessarily represent enhancers, with some likely harboring other types of cis regulatory elements with different impacts on gene expression. Addition of other types of epigenomic (i.e. histone modification) data could aid in more deeply characterizing the direction and nature of regulatory relationships underlying GWAS signals.

Our discovery paradigm also relies on filtering GWAS SNPs by open ATAC peaks. While this is a reasonable strategy, it will inevitably be impacted by the robustness of the identified ATAC peaks, with highly variable peaks necessitating thorough experimental validation. With more samples and replicates per sample, inferred peaks would be less impacted by biological noise as well as technical variation. This could enable less experimental validation of individual hits, as the putative candidates identified are more likely to be true positives. . It is also possible that some of the SNPs we examined exert their effects on heart disease in a different cell type; most GWAS signals are likely cell-type specific, and the types we collected data for here are by no means comprehensive.

CRISPR-mediated deletion of identified non-coding regions containing SNPs and TF binding footprints was able to both modify expression of predicted target genes > 10 kilobases away and alter fibroblast gene signatures across inflammatory and fibrotic statuses associated with heart disease. Intriguingly, we observed that region-specific deletion caused some target genes to be upregulated (*GJA1*), while other genes (*CXCL12, FURIN*) were downregulated.

This can be caused by several interdependent reasons. First, DNA elements may function either as transcriptional enhancers or repressors, which in turn could assign a protective or exacerbating role for target genes. Second, genetic ablation of the entire TF binding site in our experimental setting may modify TF and co-factor binding differently than variation of only a single nucleotide change in diseased individuals, thus representing a distinct impact on enhancer sites and target gene transcription. Third, our Perturb-seq experiments focused on primary cardiac fibroblasts from healthy donors, and their baseline gene expression may differ from fibroblast gene expression during disease or biology in cardiomyocytes , in which we found some of the SNP-gene pairs. Future studies that investigate the physical interactions between SNPs, TF binding activity, and transcriptional regulation of target genes can further elucidate how enhancer regions and their target genes change during disease.

## Conclusions

There is a multitude of GWAS signals for numerous phenotypes, including heart disease-related traits antecedent to heart failure. While discovery of these and other disease-associated variants is, in and of itself, useful for screening and lifestyle interventions, deeper functional characterization of such associations is needed to identify gene targets and corresponding novel therapies that can directly address even reverse disease. One major challenge in pursuit of this goal is determining which epigenetics data to collect: in most cases, resources are limited, both fiscally and biologically. Our study demonstrates the strong utility of incorporating tissue-relevant Hi-C and ATAC-seq data into GWAS signal characterization, as they enable a robust linking of SNPs, target genes, and cis regulatory elements. These are by no means the only epigenomic data worth collecting to understand and functionally characterize disease-associated variants, but they represent a compelling initial avenue that helps address one of the largest outstanding issues in GWAS: connecting disease-associated variants to their gene targets. Using DNA interaction and epigenetic data to construct regulatory circuits between non-coding GWAS loci and their target genes, we identified genes and transcription factors with established relevance to heart failure and provide evidence to suggest that SNP-gene pairs discovered using our method may represent novel targets for therapeutic intervention.

## Acknowledgements

We would like to thank Ryan Potts, Simon Jackson, and Bill Richards for providing helpful scientific input. We would also like to acknowledge Andy Rampersaud and Vanessa Arias for providing technical support.

## Author Contributions

YH and RG designed the study and conceived the experiments. DL, YA, JC, JL, TY, JY, HZ, and CL performed experiments. RG, JMA, and DL processed data. RG, JMA, DL, and IE analyzed data. CL and YH oversaw data generation and analysis. CL, SW, KL, BA, and YH advised and provided resources to support the project. RG, DL and IE wrote the manuscript. RG, DL, IE, JMA, KL, BA, CL and YH finalized the manuscript.

RG designed the study, conceived experiments, processed data, analyzed data, wrote the paper, and revised the paper

DL performed gene editing experiments, processed data, analyzed data, wrote the paper, and revised the paper

IE analyzed data, wrote the paper, and revised the paper

JC performed cell culture and gene editing experiments JL performed cell culture and gene editing experiments YA participated in data analysis to prioritize targets

TY performed sequencing experiments and processed data JY performed cell culture experiments

HZ performed sequencing experiments BP contributed to data analysis

SW supported the project

CL advised experiments and revised the paper

YH designed the study, conceived the study, conceived and advised experiments, oversaw data analysis, and revised the paper

## SUPPLEMENTARY FIGURES

**Figure S1:**
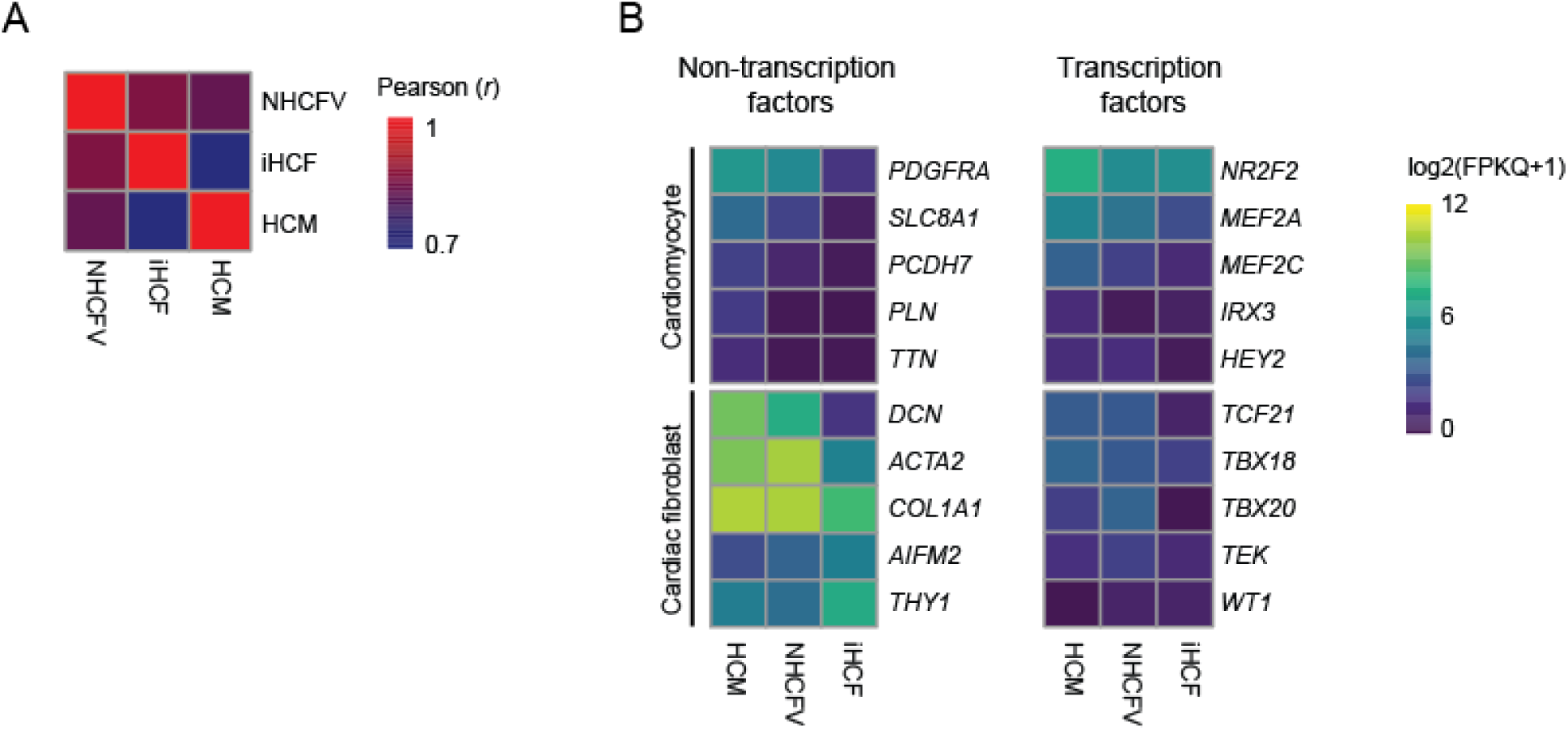
Bulk RNA-seq of HCM, NHCFV, and iHCF cells. **A)** Pearson correlation coefficients of transcriptome-wide expression levels in HCM, NHCFV, and iHCF. **B)** Heatmap of expression levels (log_2_(FPKQ+1) of cardiomyocyte marker genes and cardiac fibroblast marker genes in cell types used in this study.

**Figure S2:**
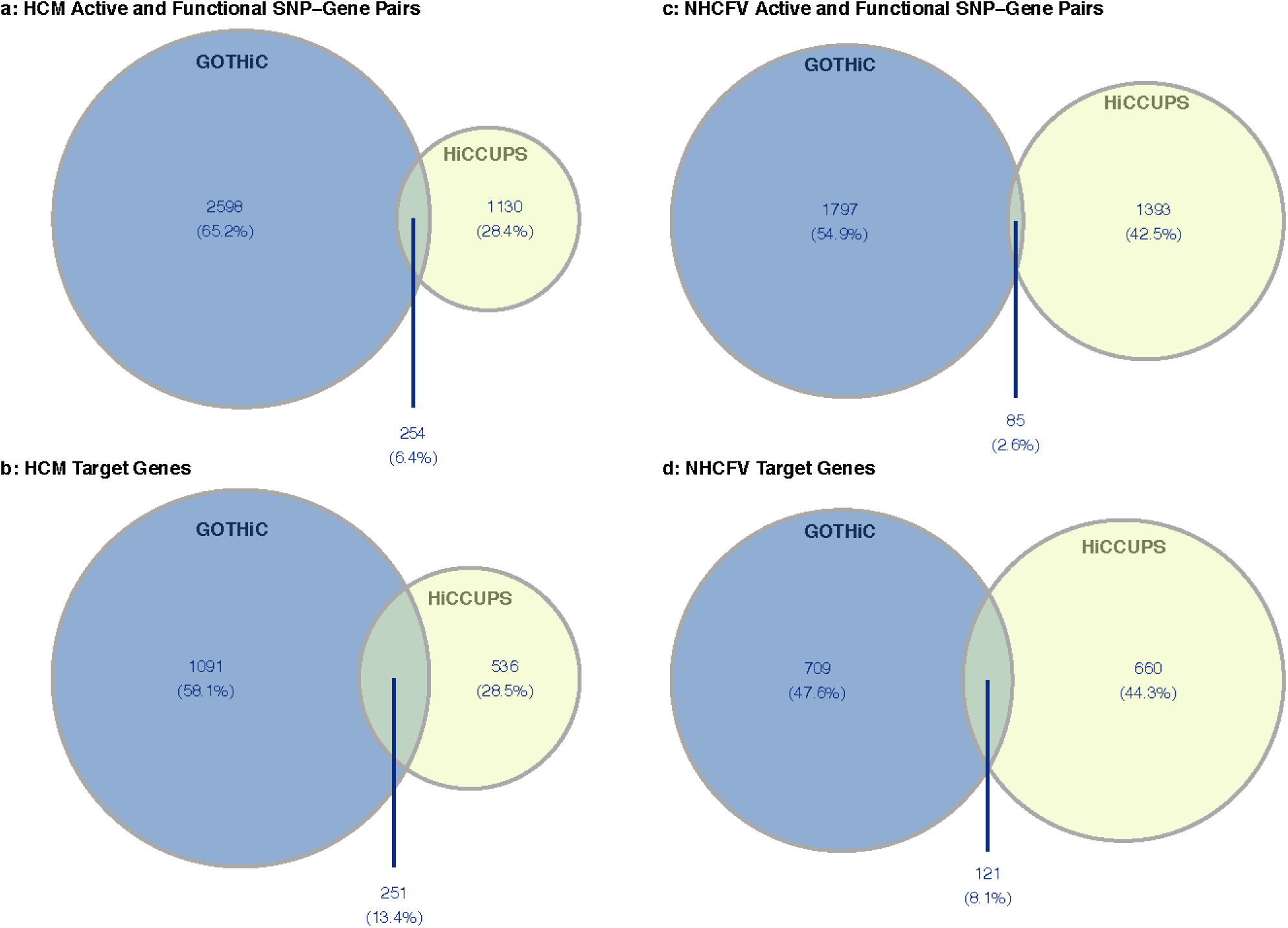
Number and overlap of active and functional SNP-gene pairs identified by GOTHiC and HiCCUPS in HCM and NHCFV cells (**A** and **C**) and corresponding unique target genes (**B** and **D**).

**Figure S3:**
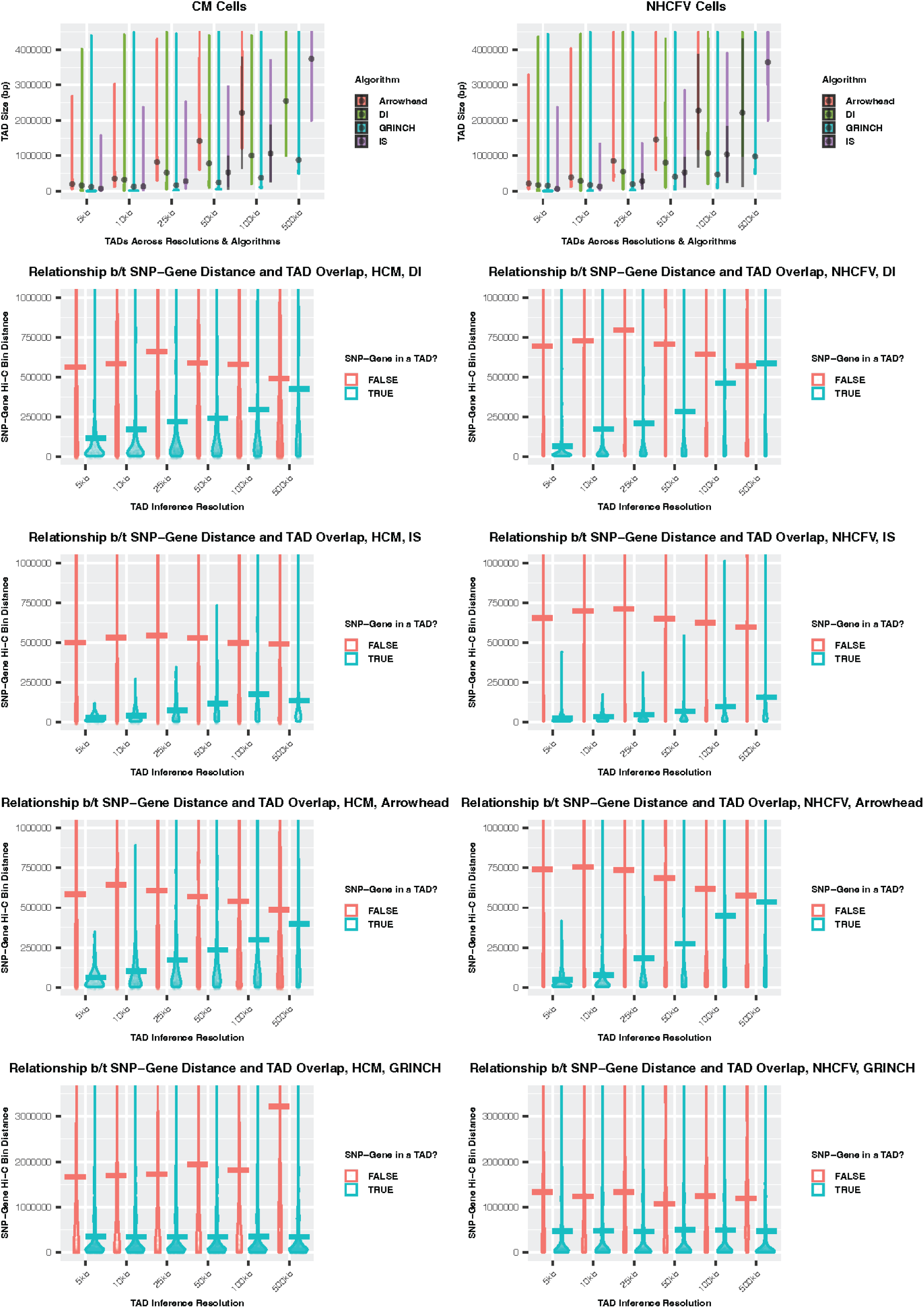
Inter-algorithm examination of TAD sizes and SNP-gene pair distances stratified by TAD inclusion status. **A)** and **B)** show the distribution of TAD sizes inferred across resolutions in HCM and NHCFV cells, respectively, using the four different algorithms applied. Note arrowhead TADs do exist at 500kb resolution but are not visible in this distribution due to their large size (smallest found was 4 Mb). **C)** and **D**) For TADs called in HCM and NHCFV using directionality index (DI), we show distance between the Hi-C bins of an identified SNP-gene pair (y-axis) across different TAD inference resolutions (x-axis), stratified by whether the SNP-gene pair could be found within the same TAD (colors). **E-J)** Similar to a and b, but for insulation score (IS), arrowhead, and GRiNCH-defined TADs, respectively.

**Figure S4:**
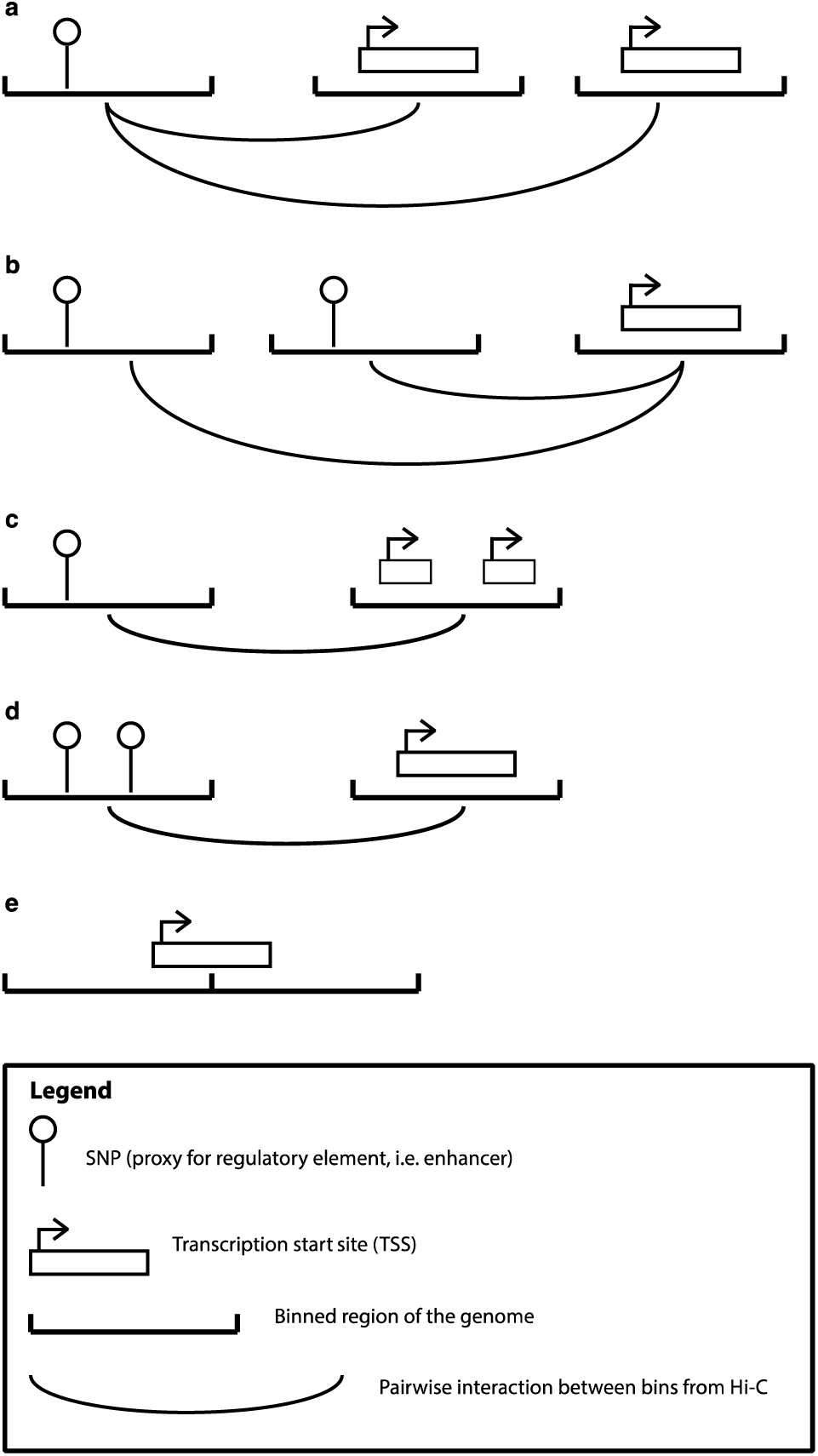
Different types of promiscuous SNP-gene interactions in pairwise Hi-C data. **A)** SNPs interacting with multiple target genes, **B)** Target genes interacting with SNPs, **C)** SNPs with ambiguous target genes, **D)** Target genes interacting with ambiguous SNPs, **E)** Target genes spanning multiple Hi-C bins.

**Figure S5:**
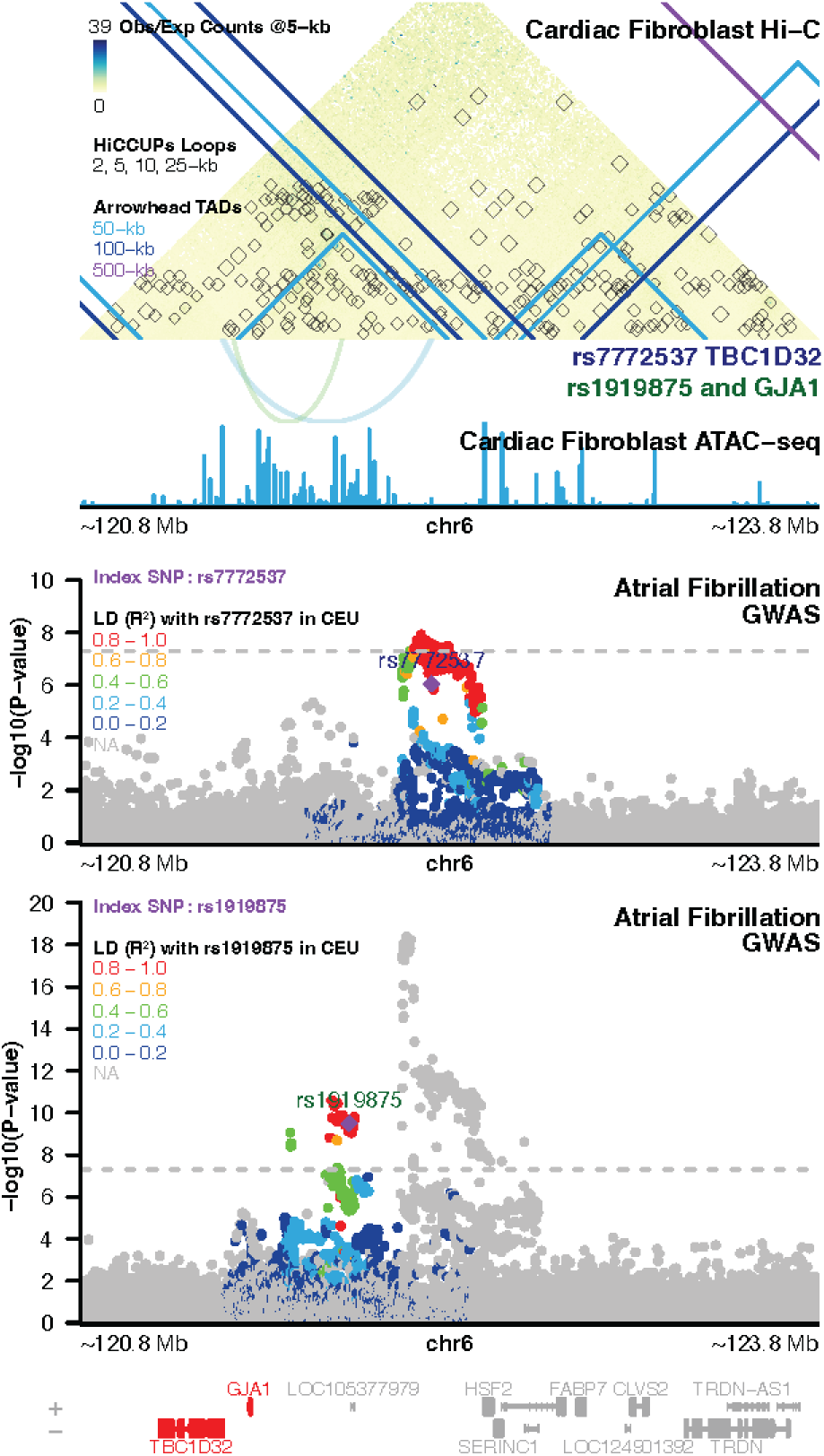
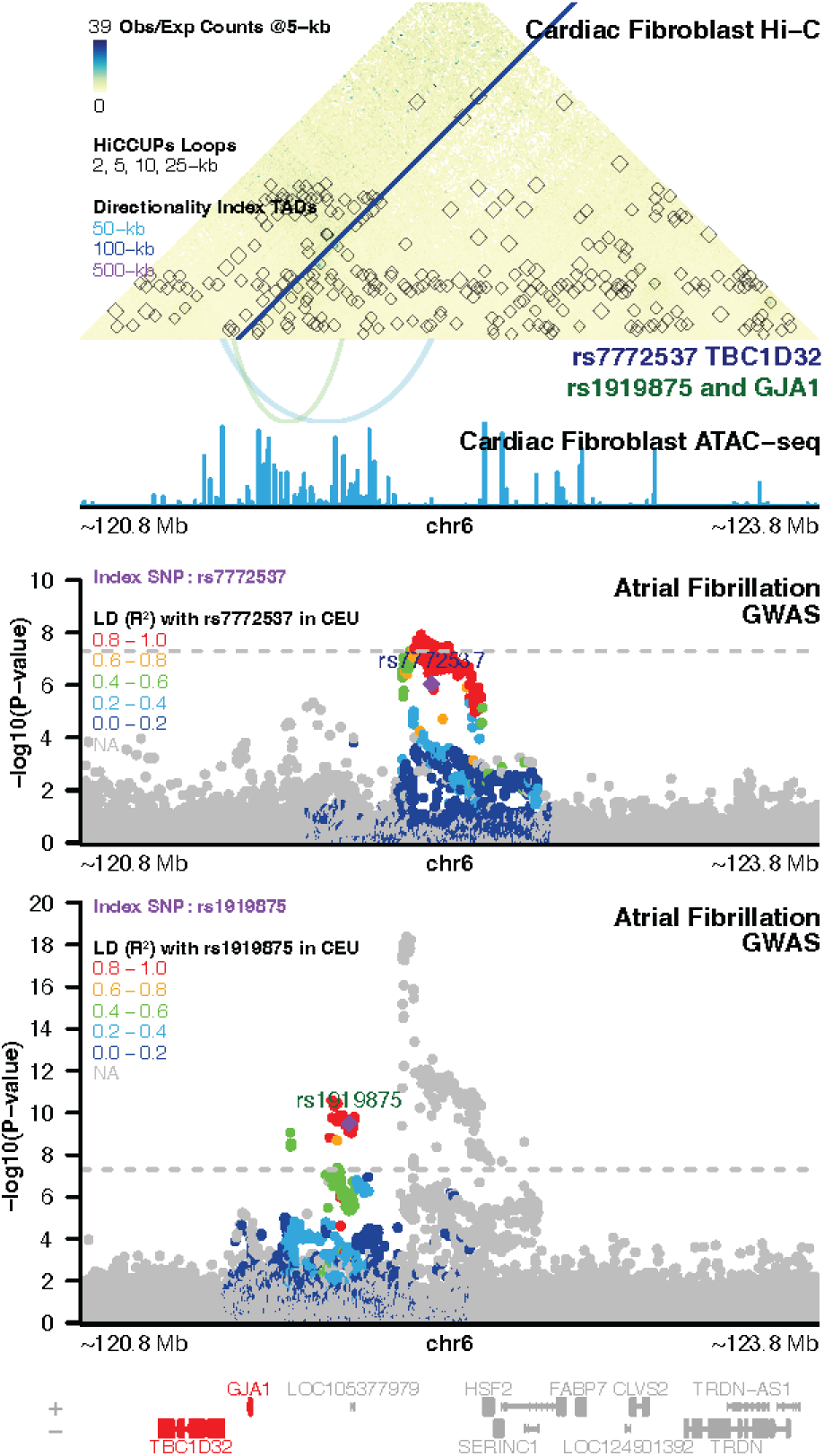
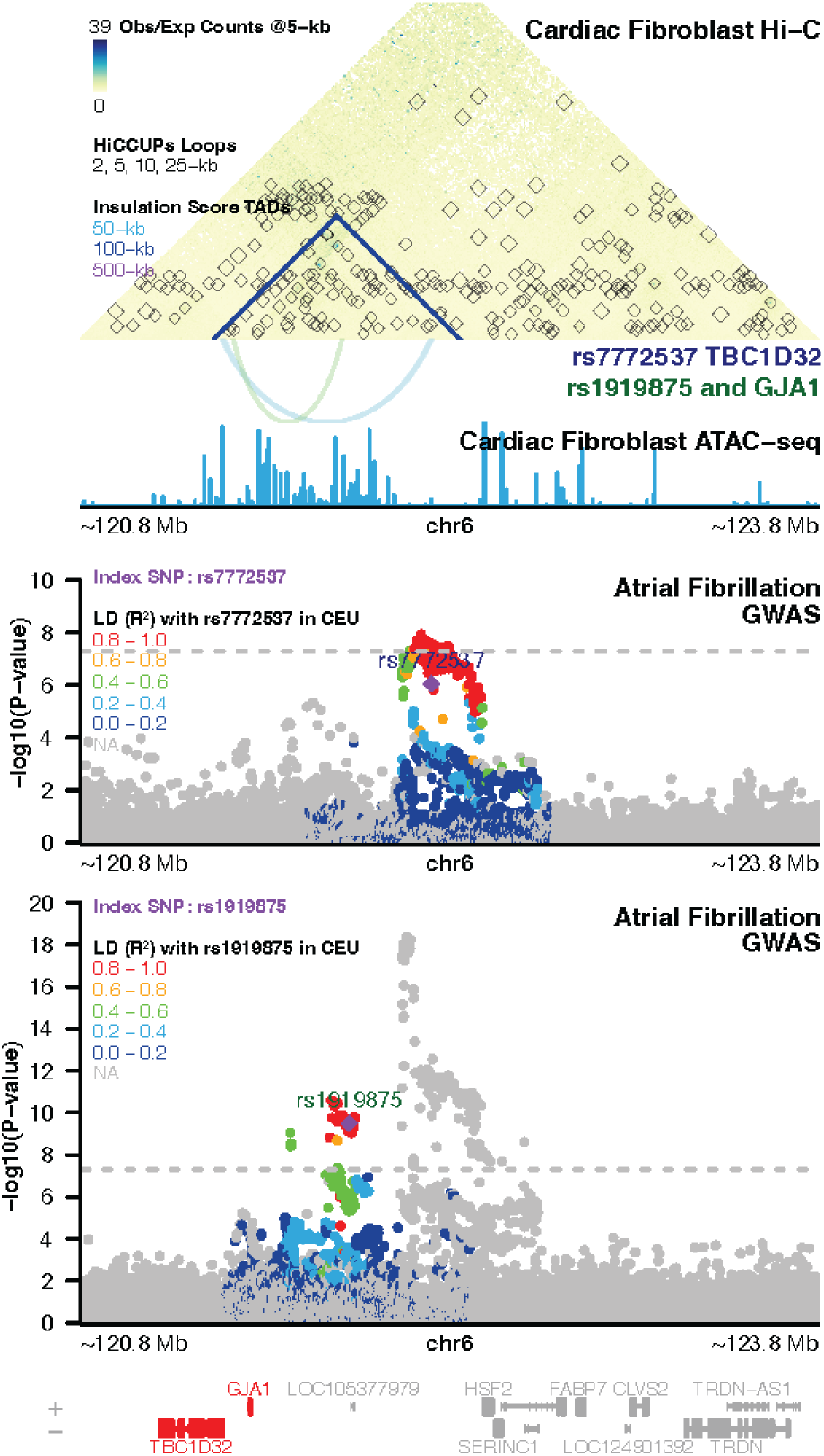
Significant interactions between atrial fibrillation-associated open chromatin at rs7772537 with *TBC1D32* and rs1919875 with *GJA1* in cardiac fibroblasts, overlaid with **A)** Arrowhead, B) Insulation Score, and C) Directionality Index TAD calls

**Figure S6:**
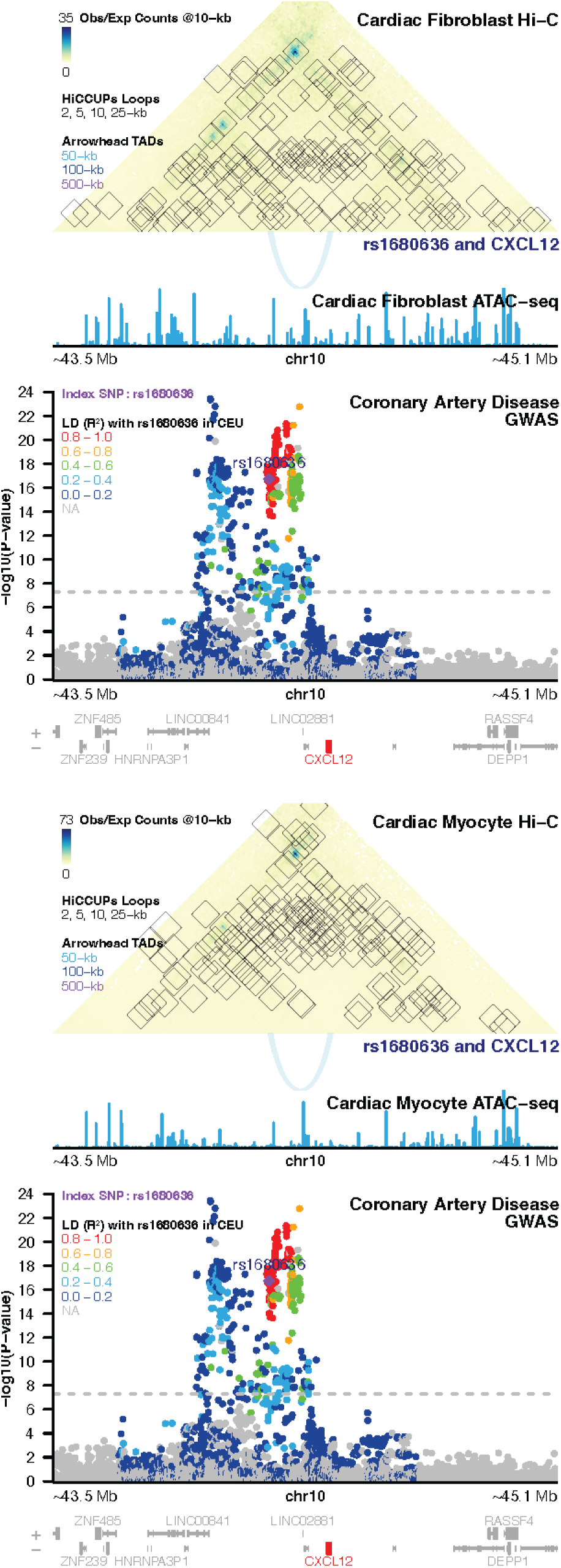
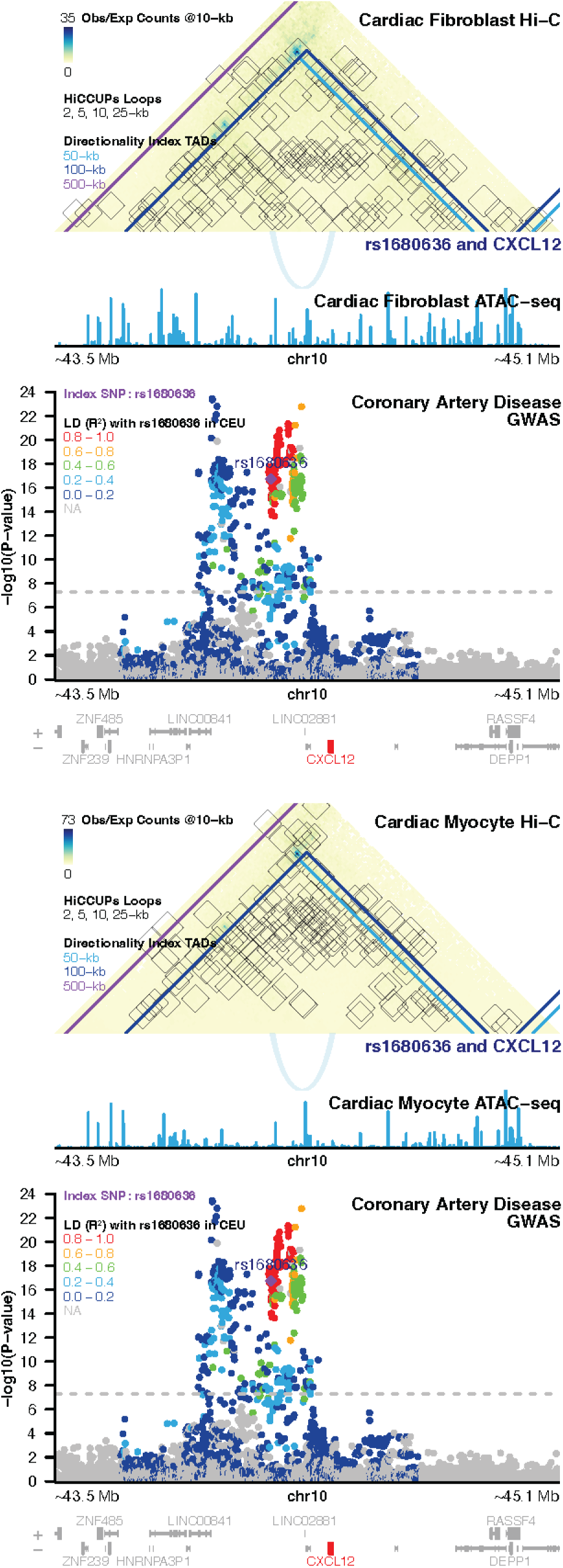
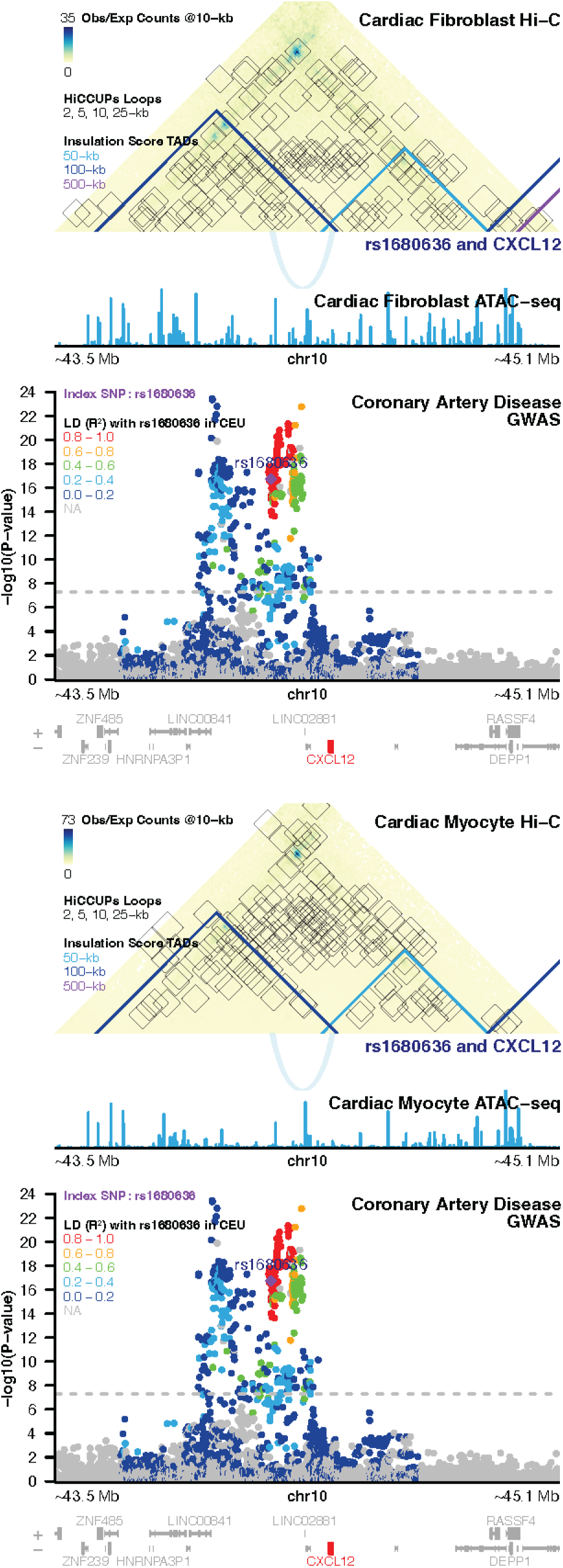
Significant interaction between coronary artery disease-associated open chromatin at rs1680636 and *CXCL12* in both cardiac fibroblasts and myocytes, overlaid with **A-B)** Arrowhead, **C-D)** Insulation Score, and **E-F)** Directionality Index TAD calls

**Figure S7:**
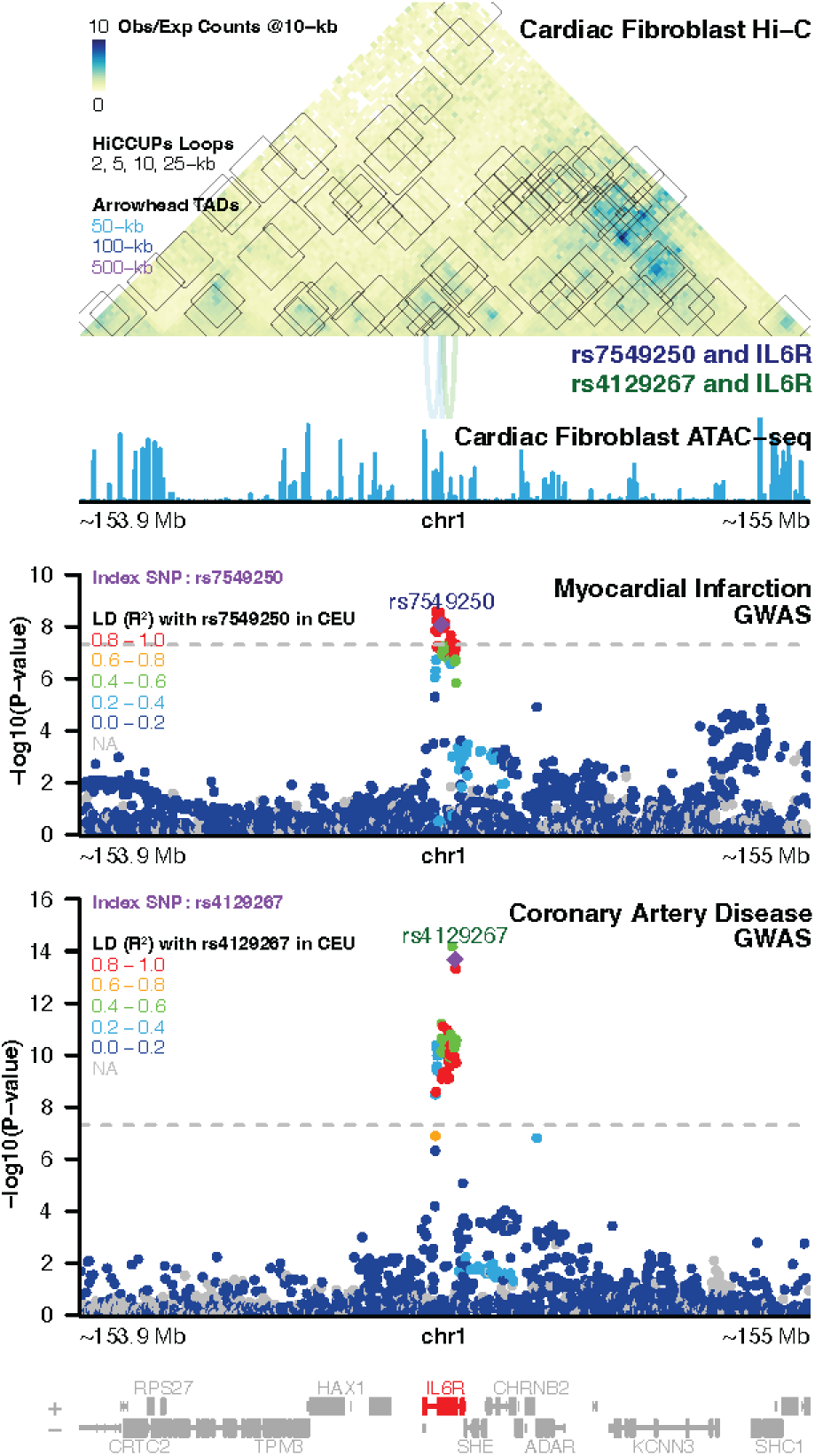
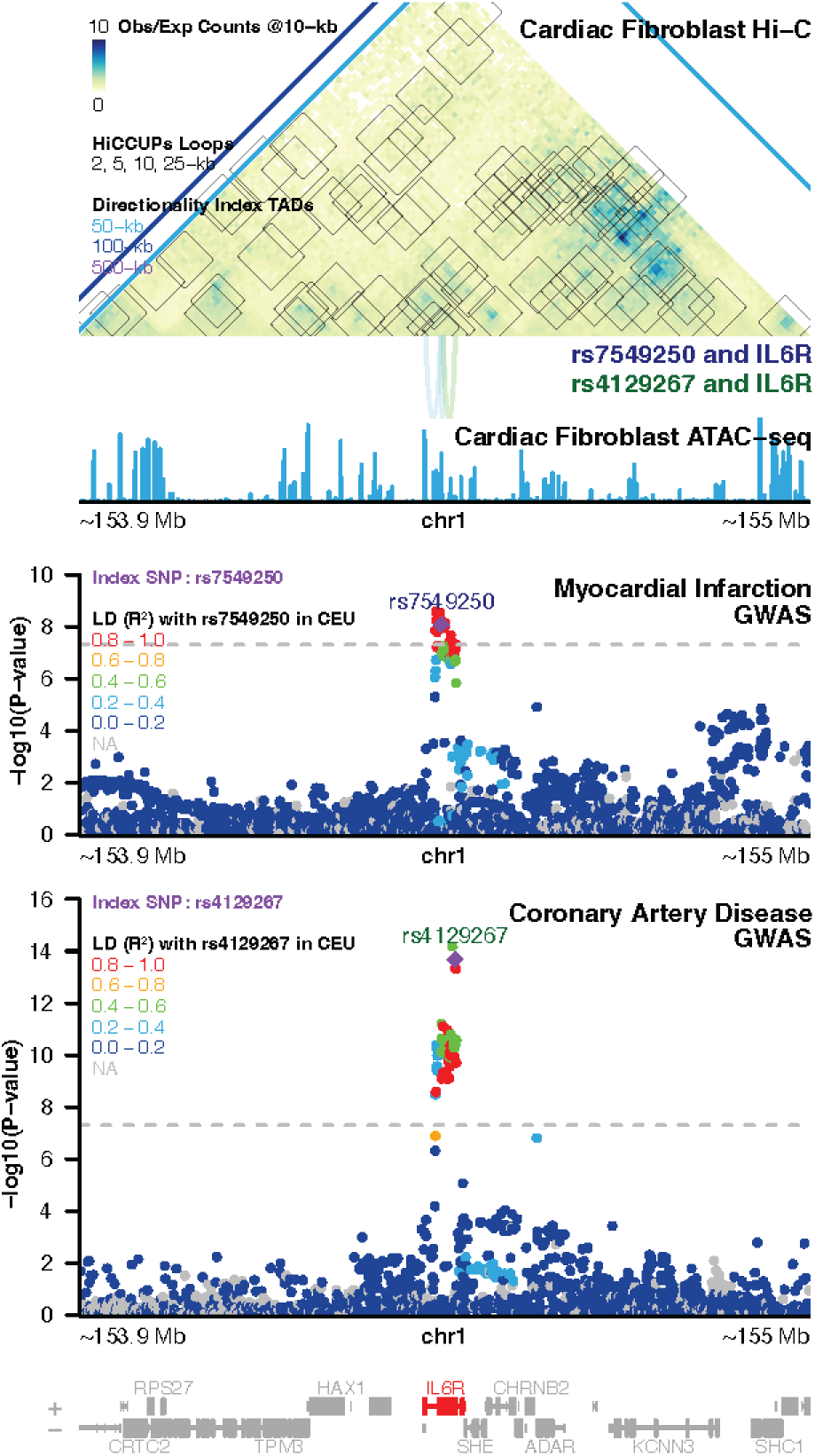
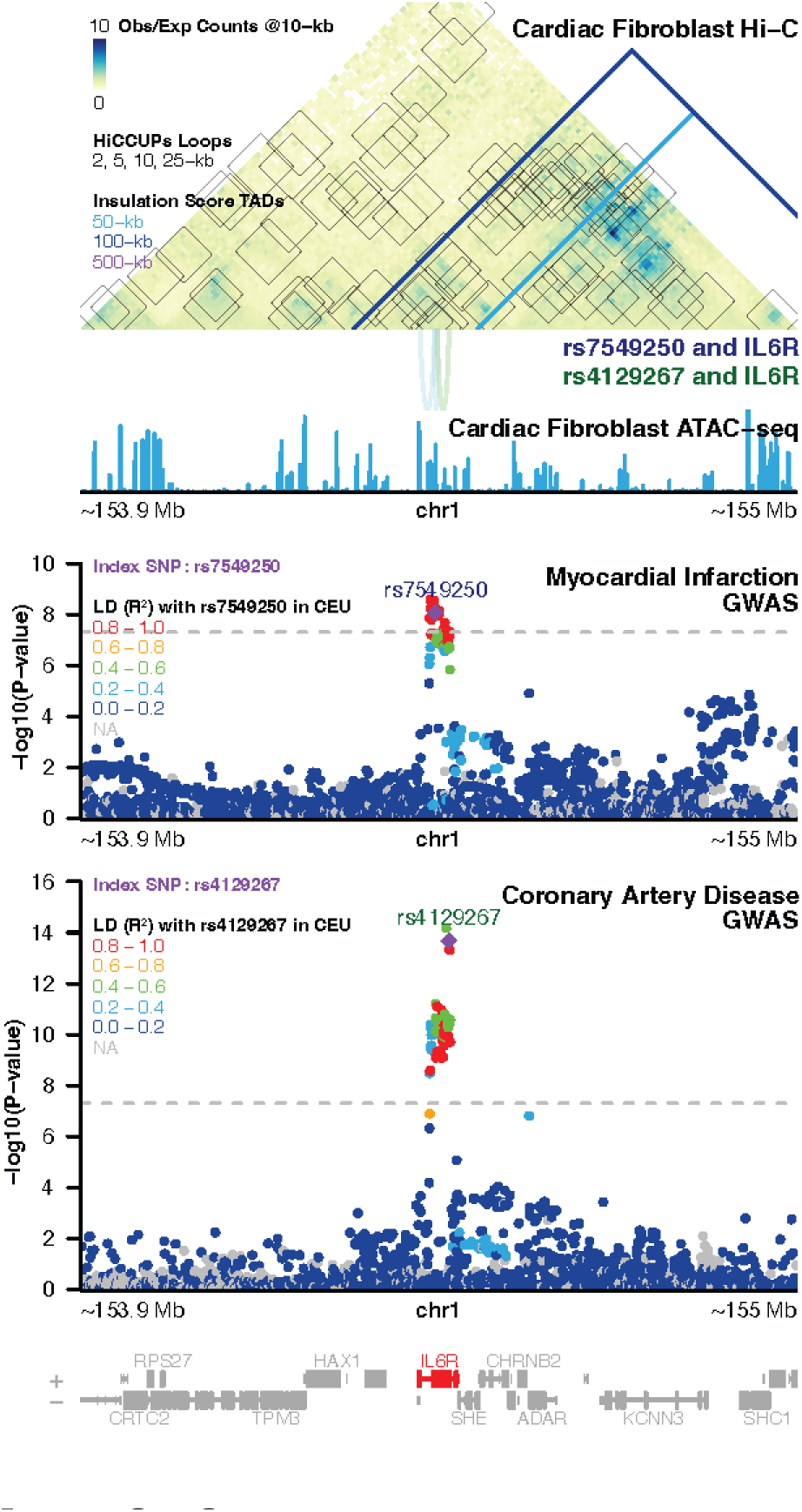
Significant interactions between myocardial infarction- and coronary artery disease-associated open chromatin at rs7549250 and rs4129267 with *IL6R* in cardiac fibroblasts, overlaid with **a)** Arrowhead, **b)** Directionality Index, and **c)** Insulation Score TAD calls

**Figure S8:**
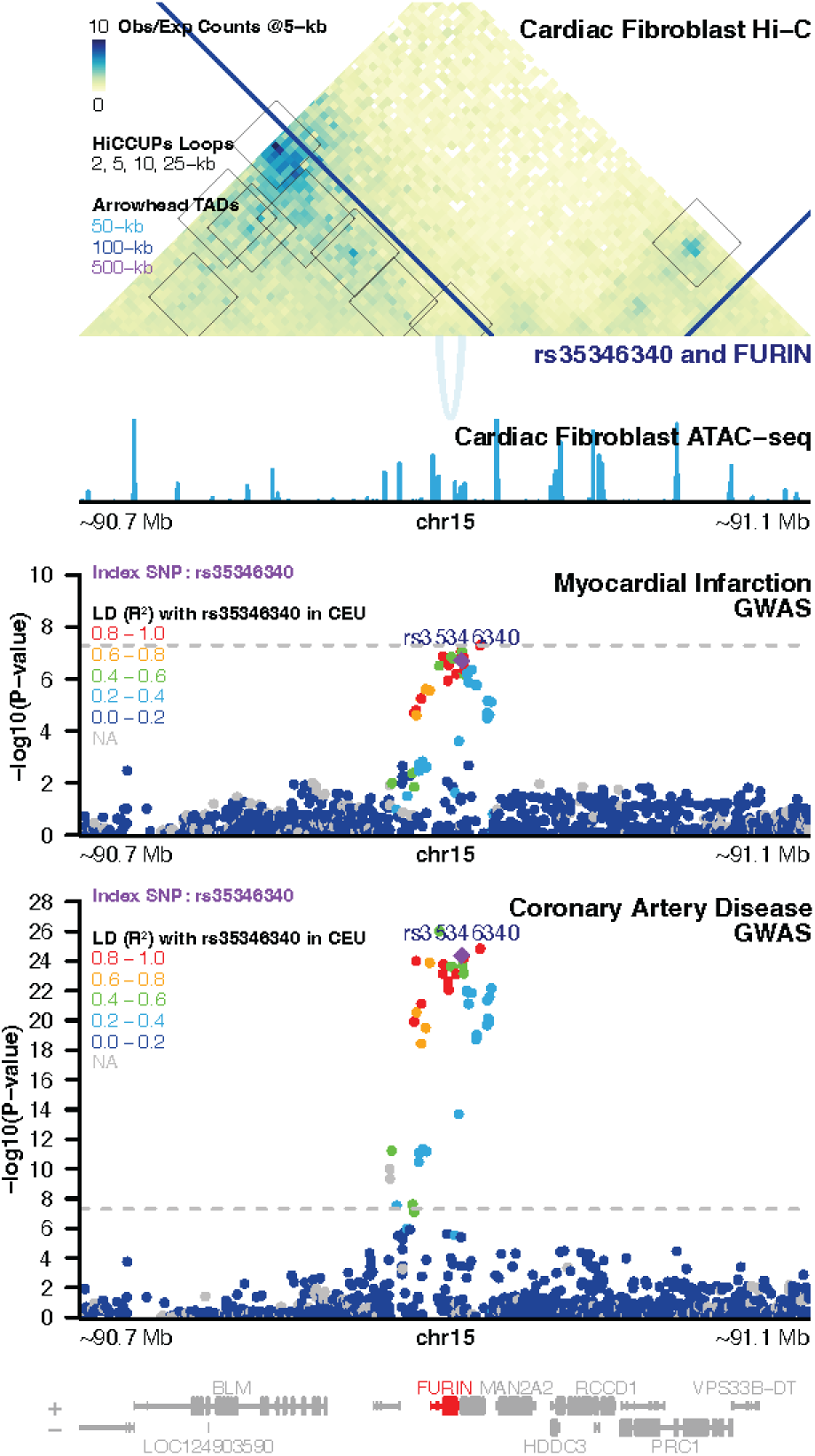
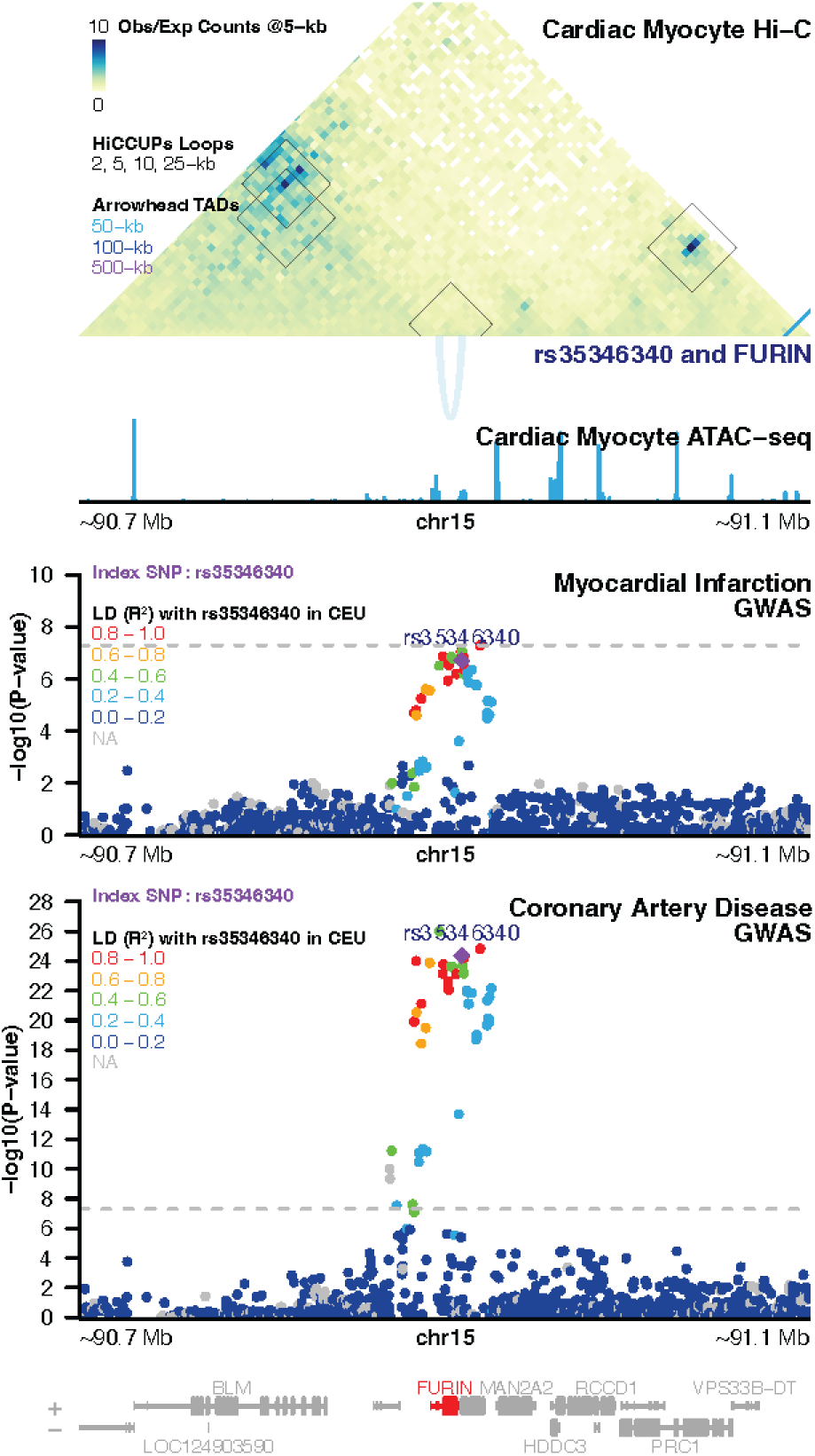
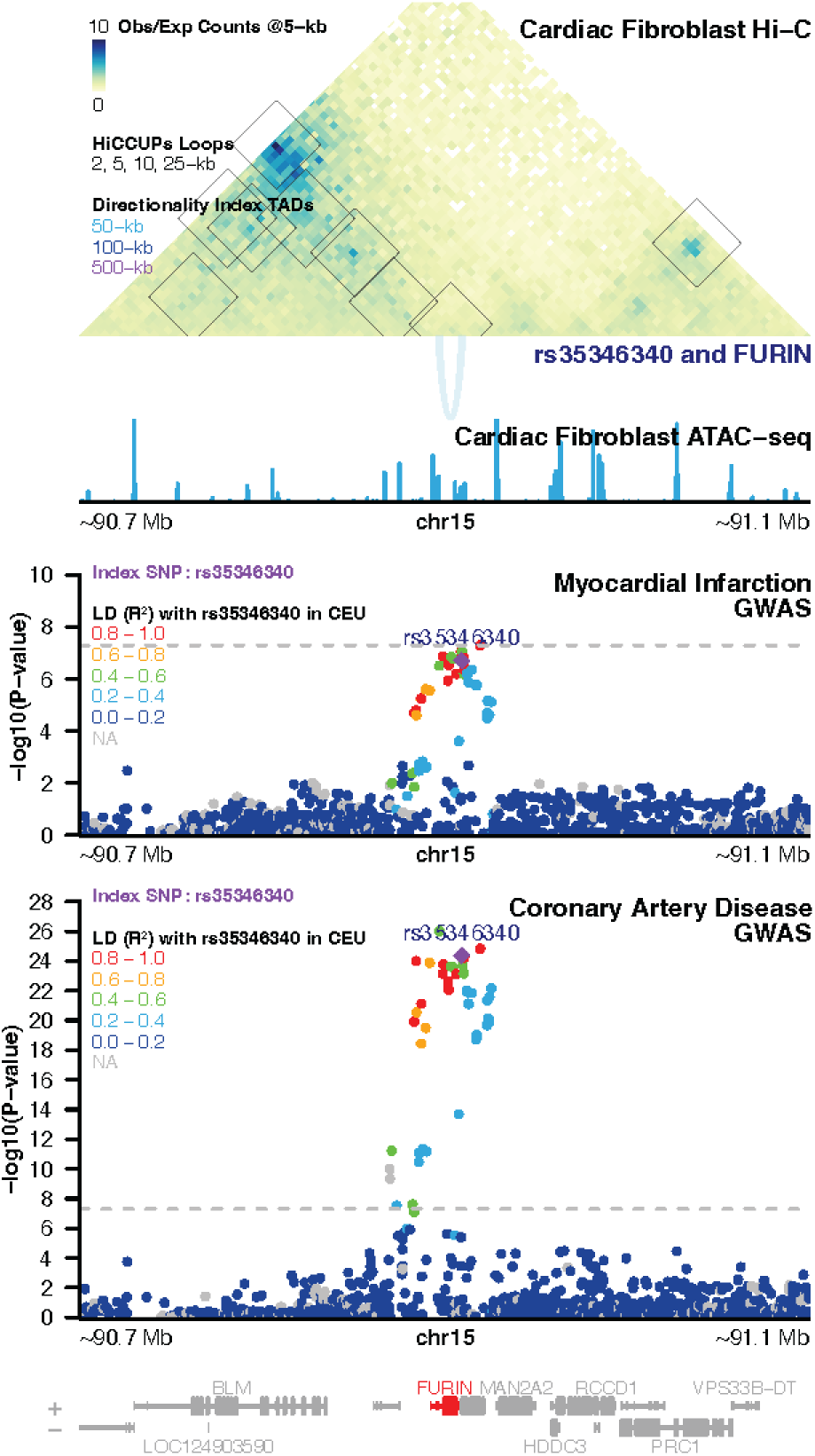
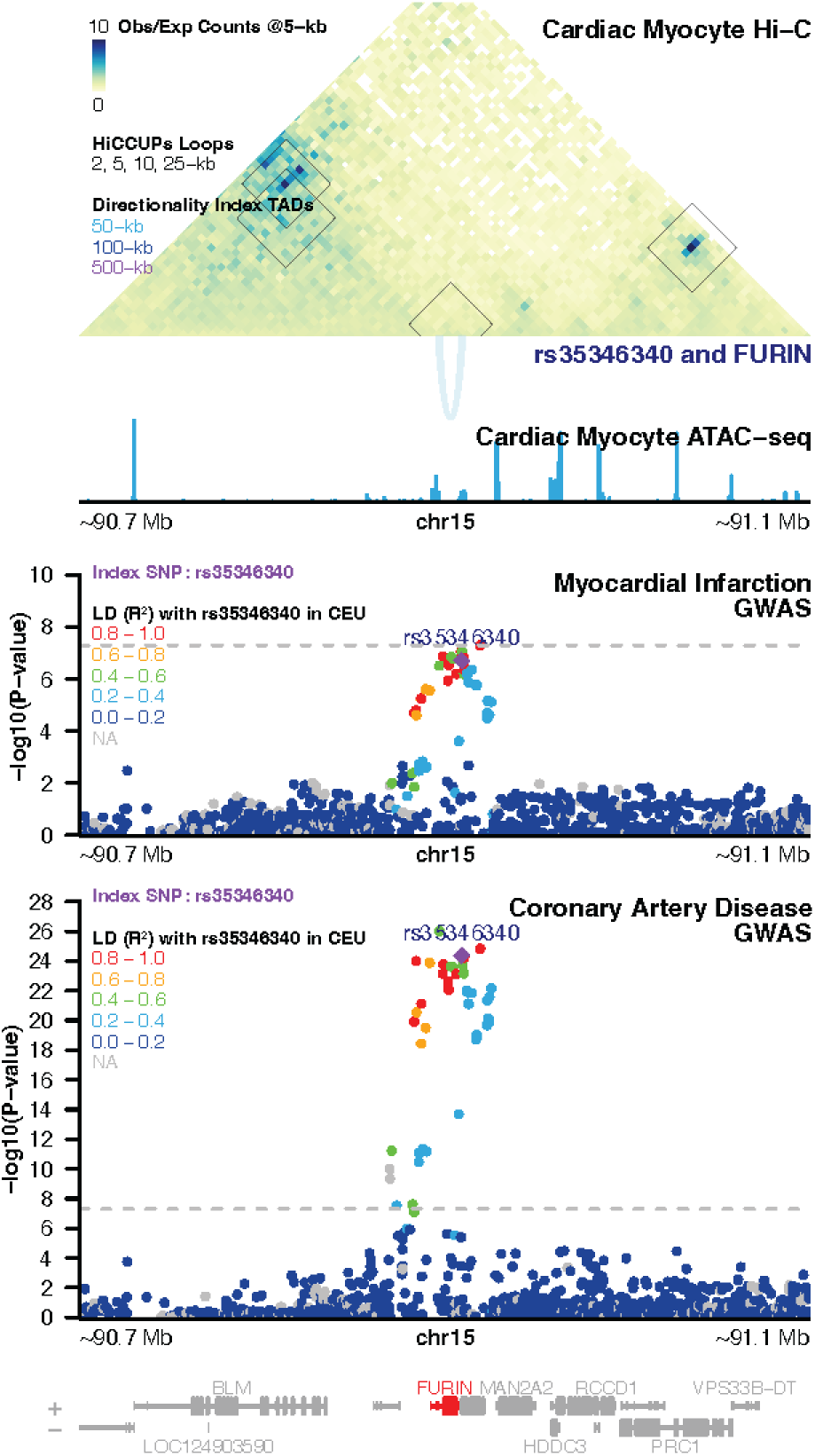
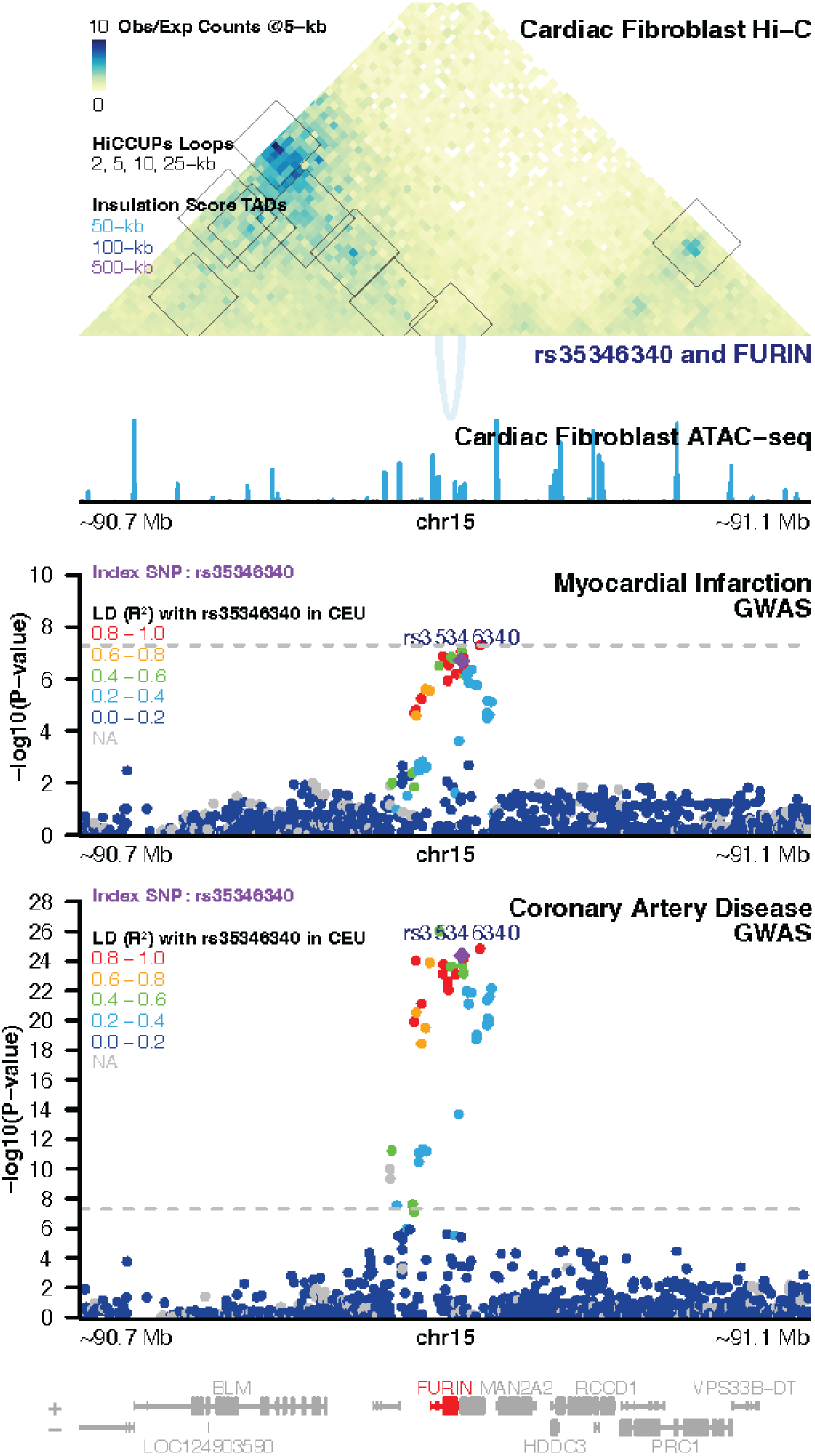
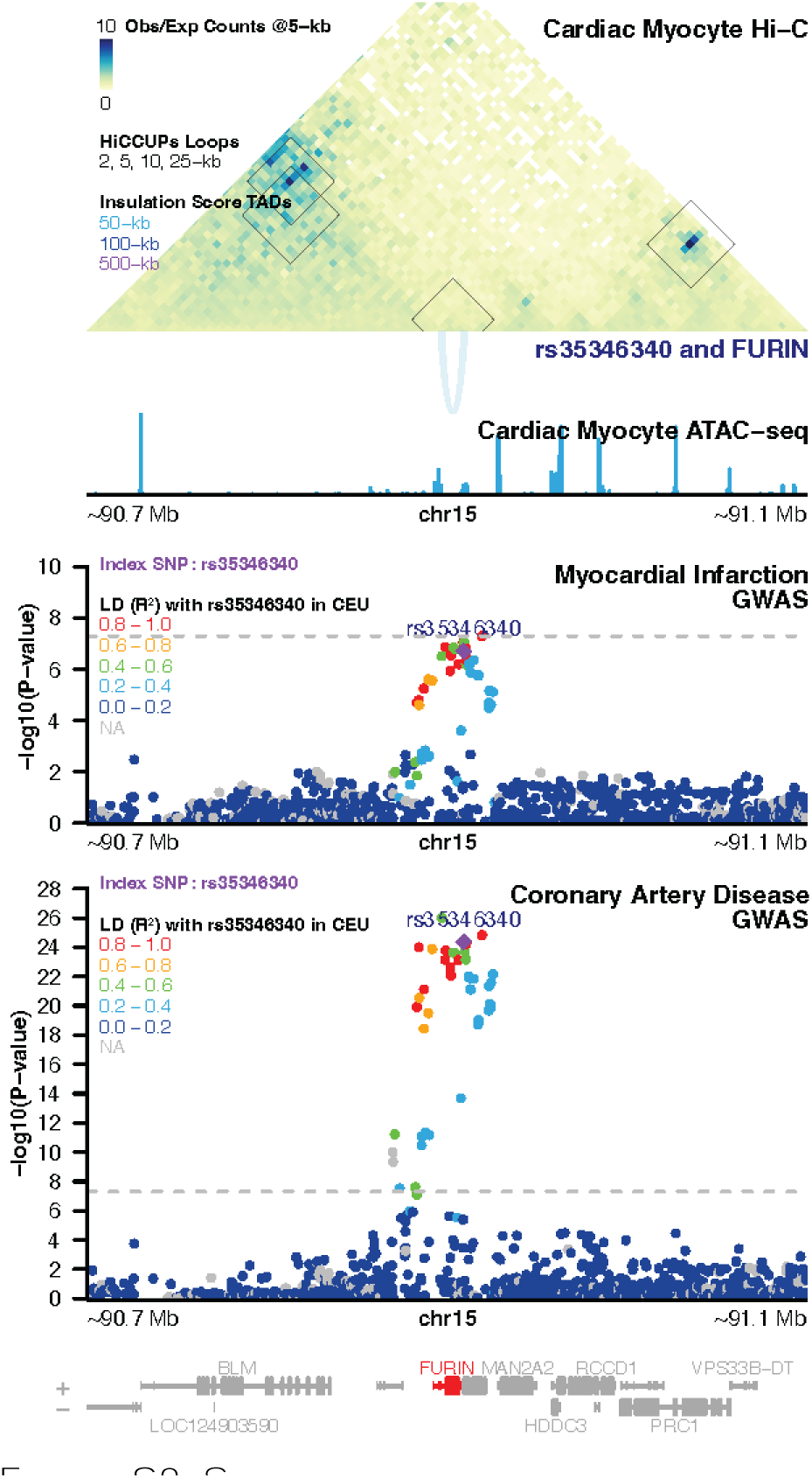
Significant interaction between myocardial infarction- and coronary artery disease-associated open chromatin at rs35346340 with *FURIN* in both cardiac fibroblasts and cardiac myocytes, overlaid with **A-B)** Arrowhead, **C-D)** Directionality Index, and **E-F)** Insulation Score TAD Calls

**Figure S9.**
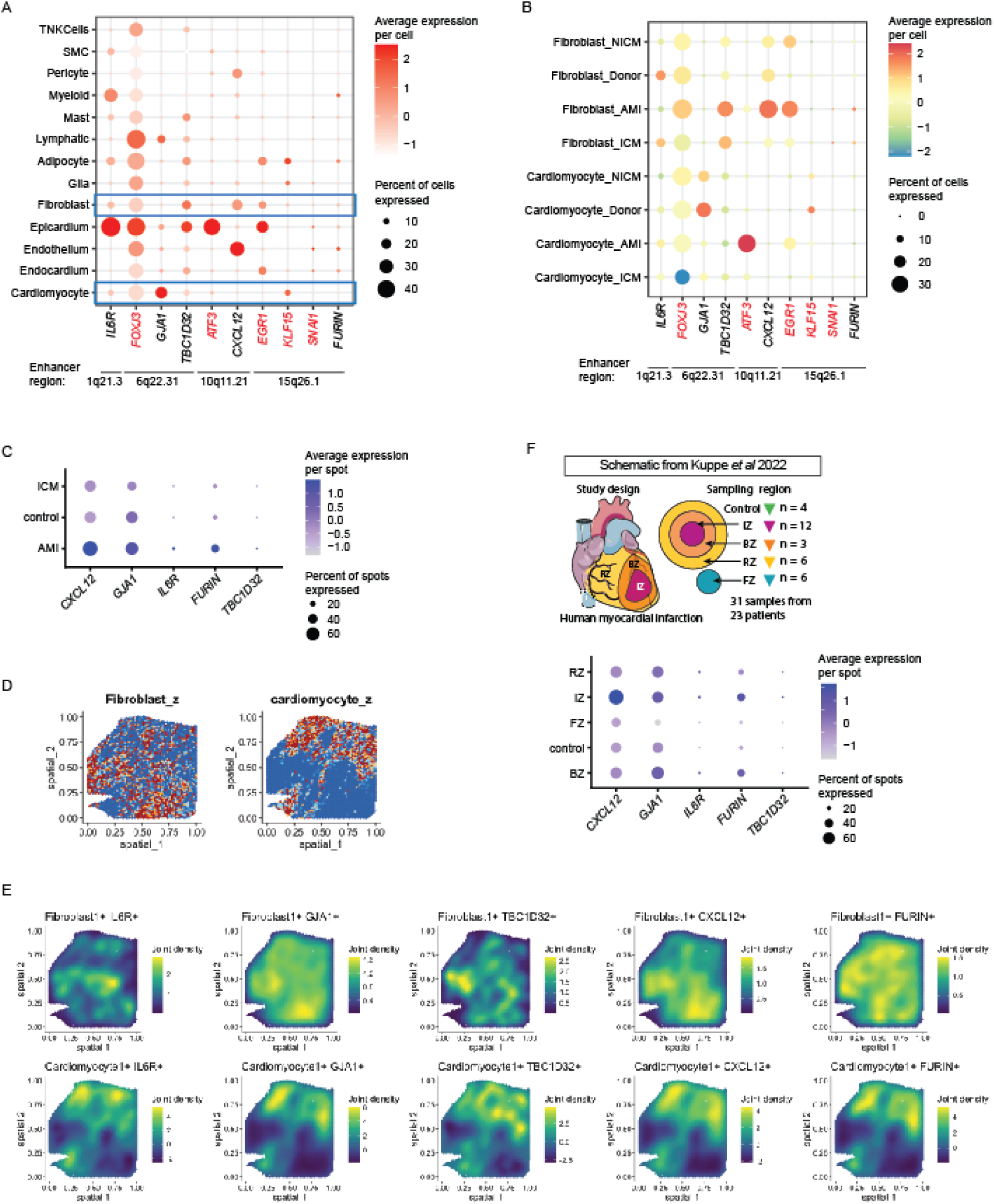
Expression of identified disease-associated target genes and transcription factors in heart disease single-nuclear RNA-seq from Amrute *et al* 2024 (a-b) and spatial transcriptomic data from Kuppe *et al* 2022 (c-f). **A)** Dot plot by cell type showing expression of relevant transcription factors (labeled in red) and target genes in human heart single-nuclear RNA-seq. **B)** Dot plot of heart fibroblasts and cardiomyocytes showing differential expression of transcription factors (labeled in red) and target genes in non-ischemic cardiomyopathy (NICM), non-heart disease (Donor), ischemic cardiomyopathy (ICM), and acute myocardial infarction (AMI). In **A)** and **B)**, the chromosome band of the enhancer region from which transcription factors and target genes were derived is depicted next to each gene. **C)** Dot plot of spatial transcriptomic Visium data showing target gene expression between 31 ICM, non-cardiomyopathy (control), and AMI heart sections. **D)** Spatial map of heart tissue section from donor with myocardial infarction, showing cell2location regions predicted to have fibroblasts and cardiomyocytes. **E)** Spatial co-expression of fibroblast/cardiomyocyte and target gene expression in heart section from the same donor as **D)**. **F)** Dot plot from spatial transcriptomic data showing target gene expression distribution across the remote zone (RZ), ischemic zone (IZ), fibrotic zone (FZ), and border zone (BZ) in heart sections from donors with myocardial infarction. Plot also includes expression from donors without cardiomyopathies (control).

**Figure S10:**
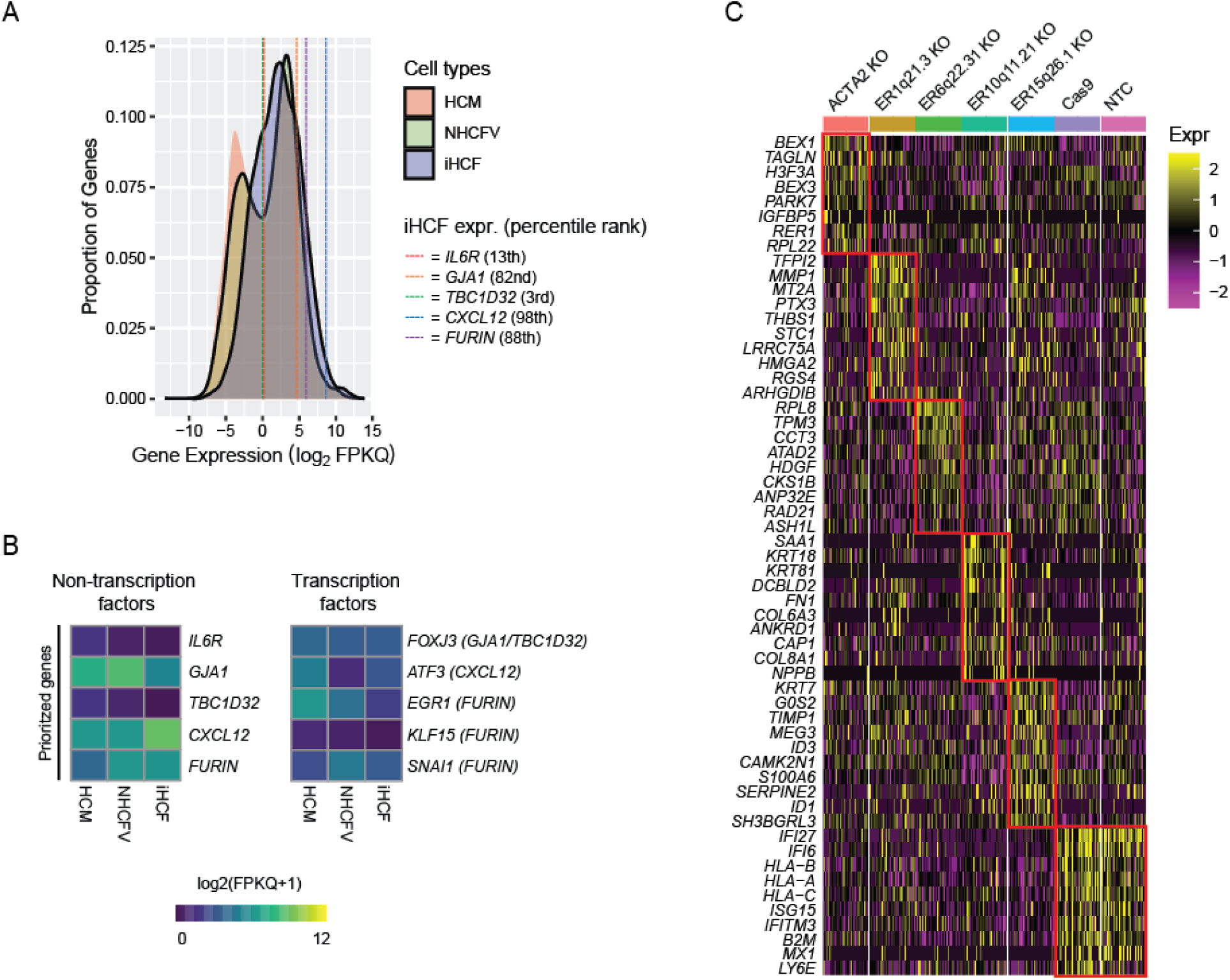
Analysis of transcriptional consequences following enhancer perturbation. **A)** Density plot of RNA-seq expression for expressed genes (FPKQ>0) in HCM, NHCFV, and iHCF cells. Target gene expression levels of iHCF are shown as dotted lines on the density plot. Percentile rank of each gene in iHCF compared to all expressed genes is shown next to gene labels. **B)** Heatmap of RNA-seq expression levels (log2(FPKQ+1) of prioritized target genes from each of the cell types used in this study. **C)** Heatmap showing the top highly expressed genes unique to each perturbation group. Each perturbation group was downsampled to 1,000 cells for visualization, and expression values are Z-scaled.

## SUPPLEMENTARY TABLES

**Table S1:** Statistics from processing 2-kb HCM and NHCFV data using HiC-Pro and Juicer

|  | HCM | NHCFV | HCM | NHCFV |
| --- | --- | --- | --- | --- |
|  | HiC-Pro |  | Juicer |  |
| **Deduplicated Valid Pairs** | 1,458,270,903 | 1,432,004,866 | 1,492,218,605 | 1,290,852,001 |
| **Duplicate Fraction** | 26.1% | 17.5% | 16.3% | 20.0% |
| **Trans interactions** | 208,202,698<br>(14.3%) | 224,662,049<br>(15.7%) | 167,702,055<br>(11.2%) | 165,994,213<br>(12.9%) |
| **Cis Interactions** | 1,250,068,205<br>(85.7%) | 1,207,342,817<br>(84.3%) | 1,324,516,550<br>(88.8%) | 1,124,857,788<br>(87.1%) |
| **Cis short-range (<20-kb)** | 532,593,173<br>(36.5%) | 255,375,571<br>(17.8%) | 716,530,612<br>(48.0%) | 355,404,192<br>(27.5%) |
| **Cis long-range (≥20-kb)** | 717,475,032<br>(49.2%) | 951,967,246<br>(66.5%) | 607,968,270<br>(40.7%) | 769,451,458<br>(59.6%) |
| **Resolution** |  |  | 1.9-kb | 2.3-kb |

HCM: Human cardiac myocytes; NHCFV: Normal human cardiac fibroblasts – ventricular

**Table S2:** Statistics from processing HCM, NHCFV, and iHCF ATAC-seq data using the ENCODE pipeline and HINT

|  | HCM |  | NHCFV | iHCF |
| --- | --- | --- | --- | --- |
|  | Replicate 1 | Replicate 2 |  |  |
| **Total aligned paired-end reads** | 154,396,643 | 155,600,836 | 214,927,706 | 208,495,502 |
| **Duplicate fraction** | 63.7% | 61.8% | 27.2% | 25.2% |
| **Retained reads** | 55,174,589<br>(25.1%) | 58,370,783<br>(26.6%) | 108,751,375<br>(50.4%) | 124,036,151<br>(59.3%) |
| **Non-redundant fraction** | 0.39 | 0.42 | 0.75 | 0.76 |
| **PCR Bottlenecking Coefficient 1** | 0.42 | 0.44 | 0.75 | 0.75 |
| **PCR Bottlenecking Coefficient 2** | 2.11 | 2.08 | 3.95 | 3.74 |
| **Peaks** | (IDR) 181,450 |  | (Naïve overlap)<br>284,571 | (Naïve overlap)<br>251,196 |
| **Footprinted regions** | 321,166 |  | 534,243 | 417,446 |
| **HOCOMOCO TF footprints** | 1,808,531 |  | 2,667,273 | 2,421,363 |
| **JASPAR TF footprints** | 866,389 |  | 1,373,866 | 1,320,785 |

HCM: Human cardiac myocytes; NHCFV: Normal human cardiac fibroblasts – ventricular; iHCF: Immortalized human cardiac fibroblasts

**Table S3:** Expression levels of expressed (FPKQ>0) genes in HCM, NHCFV, and iHCF cells

|  | HCM | NHCFV | iHCF |
| --- | --- | --- | --- |
| **Median FPKQ** | 1.4 | 2.5 | 4.4 |
| **Mean FPKQ** | 20.8 | 24.4 | 49.5 |
| **Genes** | 22,950 | 20,447 | 16,019 |

HCM: Human cardiac myocytes; NHCFV: Normal human cardiac fibroblasts – ventricular; iHCF: Immortalized human cardiac fibroblasts; FPKQ: Fragments per kilobase per million sequenced, quantile-normalized

**Table S4:**
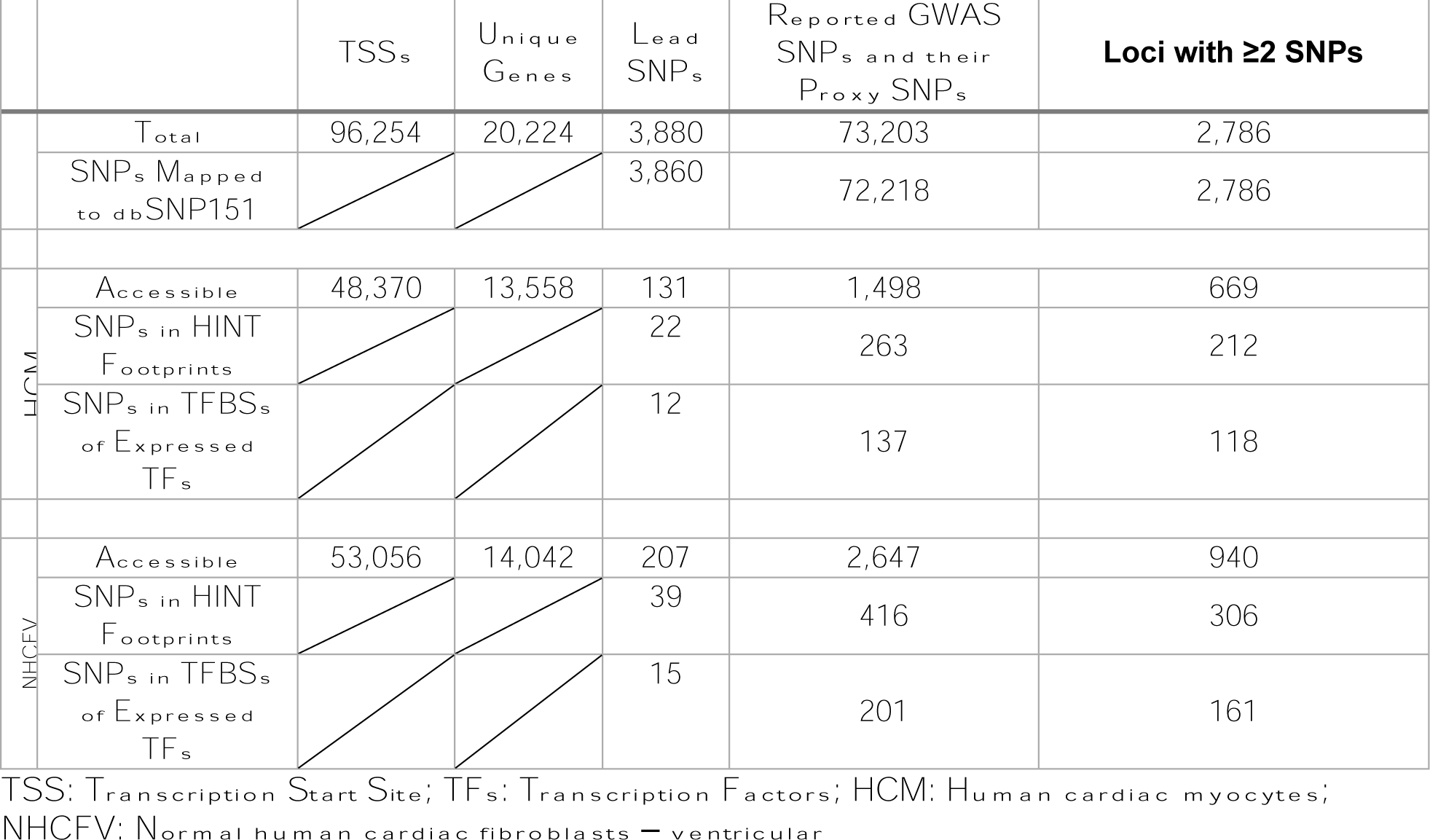
FANTOM5 TSSs and heart disease-associated SNPs and loci used to annotate chromatin loops

**Table S5:** Number and proportion of heart disease-related SNPs, TSSs, and their corresponding genes, with or without ATAC-based accessibility, mapping to significant interactions identified by GOTHiC and HiCCUPS. Proportions are relative to the totals in Table S4.

|  | **HCM** |  |  | **NHCFV** |  |  |
| --- | --- | --- | --- | --- | --- | --- |
|  | GOTHIC | HiCCUPS | Total | GOTHIC | HiCCUPS | Total |
| **SNPs Mapping to Significant Interactions** | 72,116<br>(99.9%) | 55,655<br>(77.1%) | 127,771 | 71,037<br>(98.4%) | 58,294<br>(80.7%) | 129,331 |
| **Accessible SNPs Mapping to Significant Interactions** | 1,497<br>(99.9%) | 1,319<br>(88.1%) | 2,816 | 2,606<br>(98.5%) | 2,182<br>(82.4%) | 4,788 |
| **TSSs Mapping to Significant Interactions** | 96,124<br>(99.9%) | 80,291<br>(83.4%) | 176,415 | 94,374<br>(98.0%) | 78,883<br>(82.0%) | 173,257 |
| **Accessible TSSs Mapping to Significant Interactions** | 48,335<br>(99.9%) | 43,459<br>(89.8%) | 91,794 | 52,053<br>(98.1%) | 44,630<br>(84.1%) | 96,683 |
| **Genes with TSSs Mapping to Significant Interactions** | 20,183<br>(99.8%) | 17,381<br>(85.9%) | 37,564 | 19,981<br>(98.8%) | 17,267<br>(85.4%) | 37,248 |
| **Genes with Accessible TSSs Mapping to Significant Interactions** | 13,540<br>(99.9%) | 12,118<br>(89.4%) | 25,658 | 13,829<br>(98.5%) | 11,950<br>(85.1%) | 25,779 |

HCM: Human cardiac myocytes; NHCFV: Normal human cardiac fibroblasts – ventricular

**Table S6A:** Number and attributes of significant interactions identified by GOTHiC and HiCCUPs in HCM and NHCFV cells

|  | HCM |  |  | NHCFV |  |  |
| --- | --- | --- | --- | --- | --- | --- |
|  | Combined | GOTHic | HiCCUPS | Combined | GOTHic | HiCCUPS |
| **Trans Interactions** | 2 | 2 | - | - | - | - |
| **Cis Interactions** | 2,927 | 2,224 | 703 | 2,149 | 1,437 | 712 |
| **Median Distance (Range)** | 78-kb<br>(2-kb to 134-Mb) | 52-kb<br>(2-kb to 134-kb) | 700-kb<br>(35-kb to 38-Mb) | 46-kb<br>(2-kb to 25-Mb) | 20-kb<br>(2-kb to 850-kb) | 600-kb<br>(35-kb to 25-Mb) |
| **Cis Short-Range (<20-kb)** | 541<br>(18.5%) | 541<br>(24.3%) | - | 718<br>(33.4%) | 718<br>(50%) | 1<br>(0.1%) |
| **Cis Long-Range (≥20-kb)** | 2,388<br>(81.6%) | 1,685<br>(75.8%) | 703<br>(100%) | 1,431<br>(66.6%) | 719<br>(50%) | 712<br>(100%) |

HCM: Human cardiac myocytes; NHCFV: Normal human cardiac fibroblasts – ventricular

**Table S6B:** Number and attributes of accessible SNP-gene pairs

|  |  | HCM |  |  | NHCFV |  |  |
| --- | --- | --- | --- | --- | --- | --- | --- |
|  |  | Combined | GOTHic | HiCCUPS | Combined | GOTHic | HiCCUPS |
| **SNP-Gene Pairs** | Active and Functional SNP-Gene Pairs | 3,982 | 2,852 | 1,384 | 3,275 | 1,882 | 1,478 |
|  | SNPs with Novel Predicted Target Genes | 3,001<br>(75.4%) | 1,887<br>(66.2%) | 1,299<br>(93.9%) | 2,362<br>(72.1%) | 1,022<br>(54.3%) | 1,393<br>(94.2%) |
|  | GTEX Heart (LV or RA) eQTLs | 78 (2.0%) | 77 (2.7%) | 10 (0.7%) | 61 (1.9%) | 58 (3.1%) | 4 (0.3%) |
|  | eQTLs in Any GTEX Tissue | 177<br>(4.4%) | 170<br>(6.0%) | 27 (2.0%) | 133<br>(4.1%) | 116<br>(6.2%) | 20 (1.4%) |
| **SNPs** | Unique SNPs | 1,275 | 1,128 | 744 | 1,531 | 1,155 | 858 |
|  | Underlying LD Clusters | 107 | 100 | 64 | 98 | 71 | 63 |
| **Genes** | Unique Genes | 1,878 | 1,342 | 787 | 1,490 | 830 | 781 |

HCM: Human cardiac myocytes; NHCFV: Normal human cardiac fibroblasts – ventricular; LV: left ventricle; RA: right atrium

**Table S7:** Number of TADs inferred across different algorithms and resolutions, and proportion of identified SNP-gene pairs falling into the same TAD per parameter choice.

| Cell Type,<br>Resolution | Arrowhead |  | Insulation Score |  | Directionality Index |  | GRiNCH |  |
| --- | --- | --- | --- | --- | --- | --- | --- | --- |
|  | Total TADs | % Same TAD | Total TADs | % Same TAD | Total TADs | % Same TAD | Total TADs | % Same TAD |
| NHCFV, 5 Kb | 8,476 | 39% | 4,479 | 13% | 8,203 | 19% | 2,519 | 88% |
| NHCFV, 10 Kb | 8,124 | 57% | 2,711 | 19% | 3,865 | 28% | 2,318 | 88% |
| NHCFV, 25 Kb | 4,406 | 63% | 1,518 | 21% | 1,649 | 38% | 1,954 | 88% |
| NHCFV, 50 Kb | 2,148 | 57% | 857 | 13% | 850 | 31% | 1,857 | 87% |
| NHCFV, 100 Kb | 914 | 45% | 470 | 9% | 428 | 38% | 1,511 | 89% |
| NHCFV, 500 Kb | 53 | 11% | 89 | 5% | 81 | 26% | 1,266 | 86% |
| HCM, 5 Kb | 8,688 | 27% | 4,058 | 5% | 8,634 | 19% | 2,645 | 91% |
| HCM, 10 Kb | 8,157 | 48% | 2,556 | 11% | 4,053 | 26% | 2,499 | 91% |
| HCM, 25 Kb | 4,400 | 49% | 1,563 | 14% | 1,670 | 41% | 1,976 | 91% |
| HCM, 50 Kb | 2,109 | 44% | 878 | 13% | 846 | 32% | 1,659 | 92% |
| HCM, 100 Kb | 862 | 40% | 443 | 6% | 409 | 36% | 1,489 | 92% |
| HCM, 500 Kb | 45 | 13% | 89 | 3% | 73 | 22% | 803 | 95% |

HCM: Human cardiac myocytes; NHCFV: Normal human cardiac fibroblasts – ventricular

**Table S7:** KEGG pathways enriched in differentially expressed genes from CRISPR knockout experiments

| Enhancer | Predicted Target Genes | Heart Failure-Relevant KEGG Pathway | Core Enrichment Genes | Normalized Enrichment Score (NES) | Adjusted *P*-value |
| --- | --- | --- | --- | --- | --- |
| **C1756** | *GJA1*,<br>*TBC1D32* | Cardiac muscle contraction | *COX7A1* | 2.41 | 2.39E-09 |
|  |  | Diabetic cardiomyopathy |  | 2.35 | 2.39E-09 |
|  |  | Vascular smooth muscle contraction |  | -2.10 | 2.39E-09 |
| **C1758** | *FURIN* | Dilated cardiomyopathy | *ACTG1* | 2.00 | 5.96E-03 |
|  |  | Hypertrophic cardiomyopathy |  | 1.84 | 2.26E-02 |

## Bibliography

1. Go, A.S., et al., Executive summary: heart disease and stroke statistics--2013 update: a report from the American Heart Association. Circulation, 2013. 127(1): p. 143–52.

2. Ziaeian, B. and G.C. Fonarow, Epidemiology and aetiology of heart failure. Nat Rev Cardiol, 2016. 13(6): p. 368–78.

3. Ahmad, F., J.G. Seidman, and C.E. Seidman, The genetic basis for cardiac remodeling. Annu Rev Genomics Hum Genet, 2005. 6: p. 185–216.

4. Dubois, C.M., et al., Evidence that furin is an authentic transforming growth factor-beta1-converting enzyme. Am J Pathol, 2001. 158(1): p. 305–16.

5. Nakajima, Y., et al., Expression of smooth muscle alpha-actin in mesenchymal cells during formation of avian endocardial cushion tissue: a role for transforming growth factor beta3. Dev Dyn, 1997. 209(3): p. 296–309.

6. Chen, M.M., et al., CTGF expression is induced by TGF-beta in cardiac fibroblasts and cardiac myocytes: a potential role in heart fibrosis. J Mol Cell Cardiol, 2000. 32(10): p. 1805–19.

7. Wang, B., et al., The Kruppel-like factor KLF15 inhibits connective tissue growth factor (CTGF) expression in cardiac fibroblasts. J Mol Cell Cardiol, 2008. 45(2): p. 193–7.

8. Campbell, S.E. and L.C. Katwa, Angiotensin II stimulated expression of transforming growth factor-beta1 in cardiac fibroblasts and myofibroblasts. J Mol Cell Cardiol, 1997. 29(7): p. 1947–58.

9. Leask, A., Getting to the heart of the matter: new insights into cardiac fibrosis. Circ Res, 2015. 116(7): p. 1269–76.

10. Martin, P., et al., Effect of a highly selective endothelin-converting enzyme inhibitor on cardiac remodeling in rats after myocardial infarction. J Cardiovasc Pharmacol, 2000. 36(5 Suppl 1): p. S367–70.

11. Porter, K.E. and N.A. Turner, Cardiac fibroblasts: at the heart of myocardial remodeling. Pharmacol Ther, 2009. 123(2): p. 255–78.

12. Kong, P., P. Christia, and N.G. Frangogiannis, The pathogenesis of cardiac fibrosis. Cell Mol Life Sci, 2014. 71(4): p. 549–74.

13. Jackson, E.K., et al., SDF-1alpha (Stromal Cell-Derived Factor 1alpha) Induces Cardiac Fibroblasts, Renal Microvascular Smooth Muscle Cells, and Glomerular Mesangial Cells to Proliferate, Cause Hypertrophy, and Produce Collagen. J Am Heart Assoc, 2017. 6(11).

14. Consortium, C.A.D., et al., Large-scale association analysis identifies new risk loci for coronary artery disease. Nat Genet, 2013. 45(1): p. 25–33.

15. Nikpay, M., et al., A comprehensive 1,000 Genomes-based genome-wide association meta-analysis of coronary artery disease. Nat Genet, 2015. 47(10): p. 1121–1130.

16. Webb, T.R., et al., Systematic Evaluation of Pleiotropy Identifies 6 Further Loci Associated With Coronary Artery Disease. J Am Coll Cardiol, 2017. 69(7): p. 823–836.

17. van der Harst, P. and N. Verweij, Identification of 64 Novel Genetic Loci Provides an Expanded View on the Genetic Architecture of Coronary Artery Disease. Circ Res, 2018. 122(3): p. 433–443.

18. Christophersen, I.E., et al., Large-scale analyses of common and rare variants identify 12 new loci associated with atrial fibrillation. Nat Genet, 2017. 49(6): p. 946–952.

19. Ellinor, P.T., et al., Meta-analysis identifies six new susceptibility loci for atrial fibrillation. Nat Genet, 2012. 44(6): p. 670–5.

20. Dichgans, M., et al., Shared genetic susceptibility to ischemic stroke and coronary artery disease: a genome-wide analysis of common variants. Stroke, 2014. 45(1): p. 24–36.

21. Aulchenko, Y.S., et al., Loci influencing lipid levels and coronary heart disease risk in 16 European population cohorts. Nat Genet, 2009. 41(1): p. 47–55.

22. Nelson, C.P., et al., Association analyses based on false discovery rate implicate new loci for coronary artery disease. Nat Genet, 2017. 49(9): p. 1385–1391.

23. Schunkert, H., et al., Large-scale association analysis identifies 13 new susceptibility loci for coronary artery disease. Nat Genet, 2011. 43(4): p. 333–8.

24. Yeo, A., et al., Pharmacogenetic meta-analysis of baseline risk factors, pharmacodynamic, efficacy and tolerability endpoints from two large global cardiovascular outcomes trials for darapladib. PLoS One, 2017. 12(7): p. e0182115.

25. Elliott, P., et al., Genetic Loci associated with C-reactive protein levels and risk of coronary heart disease. JAMA, 2009. 302(1): p. 37–48.

26. Smith, N.L., et al., Association of genome-wide variation with the risk of incident heart failure in adults of European and African ancestry: a prospective meta-analysis from the cohorts for heart and aging research in genomic epidemiology (CHARGE) consortium. Circ Cardiovasc Genet, 2010. 3(3): p. 256–66.

27. Shah, S., et al., Genome-wide association and Mendelian randomisation analysis provide insights into the pathogenesis of heart failure. Nat Commun, 2020. 11(1): p. 163.

28. Nielsen, J.B., et al., Biobank-driven genomic discovery yields new insight into atrial fibrillation biology. Nat Genet, 2018. 50(9): p. 1234–1239.

29. Maurano, M.T., et al., Systematic localization of common disease-associated variation in regulatory DNA. Science, 2012. 337(6099): p. 1190–5.

30. Roadmap Epigenomics, C., et al., Integrative analysis of 111 reference human epigenomes. Nature, 2015. 518(7539): p. 317–30.

31. Umans, B.D., A. Battle, and Y. Gilad, Where Are the Disease-Associated eQTLs? Trends Genet, 2021. 37(2): p. 109–124.

32. Mostafavi, H., et al., Limited overlap of eQTLs and GWAS hits due to systematic differences in discovery. 2022: p. 2022.05.07.491045.

33. Musunuru, K., et al., From noncoding variant to phenotype via SORT1 at the 1p13 cholesterol locus. Nature, 2010. 466(7307): p. 714–9.

34. Smemo, S., et al., Obesity-associated variants within FTO form long-range functional connections with IRX3. Nature, 2014. 507(7492): p. 371–5.

35. Gupta, R.M., et al., A Genetic Variant Associated with Five Vascular Diseases Is a Distal Regulator of Endothelin-1 Gene Expression. Cell, 2017. 170(3): p. 522–533 e15.

36. Wright, J.B., S.J. Brown, and M.D. Cole, Upregulation of c-MYC in cis through a large chromatin loop linked to a cancer risk-associated single-nucleotide polymorphism in colorectal cancer cells. Mol Cell Biol, 2010. 30(6): p. 1411–20.

37. Cowper-Sal lari, R., et al., Breast cancer risk-associated SNPs modulate the affinity of chromatin for FOXA1 and alter gene expression. Nat Genet, 2012. 44(11): p. 1191–8.

38. Symmons, O., et al., Functional and topological characteristics of mammalian regulatory domains. Genome Res, 2014. 24(3): p. 390–400.

39. Sexton, T. and G. Cavalli, The role of chromosome domains in shaping the functional genome. Cell, 2015. 160(6): p. 1049–59.

40. Dixon, J.R., et al., Topological domains in mammalian genomes identified by analysis of chromatin interactions. Nature, 2012. 485(7398): p. 376–80.

41. Schmitt, A.D., et al., A Compendium of Chromatin Contact Maps Reveals Spatially Active Regions in the Human Genome. Cell Rep, 2016. 17(8): p. 2042–2059.

42. Rao, S.S., et al., A 3D map of the human genome at kilobase resolution reveals principles of chromatin looping. Cell, 2014. 159(7): p. 1665–80.

43. Mifsud, B., et al., Mapping long-range promoter contacts in human cells with high-resolution capture Hi-C. Nat Genet, 2015. 47(6): p. 598–606.

44. Cairns, J., et al., CHiCAGO: robust detection of DNA looping interactions in Capture Hi-C data. Genome Biology, 2016. 17(1): p. 127.

45. Choy, M.K., et al., Promoter interactome of human embryonic stem cell-derived cardiomyocytes connects GWAS regions to cardiac gene networks. Nat Commun, 2018. 9(1): p. 2526.

46. Montefiori, L.E., et al., A promoter interaction map for cardiovascular disease genetics. Elife, 2018. 7.

47. Brandao, K.O., et al., Human pluripotent stem cell models of cardiac disease: from mechanisms to therapies. Dis Model Mech, 2017. 10(9): p. 1039–1059.

48. Servant, N., et al., HiC-Pro: an optimized and flexible pipeline for Hi-C data processing. Genome Biol, 2015. 16: p. 259.

49. Durand, N.C., et al., Juicer Provides a One-Click System for Analyzing Loop-Resolution Hi-C Experiments. Cell Syst, 2016. 3(1): p. 95–8.

50. Mifsud, B., et al., GOTHiC, a probabilistic model to resolve complex biases and to identify real interactions in Hi-C data. PLoS One, 2017. 12(4): p. e0174744.

51. Forcato, M., et al., Comparison of computational methods for Hi-C data analysis. Nat Methods, 2017. 14(7): p. 679–685.

52. Schoenfelder, S., et al., The pluripotent regulatory circuitry connecting promoters to their long-range interacting elements. Genome Res, 2015. 25(4): p. 582–97.

53. Lee, D.-I. and S. Roy, GRiNCH: simultaneous smoothing and detection of topological units of genome organization from sparse chromatin contact count matrices with matrix factorization. Genome Biology, 2021. 22(1): p. 164.

54. Kruse, K., et al., TADtool: visual parameter identification for TAD-calling algorithms. Bioinformatics, 2016. 32(20): p. 3190–3192.

55. Quinlan, A.R. and I.M. Hall, BEDTools: a flexible suite of utilities for comparing genomic features. Bioinformatics, 2010. 26(6): p. 841–2.

56. Buenrostro, J.D., et al., ATAC-seq: A Method for Assaying Chromatin Accessibility Genome-Wide. Curr Protoc Mol Biol, 2015. 109: p. 21 29 1-9.

57. Lee, J.W., Chuan, S. F., Kim, D., Boley, N., Kundaje, A. ATAC-Seq / DNase-Seq Pipeline. 2017 [cited 2017 August 7]; Available from: https://github.com/kundajelab/atac_dnase_pipelines.

58. Gusmao, E.G., et al., Analysis of computational footprinting methods for DNase sequencing experiments. Nat Methods, 2016. 13(4): p. 303–9.

59. Gusmao, E.G., et al., Detection of active transcription factor binding sites with the combination of DNase hypersensitivity and histone modifications. Bioinformatics, 2014. 30(22): p. 3143–51.

60. Li, Z., et al., Identification of transcription factor binding sites using ATAC-seq. Genome Biol, 2019. 20(1): p. 45.

61. Li, Z., et al., RGT: a toolbox for the integrative analysis of high throughput regulatory genomics data. BMC Bioinformatics, 2023. 24(1): p. 79.

62. Kulakovskiy, I.V., et al., HOCOMOCO: towards a complete collection of transcription factor binding models for human and mouse via large-scale ChIP-Seq analysis. Nucleic Acids Res, 2018. 46(D1): p. D252–D259.

63. Khan, A., et al., JASPAR 2018: update of the open-access database of transcription factor binding profiles and its web framework. Nucleic Acids Res, 2018. 46(D1): p. D260–D266.

64. Sandelin, A., et al., JASPAR: an open-access database for eukaryotic transcription factor binding profiles. Nucleic Acids Res, 2004. 32(Database issue): p. D91–4.

65. Wilczynski, B., et al., Finding evolutionarily conserved cis-regulatory modules with a universal set of motifs. BMC Bioinformatics, 2009. 10: p. 82.

66. Parkhomchuk, D., et al., Transcriptome analysis by strand-specific sequencing of complementary DNA. Nucleic Acids Res, 2009. 37(18): p. e123.

67. Sultan, M., et al., A global view of gene activity and alternative splicing by deep sequencing of the human transcriptome. Science, 2008. 321(5891): p. 956–60.

68. MacArthur, J., et al., The new NHGRI-EBI Catalog of published genome-wide association studies (GWAS Catalog). Nucleic Acids Res, 2017. 45(D1): p. D896–D901.

69. Machiela, M.J. and S.J. Chanock, LDlink: a web-based application for exploring population-specific haplotype structure and linking correlated alleles of possible functional variants. Bioinformatics, 2015. 31(21): p. 3555–7.

70. Abugessaisa, I., et al., The FANTOM5 Computation Ecosystem: Genomic Information Hub for Promoters and Active Enhancers. Methods Mol Biol, 2017. 1611: p. 199–217.

71. Yates, B., et al., Genenames.org: the HGNC and VGNC resources in 2017. Nucleic Acids Res, 2017. 45(D1): p. D619–D625.

72. Carithers, L.J., et al., A Novel Approach to High-Quality Postmortem Tissue Procurement: The GTEx Project. Biopreserv Biobank, 2015. 13(5): p. 311–9.

73. Durand, N.C., et al., Juicebox Provides a Visualization System for Hi-C Contact Maps with Unlimited Zoom. Cell Syst, 2016. 3(1): p. 99–101.

74. Kerpedjiev P., A.N., Lekschas F., McCallum C., Dinkla K., Strobelt H., Luber J. M., Ouellette S., Azhir A., Kumar N., Hwang J., Lee S., Alver B. H., Pfister H., Mirny L. A., Park P. J., Gehlenborg N., HiGlass: Web-based Visual Exploration and Analysis of Genome Interaction Maps. bioRxiv, 121889.

75. Kramer, N.E., et al., Plotgardener: cultivating precise multi-panel figures in R. Bioinformatics, 2022. 38(7): p. 2042–2045.

76. Sigma, M. CRISPR Design Tools. 2021 [cited 2021; Available from: https://www.milliporesigmabioinfo.com/bioinfo_tools/faces/secured/crispr/crispr.xhtml.

77. Labun, K., et al., CHOPCHOP v3: expanding the CRISPR web toolbox beyond genome editing. Nucleic Acids Res, 2019. 47(W1): p. W171–W174.

78. Pliatsika, V. and I. Rigoutsos, “Off-Spotter”: very fast and exhaustive enumeration of genomic lookalikes for designing CRISPR/Cas guide RNAs. Biol Direct, 2015. 10: p. 4.

79. Bae, S., J. Park, and J.S. Kim, Cas-OFFinder: a fast and versatile algorithm that searches for potential off-target sites of Cas9 RNA-guided endonucleases. Bioinformatics, 2014. 30(10): p. 1473–5.

80. Hwang, G.H., J.S. Kim, and S. Bae, Web-Based CRISPR Toolkits: Cas-OFFinder, Cas-Designer, and Cas-Analyzer. Methods Mol Biol, 2021. 2162: p. 23–33.

81. Horlbeck, M.A., et al., Compact and highly active next-generation libraries for CRISPR-mediated gene repression and activation. Elife, 2016. 5.

82. Satpathy, A.T., et al., Massively parallel single-cell chromatin landscapes of human immune cell development and intratumoral T cell exhaustion. Nat Biotechnol, 2019. 37(8): p. 925–936.

83. Mimitou, E.P., et al., Multiplexed detection of proteins, transcriptomes, clonotypes and CRISPR perturbations in single cells. Nat Methods, 2019. 16(5): p. 409–412.

84. Stuart, T., et al., Comprehensive Integration of Single-Cell Data. Cell, 2019. 177(7): p. 1888–1902 e21.

85. Tirosh, I., et al., Dissecting the multicellular ecosystem of metastatic melanoma by single-cell RNA-seq. Science, 2016. 352(6282): p. 189–96.

86. Papalexi, E., et al., Characterizing the molecular regulation of inhibitory immune checkpoints with multimodal single-cell screens. Nat Genet, 2021. 53(3): p. 322–331.

87. Finak, G., et al., MAST: a flexible statistical framework for assessing transcriptional changes and characterizing heterogeneity in single-cell RNA sequencing data. Genome Biol, 2015. 16: p. 278.

88. Wu, T., et al., clusterProfiler 4.0: A universal enrichment tool for interpreting omics data. Innovation (Camb), 2021. 2(3): p. 100141.

89. Yu, G., et al., clusterProfiler: an R package for comparing biological themes among gene clusters. OMICS, 2012. 16(5): p. 284–7.

90. Andersson, R., et al., An atlas of active enhancers across human cell types and tissues. Nature, 2014. 507(7493): p. 455–461.

91. Eres, I.E. and Y. Gilad, A TAD Skeptic: Is 3D Genome Topology Conserved? Trends Genet, 2021. 37(3): p. 216–223.

92. Zhang, S., et al., The role of blood CXCL12 level in prognosis of coronary artery disease: A meta-analysis. Front Cardiovasc Med, 2022. 9: p. 938540.

93. Amrute, J.M., et al., Targeting Immune-Fibroblast Crosstalk in Myocardial Infarction and Cardiac Fibrosis. Res Sq, 2023.

94. Kuppe, C., et al., Spatial multi-omic map of human myocardial infarction. Nature, 2022. 608(7924): p. 766–777.

95. Gilbert, L.A., et al., CRISPR-mediated modular RNA-guided regulation of transcription in eukaryotes. Cell, 2013. 154(2): p. 442–51.

96. Zhu, Z., et al., Integration of summary data from GWAS and eQTL studies predicts complex trait gene targets. Nat Genet, 2016. 48(5): p. 481–7.

97. Miyazawa, K. and K. Ito, The Evolving Story in the Genetic Analysis for Heart Failure. Front Cardiovasc Med, 2021. 8: p. 646816.

98. Asazuma-Nakamura, Y., et al., Cx43 contributes to TGF-beta signaling to regulate differentiation of cardiac fibroblasts into myofibroblasts. Exp Cell Res, 2009. 315(7): p. 1190–9.

99. Panek, A.N., et al., Connective tissue growth factor overexpression in cardiomyocytes promotes cardiac hypertrophy and protection against pressure overload. PLoS One, 2009. 4(8): p. e6743.

100. Li, Y., et al., Global genetic analysis in mice unveils central role for cilia in congenital heart disease. Nature, 2015. 521(7553): p. 520–4.

101. Huttemann, M., et al., Mice deleted for heart-type cytochrome c oxidase subunit 7a1 develop dilated cardiomyopathy. Mitochondrion, 2012. 12(2): p. 294–304.

102. Doring, Y., et al., The CXCL12/CXCR4 chemokine ligand/receptor axis in cardiovascular disease. Front Physiol, 2014. 5: p. 212.

103. Li, Y., et al., Cardiac Fibroblast-Specific Activating Transcription Factor 3 Protects Against Heart Failure by Suppressing MAP2K3-p38 Signaling. Circulation, 2017. 135(21): p. 2041–2057.

104. Shi, Y., D.J. Riese, 2nd, and J. Shen, The Role of the CXCL12/CXCR4/CXCR7 Chemokine Axis in Cancer. Front Pharmacol, 2020. 11: p. 574667.

105. Tanaka, Y., et al., Systems analysis of ATF3 in stress response and cancer reveals opposing effects on pro-apoptotic genes in p53 pathway. PLoS One, 2011. 6(10): p. e26848.

106. Collaboration, I.R.G.C.E.R.F., et al., Interleukin-6 receptor pathways in coronary heart disease: a collaborative meta-analysis of 82 studies. Lancet, 2012. 379(9822): p. 1205–13.

107. Gonzalez, G.E., et al., Deletion of interleukin-6 prevents cardiac inflammation, fibrosis and dysfunction without affecting blood pressure in angiotensin II-high salt-induced hypertension. J Hypertens, 2015. 33(1): p. 144–52.

108. Interleukin-6 Receptor Mendelian Randomisation Analysis, C., et al., The interleukin-6 receptor as a target for prevention of coronary heart disease: a mendelian randomisation analysis. Lancet, 2012. 379(9822): p. 1214–24.

109. Saadane, N., L. Alpert, and L.E. Chalifour, Altered molecular response to adrenoreceptor-induced cardiac hypertrophy in Egr-1-deficient mice. Am J Physiol Heart Circ Physiol, 2000. 278(3): p. H796–805.

110. Liu, Y., et al., Snail1 is involved in de novo cardiac fibrosis after myocardial infarction in mice. Acta Biochim Biophys Sin (Shanghai), 2012. 44(11): p. 902–10.

111. Gamazon, E.R., et al., Using an atlas of gene regulation across 44 human tissues to inform complex disease-and trait-associated variation. Nat Genet, 2018. 50(7): p. 956–967.

112. Ichiki, T., et al., Differential expression of the pro-natriuretic peptide convertases corin and furin in experimental heart failure and atrial fibrosis. Am J Physiol Regul Integr Comp Physiol, 2013. 304(2): p. R102–9.

113. Yakala, G.K., et al., FURIN Inhibition Reduces Vascular Remodeling and Atherosclerotic Lesion Progression in Mice. Arterioscler Thromb Vasc Biol, 2019. 39(3): p. 387–401.

114. Vandekerckhove, J., G. Bugaisky, and M. Buckingham, Simultaneous expression of skeletal muscle and heart actin proteins in various striated muscle tissues and cells. A quantitative determination of the two actin isoforms. J Biol Chem, 1986. 261(4): p. 1838–43.

115. Consortium, G.T., et al., Genetic effects on gene expression across human tissues. Nature, 2017. 550(7675): p. 204–213.

116. Robins, C., et al., Genetic control of the human brain proteome. Am J Hum Genet, 2021. 108(3): p. 400–410.

117. Sun, B.B., et al., Genomic atlas of the human plasma proteome. Nature, 2018. 558(7708): p. 73–79.

118. He, B., et al., Genome-wide pQTL analysis of protein expression regulatory networks in the human liver. BMC Biology, 2020. 18(1): p. 97.

119. Buccitelli, C. and M. Selbach, mRNAs, proteins and the emerging principles of gene expression control. Nature Reviews Genetics, 2020. 21(10): p. 630–644.

120. Blevins, W.R., et al., Extensive post-transcriptional buffering of gene expression in the response to severe oxidative stress in baker’s yeast. Scientific Reports, 2019. 9(1): p. 11005.

121. Kusnadi, E.P., et al., Regulation of gene expression via translational buffering. Biochim Biophys Acta Mol Cell Res, 2022. 1869(1): p. 119140.

122. Stunnenberg, H.G. and N.C. Hubner, Genomics meets proteomics: identifying the culprits in disease. Hum Genet, 2014. 133(6): p. 689–700.

123. Molendijk, J. and B.L. Parker, Proteome-wide Systems Genetics to Identify Functional Regulators of Complex Traits. Cell Syst, 2021. 12(1): p. 5–22.

124. Fairfax, B.P., et al., Innate immune activity conditions the effect of regulatory variants upon monocyte gene expression. Science, 2014. 343(6175): p. 1246949.

125. Farh, K.K., et al., Genetic and epigenetic fine mapping of causal autoimmune disease variants. Nature, 2015. 518(7539): p. 337–43.

